# A Latent Space Thermodynamic Model of Cell Differentiation

**DOI:** 10.64898/2026.03.04.709512

**Authors:** Ali Poursina, Shayan Hajhashemi, Ali Saberi, Arsham Mikaeili Namini, Ariel Madrigal, Amin Emad, Hamed S. Najafabadi

**Affiliations:** Department of Human Genetics, McGill University, Montreal, QC, Canada; Victor P. Dahdaleh Institute of Genomic Medicine, Montreal, QC, Canada; Quantitative Life Sciences Program, Montreal, QC, Canada; Mila, Quebec AI Institute, Montreal, QC, Canada; Department of Electrical and Computer Engineering, McGill University, Montreal, Canada; Rosalind and Morris Goodman Cancer Institute, Montreal, QC, Canada; McGill Centre for RNA Sciences, McGill University, Montreal, Canada

## Abstract

Inferring the governing dynamics of differentiation that capture cell state evolution remains a central challenge in single-cell biology. We present Single-Cell Latent Space Dynamics (scLSD), a thermodynamics-inspired framework that models cell differentiation as evolution on a learned Waddington landscape in latent space. scLSD jointly infers a low-dimensional cell state, a differentiable potential function governing developmental flow, and a local entropy term that associates with cellular plasticity. Using a neural ordinary differential equation, scLSD reconstructs continuous differentiation trajectories from time-ordered single-cell data. Across diverse developmental systems, scLSD accurately recovers lineage hierarchies, predicts fate commitment for unseen cell types, and outperforms existing trajectory inference approaches in directional and cell fate prediction accuracy. In silico gene perturbations reveal how individual regulators reshape the landscape, and the proposed entropy measure is informative of plasticity in development and cancer.

## Introduction

Cell differentiation is a dynamical process by which gene regulatory networks, epigenetic modifications, and extracellular signals drive transitions from pluripotent or multipotent states to specialized fates, often conceptualized as attractor states within a multidimensional epigenetic landscape^1,2^. A fundamental question in developmental biology is how a cell’s current state determines its future fate. Formally, this problem can be translated to inferring a dynamical law that captures the likely dynamics of cell states along the course of differentiation.

The central computational challenge in understanding cell differentiation lies in inferring this governing law of dynamics^3^, a mathematical framework that can predict the trajectory of cells from their current state to their eventual differentiated fate. Such a governing law should capture the dynamics of cells throughout the entire differentiation process, from early pluripotent stages to fully specialized cell types. Interpreting differentiation as a dynamical system provides a natural mathematical foundation where the governing law corresponds to the equation of motion that describes how cellular states change over time. However, due to the inherent complexity of biological differentiation systems, exact analytical parameterization of these governing equations remains beyond current mathematical understanding, necessitating computational inference approaches.

Comprehensively modeling this process as a computational problem requires integration of heterogeneous data modalities, such as DNA methylation, histone marks, and post-transcriptional/translational regulation, each with distinct biophysical characteristics, leading to a combinatorial explosion of degrees of freedom. An alternative, pragmatic simplification that is often employed^4-7^ is to focus exclusively on transcriptomic dynamics, treating transcript expression levels as proxies for cellular state. This approach is particularly well-suited for single-cell RNA sequencing data. Single-cell transcriptomics has transformed our ability to study differentiation by providing high-resolution snapshots of gene expression in individual cells across developmental time^8^. These profiles reveal a snapshot of the heterogeneity spectrum of cellular states, capturing not only distinct early and late differentiation stages but also the intermediate transitional states that collectively represent the underlying dynamical system.

Despite these advances, a systematic procedure for learning a generalizable mapping from cell state to its dynamics is still lacking. Existing approaches address complementary aspects of this problem and can be broadly divided into methods that infer cell-level progression or local dynamics, and methods that infer the evolution of cell-state distributions across samples. The former include pseudo-temporal ordering methods^9-13^, which leverage cell-to-cell variation to arrange cells along a continuum, reflecting progression through assigning the dynamical variable “pseudotime” to each cell, but provide limited insights into the underlying dynamical processes. RNA velocity methods^4,5,14-18^ provide a more direct estimate of local cell-state dynamics by distinguishing unspliced from spliced transcripts to infer the instantaneous direction of transcriptomic change. However, the first-order kinetic approximation used in these methods, which assumes that each gene’s splicing kinetics follow a system of linear ordinary differential equations (ODEs)^4,14,16,18^, cannot capture hallmark features of nonlinear systems such as multistability^19-21^ and metastability^22,23^. Moreover, the ODEs utilized in RNA velocity models are decoupled across genes^24^; thus, they still do not provide a generalizable map between cell state and cell dynamics beyond simple interpolation within each dataset. A complementary class of approaches uses optimal transport^6,7,25-27^ to characterize population-level dynamics by coupling cell-state distributions observed at different biological times. Depending on the formulation, these methods infer transport plans between discrete samples^6,28-30^ or continuous vector fields^26,31-33^ whose induced dynamics reproduce the observed distributional evolution. Importantly, however, population-level evolution does not uniquely determine the underlying cell-level dynamics^34^: in general, multiple state-dependent dynamical processes can generate the same sequence of cell-state distributions. Thus, although optimal transport provides a powerful description of how populations evolve across time, additional assumptions are required to identify a generalizable mapping from individual cell state to its dynamics. Together, these limitations motivate models that directly learn such a mapping while retaining sufficient flexibility to capture the nonlinear and high-dimensional structure of differentiation.

Here, we introduce a thermodynamics-inspired conceptual framework for studying differentiation dynamics by representing Waddington’s epigenetic landscape^35^ as a learned potential in a latent cell-state space, a reduced-dimensional representation of gene expression learned from data. Our formulation combines two biological inductive biases: long-term lineage progression can be approximated by potential-guided dynamics, and cell-fate decisions involve intrinsic stochasticity^36^. The potential defines the direction of mean cell-state progression, while a probabilistic differentiation-state representation captures developmental variability. We define developmental entropy as the conditional differential entropy of the differentiation state given the cell state and evaluate it as a surrogate for cellular plasticity (the connection between this developmental entropy and thermodynamic entropy depends on additional assumptions described in Methods).

To rigorously evaluate our thermodynamic-inspired framework for differentiation, we developed a unified computational pipeline called Single-Cell Latent Space Dynamics (scLSD). scLSD jointly infers three key quantities from temporally ordered single-cell gene expression data: a latent cell state, providing a compact low-dimensional representation of each cell’s expression profile, a potential function, modeling the Waddington epigenetic landscape that governs lineage progression, and an entropy term associated with developmental variability and cellular plasticity. We first demonstrate that scLSD provides a unified and interpretable framework for modeling single-cell differentiation dynamics across diverse biological systems. By applying scLSD to a range of benchmark datasets, we show how its core output, Waddington potential, captures key features of lineage specification. We further illustrate how scLSD enables quantitative and qualitative evaluation of cell state trajectories, reveals dynamical properties, and outperforms existing approaches in reconstructing biologically meaningful differentiation directions.

Next, we demonstrate that scLSD accurately generalizes to previously unseen cell types by systematically withholding specific populations during training and predicting their fate distributions and differentiation dynamics at inference. Furthermore, to probe the resolution of our approach, we demonstrate the extent of information encoded by single-gene modulations in the Waddington landscape, enabling systematic in silico gene perturbations. We show that scLSD not only models developmental trajectories but also predicts how targeted gene knockouts influence cell fate decisions and lineage specification. Finally, we dissect the predicted entropy in various datasets to quantify plasticity in developmental and cancer contexts, confirming that regions of elevated developmental entropy coincide with the notion of cellular plasticity. These studies not only validate key predictions of our model but also highlight its potential to uncover mechanistic insights into gene-driven landscape remodeling and associations between inferred entropy and cell-fate variability.

## Results

### scLSD models the dynamics of cell states in a Waddington landscape

We represent Waddington’s epigenetic landscape as a potential energy in a cell state space. This transforms the problem of tracking transcriptomic dynamics into tracking the evolution of cell states within a compact, continuous latent space. Our framework models the temporal evolution of “cell states” (embeddings of single cells obtained from their transcriptomic profile) under the negative gradient of the Waddington potential with a local Gaussian noise:

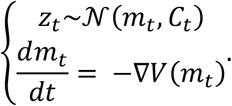

This equation captures the dynamics of the *cell state z* through time *t*, as its mean “rolls” down the *Waddington potential V* with force −∇*V*(*z*) that drives differentiation toward lineage-specific basins. Covariance term *C*_*t*_ represents stochastic fluctuations, stemming from gene expression variability or microenvironmental cues. scLSD infers *C*_*t*_ using a variational autoencoder and combines the mean dynamics with locally inferred uncertainty (see **Methods** for details). In our model, we further compress the cell states into low-dimensional embeddings that represent the *differentiation state*, which captures developmental trajectories, lineage bifurcations and positional relationships within the differentiation process. This allows us to define the *developmental entropy* as the conditional Shannon entropy^37^ of the differentiation state given the cell state.

scLSD implements this framework by jointly inferring cell states, differentiation states, and the Waddington potential from time-ordered single-cell expression data (**Fig. 1A-B**). The value of Waddington potential for each cell can be further used to infer a posterior pseudotime. Both cell states and differentiation states are modeled as Gaussian latent variables. Dedicated multilayer perceptron (MLP) encoders embed each high-dimensional expression profile into a cell state *z* and further compress *z* into a two-dimensional differentiation state, while MLP decoders reconstruct back to ensure information preservation. This encoder-decoder structure learns a nonlinear embedding of the differentiation landscape, enabling a compact yet faithful representation of the dominant axes of developmental variability. The Waddington potential is parameterized by a lower bounded MLP, ensuring landscape stability. Its negative gradient defines a neural ordinary differential equation (neural ODE), which governs the temporal evolution of the inferred cell states and reconstructs continuous differentiation trajectories^38^. The scLSD’s encoders, the neural ODE, and the decoders are jointly trained via stochastic variational inference (SVI) ^39-41^ on time-ordered trajectories derived from snapshot single-cell transcriptomic data. These random walks are probabilistically sampled according to a transition probability matrix constructed from a pseudo-temporal ordering prior (**Fig. 1C-D, Methods**).

**Figure 1.**
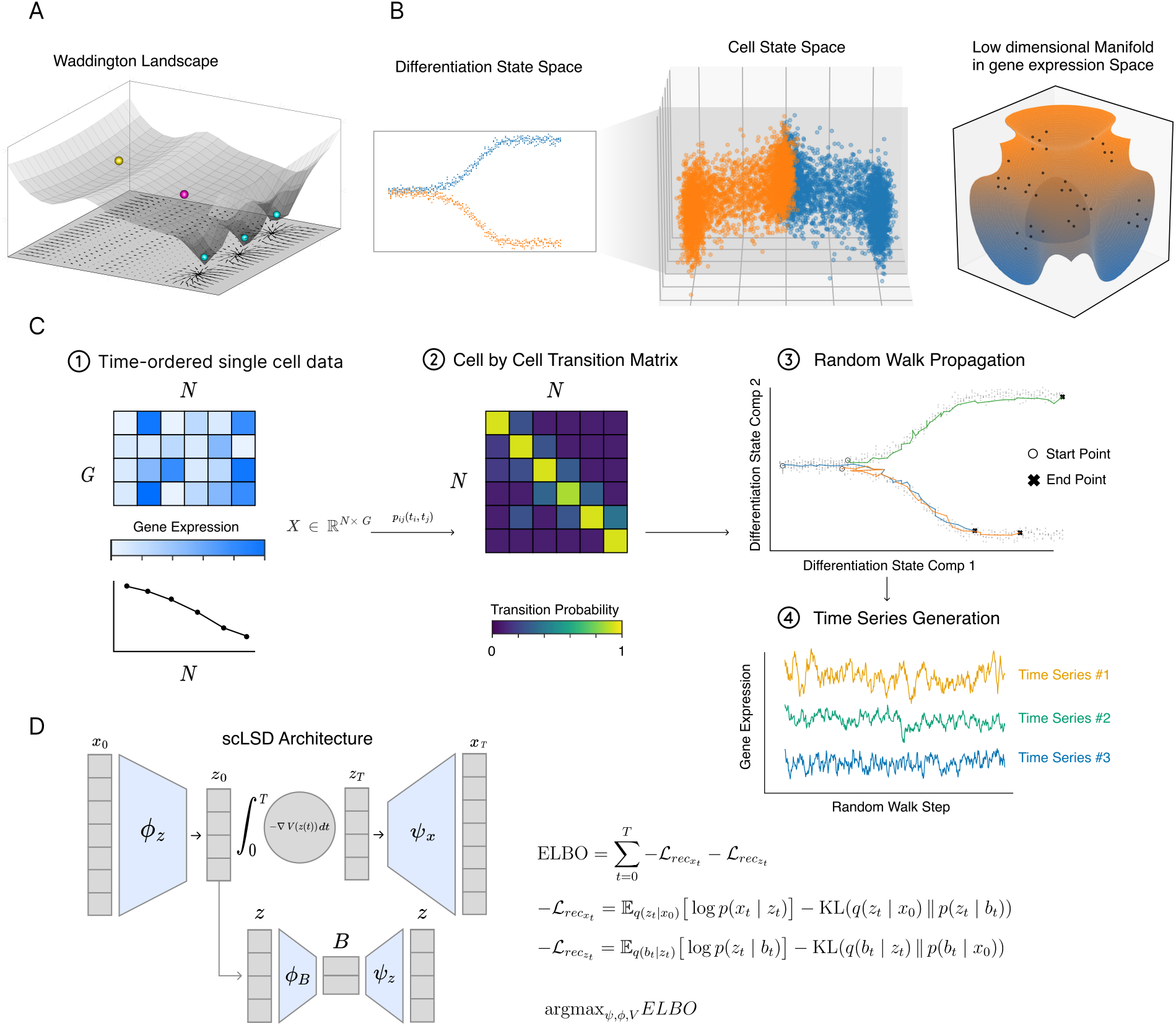
Latent Space Dynamics: a thermodynamic framework for modeling cell differentiation. **(A)** Schematic illustration of the Waddington epigenetic landscape interpreted as a potential energy surface. An initial cell state (yellow) escapes a shallow metastable basin, crosses an energy barrier, and transitions through an intermediate plastic state (pink) before committing to one of several terminal fates residing in deep potential minima (light blue). **(B)** Relationship between observed gene expression and latent representations used by scLSD. Left: the differentiation state, a low-dimensional representation that orders cells along developmental progression. Middle: the cell state space, a higher-dimensional latent representation that captures the full cellular state. Right: the high-dimensional gene expression space, where observed single-cell profiles lie on a low-dimensional manifold that is parameterized by the cell state. **(C)** Construction of synthetic temporal trajectories from time-ordered single-cell data. Transition probabilities between cells are inferred from their temporal proximity and used to generate random walks on a k-nearest-neighbor graph, producing synthetic time series that approximate continuous differentiation dynamics. **(D)** Overview of the scLSD model architecture and training objective. Gene expression profiles are first encoded into latent cell states by the cell state encoder *ϕ*_*z*_. The gene expression profile at the initial step of each random walk, *x*_0_, is mapped to an initial latent cell state *z*_0_ = *ϕ*_*z*_(*x*_0_), which is evolved in continuous time using a neural ODE following the gradient flow of a learned Waddington potential, 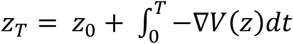, and then is decoded by *Ψ*_*x*_ to reconstruct gene expression profile at step *T, x*_*T*_, yielding the gene expression reconstruction loss 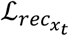. In parallel, latent cell states are mapped to a two-dimensional differentiation state *B* by the differentiation state encoder *ϕ*_*B*_ and decoded back to latent space via *Ψ*_*z*_, inducing a cell state reconstruction loss 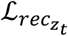. Together, these reconstruction terms define the evidence lower bound (ELBO), which is maximized through variational inference to jointly learn latent states, differentiation states, and the Waddington potential.

### scLSD reconstructs developmental dynamics

scLSD infers the Waddington potential in the high-dimensional cell state space, capturing both global and local developmental dynamics; when combined with a lower-dimensional embedding (e.g., the differentiation state), it yields a compact representation of the developmental landscape. To evaluate the extent to which scLSD’s outputs align with known biology, we analyzed several widely studied single-cell differentiation datasets representing endocrinogenesis, hematopoiesis, erythroid gastrulation, and dentate gyrus development^11,30,42,43^. These datasets encompass a range of differentiation scenarios, from linear trajectories with clear start and end points, to bifurcating and complex multilineage hierarchies with well-established biological relationships among cell types (an example is shown in **Fig. 2A**).

**Figure 2.**
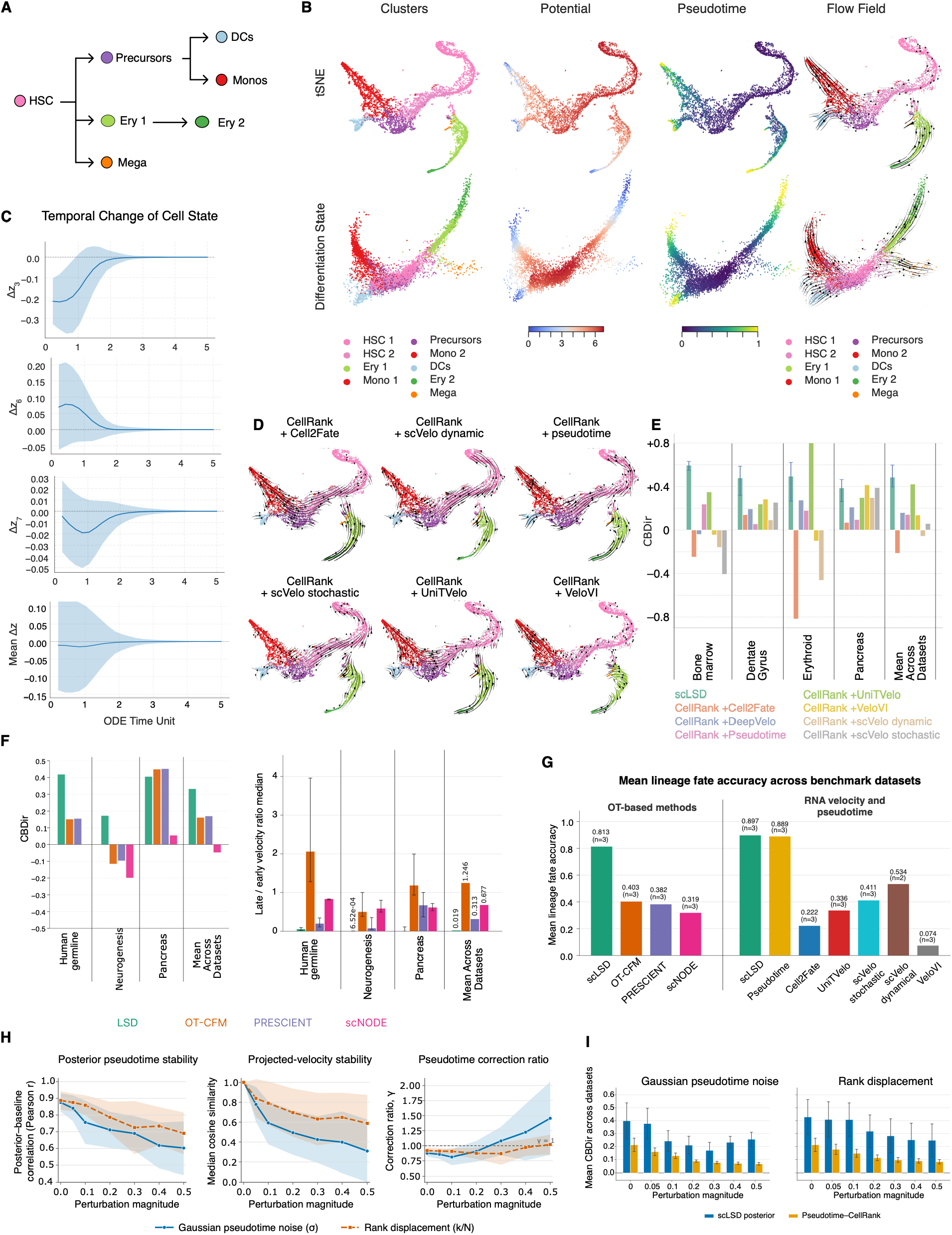
scLSD-inferred developmental landscapes, benchmarking, and sensitivity to pseudotime perturbations. **(A)** Schematic of differentiation tree in bone marrow dataset. **(B)** The bone marrow dataset. Each column shows, from left to right: cell-type clusters, Waddington potential, pseudotime, and developmental flow field. The upper row corresponds to the t-SNE embedding, while the lower row shows the scLSD-inferred differentiation state. **(C)** Temporal evolution of changes in inferred cell state components and mean of all components during trajectory propagation under the learned Waddington potential. The x-axis shows Continuous-time parameter of the neural ODE. The y-axis shows changes in coordinates of components of cell state. The shaded area represents the standard error of the mean. **(D)** Comparison of differentiation trajectories inferred by CellRank using six distinct input models on the human bone marrow dataset. Each panel shows the flow field reconstructed from CellRank transition matrices derived from Cell2Fate, scVelo (Dynamic and Stochastic), Pseudotime, UniTVelo, and VeloVI. **(E)** Bar plot summarizing CBDir scores for different methods (legend, bottom) across four benchmark single-cell differentiation datasets, as well as the mean performance across all datasets. Methods compared include scLSD as well as CellRank initialized with UniTVelo, Pseudotime, VeloVI, scVelo stochastic, DeepVelo, scVelo dynamic, and Cell2Fate. scLSD scores are reported across a set of hyperparameters with an error bar indicating 10^th^ and 90^th^ percentiles of CBDir scores for different configurations. **(F)** Comparison of scLSD, OT-CFM, PRESCIENT, and scNODE across human germline, neurogenesis, and pancreas datasets. Left: CBDir for each dataset and the mean across datasets. Right: median ratio of late to early velocity norms along trajectories initialized from cells with prior pseudotime below 0.5. Late and early velocities are averaged over the final and initial 10% of each trajectory, respectively. Error bars show the interquartile range across trajectories; the final group shows the mean of dataset medians. scLSD median ratios are below one in all three datasets. **(G)** Mean lineage fate accuracy for two benchmark groups. Left: scLSD and OT-based methods on human germline, neurogenesis, and pancreas, using accuracy macro-averaged across source lineages. Right: scLSD, RNA velocity, and pseudotime methods on bone marrow, dentate gyrus, and erythroid, using the mean probability assigned to lineage-compatible fates across eligible cells. Bars show unweighted means of available dataset scores; annotations give the mean and number of datasets (n). scLSD appears separately in each group, with mean scores of 0.813 and 0.897, respectively. **(H)** Sensitivity to Gaussian pseudotime noise (standard deviation σ) and rank displacement (relative magnitude 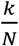) across bone marrow, dentate gyrus, erythroid, and pancreas. Left: Pearson correlation between posterior and original input pseudotime. Middle: median cell-wise cosine similarity between projected velocities from perturbed and unperturbed models. Right: correction ratio γ, defined as the posterior–original pseudotime correlation divided by the perturbed–original pseudotime correlation. Values are averaged over three seeds within each dataset, then across datasets; shaded bands show one standard deviation across datasets. Correlation and velocity similarity generally decrease with perturbation magnitude, while γ rises above one at higher Gaussian noise levels. **(I)** Mean CBDir from scLSD posterior transitions and CellRank transitions derived from the perturbed input pseudotime, shown separately for Gaussian noise and rank displacement. Bars show means across the four datasets in H after averaging across seeds; error bars represent the standard error across datasets. scLSD posterior CBDir is higher at each displayed perturbation level.

When applied to human bone marrow, mouse erythroid gastrulation, and pancreas development datasets (**Fig. 2B** and **Supplementary Fig. 1**), the inferred Waddington potential for each cell showed a consistent decrease along the course of differentiation (**Fig. 2B** and **Supplementary Fig. 1**, second column), in line with the expectation that developmental potential is progressively lost as cells mature. Similarly, the two-dimensional differentiation state embeddings inferred by scLSD provided a compact map of lineage structure consistent with the established biological relationships (**Fig. 2B** and **Supplementary Fig. 1**, lower row). For example, the terminal points of these embeddings correspond to later pseudotime points of fully differentiated states (**Fig. 2B** and **Supplementary Fig. 1**, third column).

scLSD’s primary outputs (differentiation and cell states) can be used to derive secondary representations of cellular behavior. For example, a Boltzmann distribution can be used to calculate the transition probabilities between cells from their Waddington potentials (**Methods**), enabling us to map the directionality of differentiation on data manifold. Across the above datasets, this two-dimensional visualization of resulting streamlines showed qualitative agreement with established biological hierarchies of the cells. In the human bone marrow dataset, for example, the developmental flow field projected on the differentiation state space (**Fig. 2B**, fourth column) revealed four trajectories starting from hematopoietic stem cells (HSC 1) and branching into dendritic cell, monocyte, erythroid, and megakaryocyte fates, consistent with the hematopoiesis differentiation dynamics. Similar lineage structures were observed across other datasets (**Supplementary Fig. 1**). We note that both the Waddington potential and the neural ODE dynamics are defined in the *D*-dimensional cell-state space. The two-dimensional flow fields shown in Fig. 2B are, thus, projections of the cell-to-cell transition dynamics onto the differentiation-state or other low dimensional embedding and should therefore be interpreted as visualizations of the higher-dimensional dynamics, rather than as gradients of a two-dimensional potential.

We next assessed whether the trajectories dictated by the inferred landscape converge to stable attractor states. We tracked changes in the components of the inferred cell state as trajectories were propagated forward in time (**Fig. 2C**). These changes consistently diminished to zero for all cells, demonstrating that every cell reaches a stable attractor state after an initial phase of nonlinear change. In other words, the model predicted robust and stable fate commitment at the end of differentiation trajectory. We further examined whether this behavior depended on constraining the Waddington potential to be lower bounded. Across benchmark datasets, removing the non-negativity constraint resulted in persistent long-term latent dynamics, whereas the constrained potential produced a progressive decay of the latent velocity toward zero **(Supplementary Fig. 2)**. These results support the lower-bound Waddington potential as an important inductive bias for obtaining stable long-term differentiation dynamics.

To systematically benchmark scLSD, we considered two complementary classes of dynamical inference methods. First, we compared scLSD with the initial pseudotime used by scLSD as a temporal-ordering baseline, as well as RNA velocity methods, which estimate local transcriptomic dynamics. These approaches differ fundamentally from scLSD: RNA velocity estimates the local direction of change in gene-expression space, whereas scLSD jointly learns a latent cell-state representation and a continuous state-dependent dynamical field by relying on time ordered expression profiles; pseudotime, in contrast, provides only a scalar ordering of observed cells along an axis of variation among cells. To place these heterogeneous outputs on a common footing, we used CellRank^44^ to convert RNA velocity estimates and pseudotime into cell-to-cell transition matrices, which were subsequently projected onto a common embedding to obtain developmental flow fields and calculate CBDir (cross-boundary direction correctness) scores, a performance metric ranging from –1 to 1 and representing the degree to which the projections of trajectories are consistent with the expected relationships among cell types (**Methods**). CellRank therefore serves here as a common evaluation interface rather than as an independent dynamical model.

As a complementary benchmark, we compared scLSD with continuous optimal transport methods. Similar to scLSD, these approaches parameterize continuous dynamics through a differential equation and thereby infer a velocity field over a continuous state space. Their inference principle, however, is distinct: continuous optimal transport methods learn dynamics that transport cell-state distributions between consecutive temporal snapshots while minimizing a transport objective, whereas scLSD learns the velocity field by minimizing the loss over cell-level trajectories. Thus, the two benchmark classes together test scLSD against both local cell-level and population-level dynamical estimates.

Qualitatively, we observed that trajectories inferred by CellRank often diverged from the known biological direction of differentiation across several datasets and input modalities. For instance, in the bone marrow system, CellRank trajectories failed to recapitulate the established progression from HSC 1 toward myeloid and erythroid lineages for all input methods except UniTVelo^15^ (**Fig. 2D** and **Supplementary Fig. 3**). To quantify performance, we compared scLSD to the transition matrix obtained by CellRank from a range of widely used methods, with CBDir score as the performance metric^15^. Across multiple datasets and initiating methods used as input to CellRank, we found that scLSD consistently achieved higher average CBDir scores compared to CellRank, even when CellRank was provided with the same initial time ordering used for initializing the scLSD model (**Fig. 2E** and **Supplementary Fig. 4**). This shows that scLSD captures and refines dynamical information beyond the information embedded in the initial pseudotime. Notably, this pattern held in cases where many other methods failed to produce biologically meaningful directionalities and sometimes scored negative CBDir values, predicting the inverse of the expected differentiation directions. For scLSD, CBDir was evaluated across a range of hyperparameters; **Fig. 2E** reports the mean score, with error bars indicating the 10^th^ and 90^th^ percentiles, demonstrating that scLSD’s inferred dynamics remain robust to hyperparameter variation.

We next benchmarked scLSD against three continuous dynamical models, OT-CFM^32^, PRESCIENT^33^, and scNODE^45^, across pancreas^30^, human neurogenesis^46^, and human germline differentiation^47^ datasets (**Fig. 2F, Supplementary Fig. 5,6**). scLSD almost consistently achieved higher CBDir scores across the three datasets, indicating better agreement between its inferred dynamics and the expected lineage directions (**Fig. 2F**, left). We further asked whether the inferred flow fields vanish as cells approach terminal attractor states. For each method, we compared the velocity magnitude of cells at late versus early pseudotime and quantified the median late-to-early velocity ratio (**Methods)**. This ratio approached zero only for scLSD, with a mean ratio of 0.019 across datasets, whereas the next-lowest value, obtained by PRESCIENT, was 0.313, approximately 16-fold larger (**Fig. 2F**, right). Notably, PRESCIENT, the closest comparator by this measure, is also the only one among these three methods that explicitly formulates differentiation through a Waddington-like potential, consistent with the notion that imposing potential-driven dynamics favors stable long-term behavior.

Because CBDir measures local directional consistency at cell-type boundaries and depends on the embedding used for projection, we additionally evaluated lineage fate prediction accuracy as a complementary measure of global dynamical consistency (**Methods**). For RNA velocity methods and prior pseudotime, fate predictions were obtained from CellRank transition matrices using CellRank’s fate-mapping procedure, whereas for scLSD and the continuous dynamical models, cells were propagated under the inferred vector field to terminal lineages. scLSD achieved the highest mean fate prediction accuracy in both benchmark categories, reaching 0.813 across the continuous-model datasets and 0.897 across the RNA velocity/pseudotime datasets (**Fig. 2G, Supplementary Fig. 7**). Among alternative methods, only CellRank initialized with the same pseudotime used for scLSD achieved comparable performance (0.889), indicating that scLSD preserves the global lineage structure encoded by the initial temporal ordering while learning continuous dynamics. The lower fate accuracies of several RNA velocity methods should, however, be interpreted cautiously, as they reflect both the underlying velocity estimates and subsequent CellRank fate-mapping steps, including macrostate identification.

We next assessed the robustness of scLSD to noise in its input pseudotime. We perturbed the input pseudotime using either additive Gaussian noise or rank displacement at increasing strengths and compared the resulting scLSD models with the baseline pseudotime and model (**Methods**). Although posterior pseudotime agreement with the baseline decreased with perturbation strength, indicating dependence of scLSD outputs on prior pseudotime, substantial correlation was retained even under strong perturbations (**Fig. 2H** left, **Supplementary Fig. 8A**). Likewise, the inferred vector fields remained positively aligned with the baseline model, as measured by their median cosine similarity (**Fig. 2H** mid, **Supplementary Fig. 8B**). To quantify whether scLSD could recover information degraded in the input, we defined a correction ratio, *γ* = r(τ_baseline_, τ_*posterior*_)/r(τ_baseline_, *τ*_perturbed_), where *r* indicates the Pearson correlation and *τ* indicates the pseudotime. At stronger perturbations, *γ* approached or exceeded one (**Fig. 2H** right, **Supplementary Fig. 8C**), indicating that scLSD can partially rescue the dynamical structure.

We further compared the CBDir scores of scLSD models trained on perturbed pseudotime with those obtained directly from the same perturbed pseudotime inputs. Across perturbation strengths, the scLSD-derived vector fields consistently achieved higher CBDir scores, despite being initialized from the identical noisy temporal ordering (**Fig. 2I**).

### scLSD recovers fate and maturation of unseen lineages

scLSD learns a continuous flow field over the cell state space, while variational inference represents each cell by a distribution over its latent state. Together, these components define an ensemble of trajectories with probabilistic initial conditions; in other words, conditional on a sampled initial cell state, scLSD can propagate the cell through time (**Methods**), allowing it to not only identify developmental trajectories (as shown in the previous section), but also extrapolate across time. We therefore used the scLSD-inferred flow field to examine long-term fate behavior by propagating cells until they approached terminal states, thereby assigning predicted cell fates. As shown in **Fig. 3A-B**, across different biological systems, scLSD was able to infer terminal fates that corresponded to biologically meaningful endpoints, with cluster-level fate distributions consistent with known lineage hierarchies (**Fig. 3A–B, Supplementary Fig. 9**). By sampling each cell’s initial state from its inferred latent distribution and propagating these samples independently, we were also able to obtain empirical cell-specific distributions over terminal fates and quantify uncertainty in fate commitment (**Supplementary Fig. 10**).

**Figure 3.**
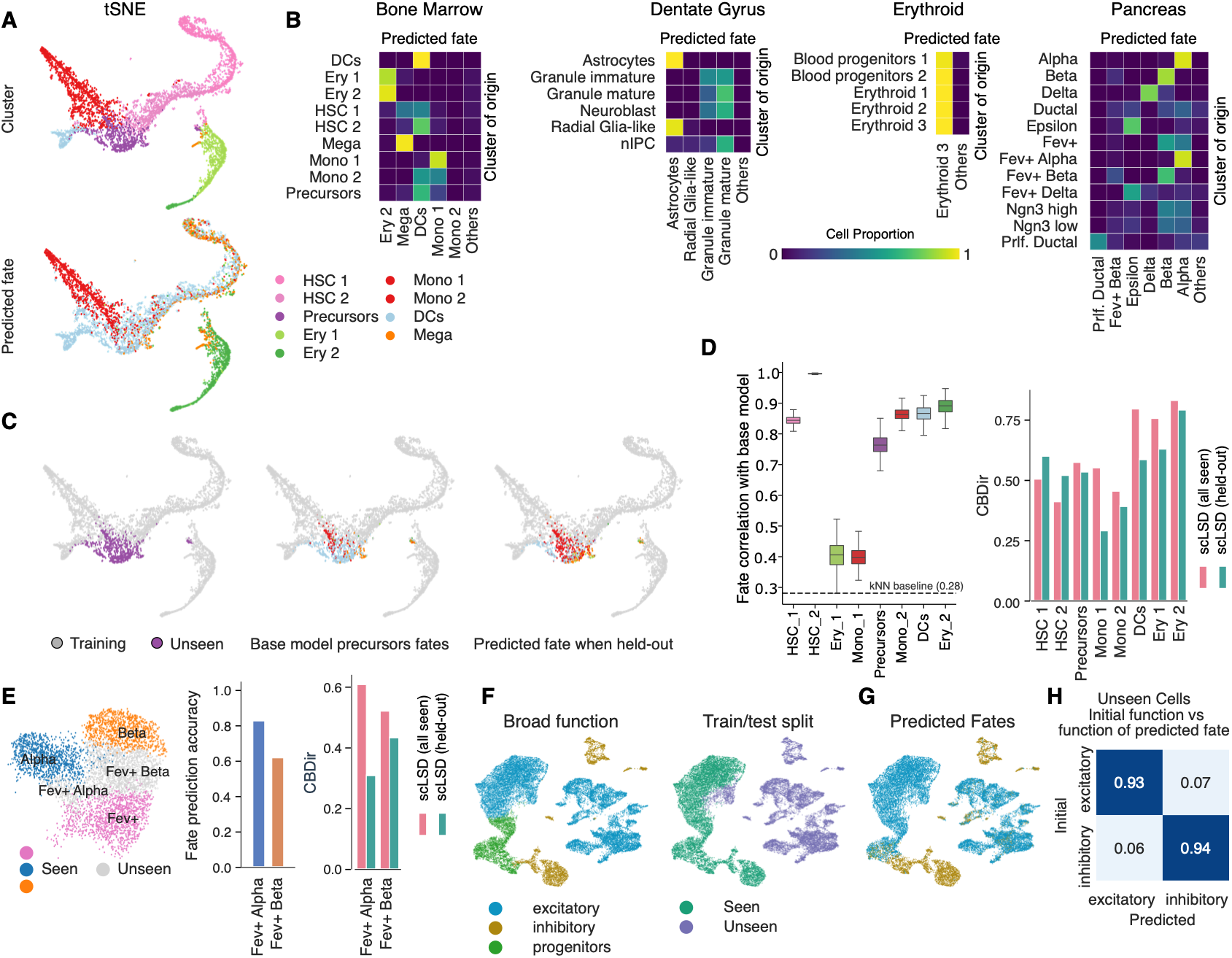
scLSD generalizes developmental dynamics and fate prediction across unseen cell populations and datasets. **(A)** Comparison of cellular clusters and terminal fates in bone marrow differentiation. The top row shows annotated cell clusters, while the bottom row displays inferred terminal fates. **(B)** Contingency heatmaps showing the frequency of predicted cell fates for each cluster across four benchmark datasets. **(C)** Left: Training setup in which only precursor cells (purple) were withheld from training and used for testing. Middle: Fate predictions for the precursor cluster when included in training. Right: Fate predictions for the same precursor cells when excluded from training. **(D)** Left: Correlation between predicted cell fate frequencies when each cluster was included (“seen”) versus excluded (“unseen”) from training, computed using 500 bootstrap replicates. Right: CBDir scores for models trained with and without each cluster. The dashed line indicates the correlation between fate predictions from a 15-nearest-neighbor baseline in PCA space and the full-data scLSD model (r = 0.28). **(E)** Left: Fev+ Alpha and Fev+ Beta cells were jointly excluded from training to evaluate scLSD’s ability to generalize beyond directly observed differentiation branches. Middle: Fate prediction accuracy for unseen cell types, shown separately for Fev+ Alpha and Fev+ Beta clusters when excluded from training. Right: CBDir comparison between models trained with all cell types and those trained with the Fev+ Alpha/Beta clusters withheld. **(F)** Left: UMAP representation of two mouse cortex datasets showing three major functional categories: progenitors, excitatory neurons, and inhibitory neurons. Right: The same UMAP colored by train/test split: training (seen) includes all three categories (progenitors, excitatory, inhibitory), whereas the held-out validation (unseen) dataset contains only fully differentiated excitatory or inhibitory cells. **(G)** UMAP representation of predicted cell fates by scLSD. **(H)** Confusion matrix showing scLSD accuracy for unseen excitatory and inhibitory neurons in predicting their function.

To assess whether the scLSD-inferred mappings between the gene expression profile, the cell state, and the Waddington potential generalize to previously unobserved cell types, we performed a systematic cell type-stratified cross-validation across three developmental single-cell RNA-seq datasets: human bone marrow^11^, pancreatic endocrinogenesis^30^, and mouse cortical development^48^. In each experiment, we excluded one cell type from the training data and trained an scLSD model on the remaining cell types. The held-out cell type was then introduced only at inference time to evaluate the model’s predictions for previously unseen populations (**Fig. 3C** and **Supplementary Fig. 9E)**. We repeated this procedure for each cell type representing various early, intermediate, and terminal developmental stages, and evaluated the model’s predictions by quantifying both the fate distribution and the CBDir metric associated with the velocity embedding. In the bone marrow dataset^11^, fate prediction distributions correlated strongly with those obtained from the full dataset (i.e., without excluding any cell type), with Pearson r > 0.8 for the majority of clusters (**Fig. 3D** left). In comparison, as a geometry-only reference, a 15-nearest-neighbor baseline based on proximity to terminal fate cells showed substantially lower agreement with the full-model scLSD fate assignments (**Methods, Fig. 3D** left). The inferred trajectories in the left-out cell types were also highly consistent with the expected direction, as measured by CBDir scores (mean CBDir 0.55, **Fig. 3D** right). Although in most cases there was a reduction in the CBDir score compared to the scLSD model trained on the full data (mean CBDir 0.61), the inferred trajectories outperformed those of UniTVelo (mean CBDir 0.35), which was the top-performing RNA velocity method on the bone marrow dataset (evaluated in **Fig. 2D-E**).

In a more challenging experiment, we excluded two cell types at the same time from a pancreatic development dataset^30^. Specifically, we simultaneously excluded the Fev+ Alpha and Fev+ Beta populations from the training data, effectively disconnecting the progenitor Fev+ cells from the terminal Alpha and Beta cell types (**Fig. 3E** left). In spite of this, an scLSD model trained on the remaining, disjoint cells was able to predict the expected fate and the trajectory of the left-out cell types: 83% of the Fev+ Alpha cells were correctly predicted to preferentially progress toward the Alpha cell fate and 62% of the Fev+ Beta cells correctly followed a trajectory toward the Beta fate (**Fig. 3E** middle, and **Supplementary Fig. 11A**), with a mean CBDir of 0.37 across the two unseen cell types (compared to the mean CBDir score of 0.57 for a model trained on the full data; **Fig. 3E** right and **Supplementary Fig. 11B**).

Finally, we designed a validation task in which an scLSD model was trained exclusively on cell types from the early stages of mouse cortical neurogenesis^48^ using early progenitor and immature neuronal populations from the same experiment. The model was then tested on a large, independent dataset of fully differentiated neuronal subtypes spanning both excitatory and inhibitory lineages (**Fig. 3F** and **Supplementary Fig. 11C)**. Our aim was to determine whether scLSD could accurately classify these mature, functionally distinct neuronal populations, despite having only encountered their developmental precursors during model training. Interestingly, the scLSD-derived predicted fates showed a seamless connection between these terminal populations and those included in training, in a manner consistent with their broad functions (inhibitory vs. excitatory; **Fig 3G** and **Supplementary Fig. 11D**), despite the fact that there was considerable separation between the profiles of their two respective experiments. scLSD achieved 93% and 94% accuracy for assigning excitatory and inhibitory identity to previously unseen mature neurons, respectively (**Fig. 3H**). Importantly, the pseudotime distribution for the test mature neuronal subtypes was biased toward the maximum pseudotime value of 1, indicating that scLSD correctly recognizes these populations as terminally differentiated (**Supplementary Fig. 11E**, mean pseudotime for unseen types 0.797 ± 0.113).

### scLSD reveals how individual genes influence the Waddington landscape

A biology-aware model of the Waddington landscape should encode information about how individual genes influence differentiation dynamics and cell fate decisions. To test whether the potential landscape learned by scLSD encodes such information, we developed an *in silico* perturbation algorithm that systematically evaluates how targeted gene perturbations influence cellular differentiation dynamics. Specifically, after training the scLSD model, we simulated perturbations by altering gene expression profiles to mimic gene knockouts, encoding these modified profiles as initial conditions in cell state space and propagating their trajectories using the inferred potential landscape (**Methods**). Perturbations were implemented using an iterative procedure: after each propagation step in the latent space, the gene expression profile was modified by resetting the gene of interest’s expression to zero, re-encoded into latent space, and propagated again. The number of iterations in this loop represents the duration of the perturbation, ranging from a single-step transient perturbation to multi-step sustained suppression. We used this framework in two developmental systems: zebrafish axial mesoderm development^49^ and mouse cortical development^48^, both of which are extensively characterized in the literature with well-established gene regulatory programs^49-54^.

We started by exploring the scLSD model trained on the zebrafish axial mesoderm development data^49^. The developmental flow field embedding and cell fate predictions of this model reflected the expected differentiation trajectories from early blastomeres toward the two distinct terminal states (notochord and prechordal plate lineages; **Fig. 4A**), and the inferred differentiation state representation showed a clear bifurcation, marking the commitment to alternative cell fates (**Fig. 4A** bottom row). We initially focused on the *in silico* perturbation of *noto* (notochord homeobox), a gene encoding a homeodomain transcription factor that serves as a master regulator in notochord development across vertebrates^55,56^ (in zebrafish, *noto* is expressed in notochord progenitor cells and is essential for proper axial development). Analysis of the perturbed trajectories revealed that, following *noto* knockout, cells in the early developmental stages are reprogrammed toward prechordal plate, as expected, while terminally differentiated notochord cells demonstrated remarkable robustness to *noto* perturbation (**Fig. 4B**), consistent with the expected irreversibility of differentiation. We also observed a clear relationship between the number of perturbation iterations and perturbation sensitivity: increasing the number of perturbation iterations progressively decreased the proportion of cells differentiating toward notochord fate up to a few iterations, after which the effect plateaued (**Fig. 4B** heatmap and **Supplementary Fig. 12**). Interestingly, even a single-step perturbation was sufficient to observe 51% of the cell fate changes (**Fig. 4B** top). This suggests that our model is able to propagate the effect of transient perturbations without the need to continuously resupply those perturbations beyond a few initial rounds.

**Figure 4.**
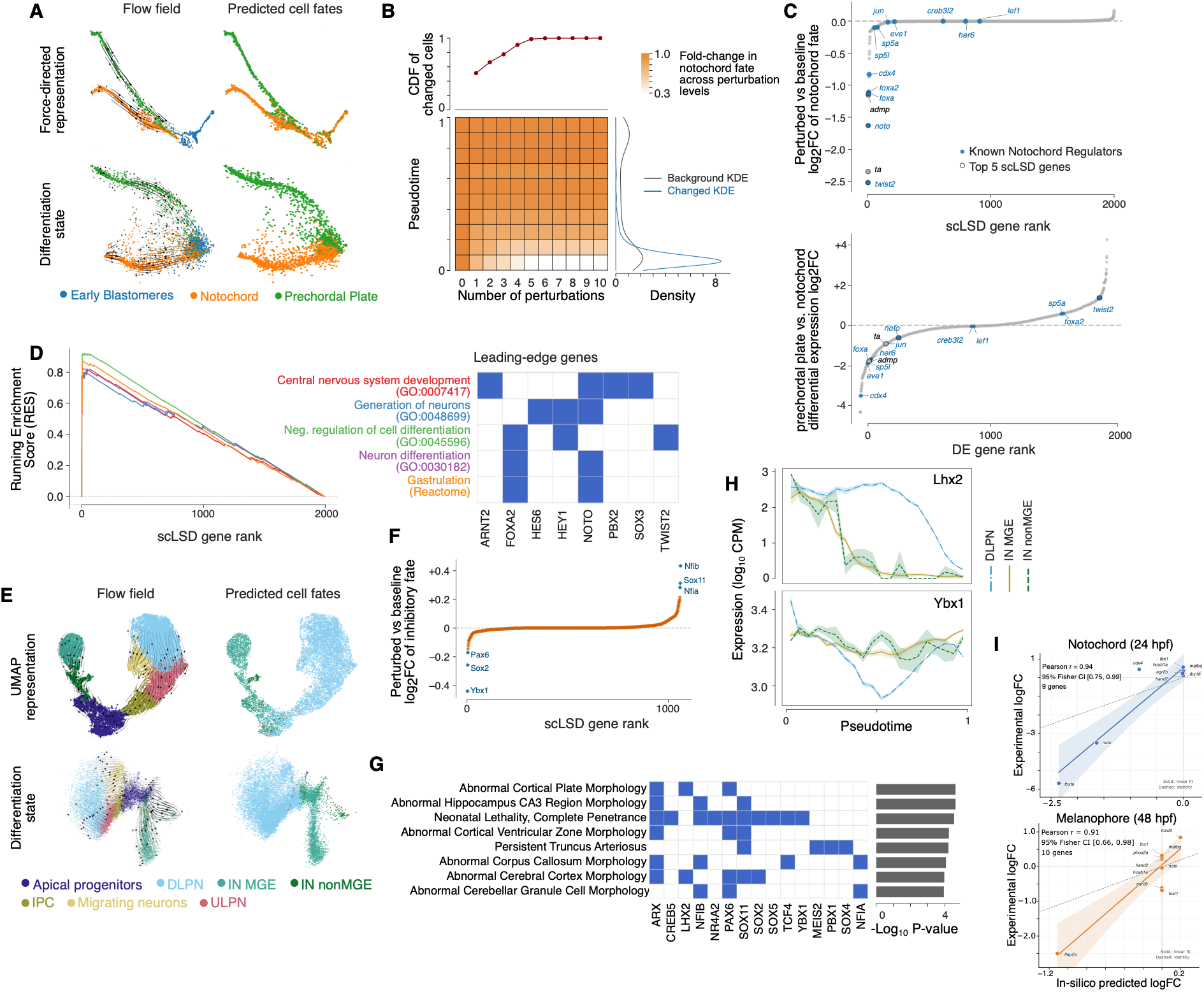
Interrogating the gene-specific structure of the inferred Waddington landscape through in silico perturbations and temporal expression analysis and experimental comparisons. **(A)** Zebrafish axial mesoderm development model. Left: Flow field inferred by scLSD over the force directed graph (top) and differentiation state (bottom). Right: scLSD-predicted cell fates mapped onto both visualizations. Note the division of early blastomeres between notochord and prechordal plate fates. **(B)** Summary of *noto* perturbation results. Top: the cumulative proportion of fate changes (y-axis) occurring at each perturbation iteration (x-axis). Bottom left: Heatmap showing the proportion of fate changes across pseudotime bins (y-axis) and perturbation iterations (x-axis). Bottom right: Distribution of pseudotime for all cells (gray, “Background”) compared to cells whose fates changed following *noto* perturbation (blue). Note the concentration of cells exhibiting fate changes at early pseudotime values. **(C)** Top: Scatter plot showing the results of large-scale perturbation analysis. Each dot represents one in silico gene knockout. The y-axis shows the notochord fate log fold-change (logFC) after knockout of each gene compared to the unperturbed condition. Negative values indicate a reduction in the frequency of cells destined for notochord fate, while positive values indicate an increase in notochord fate frequency after gene perturbation. The x axis shows the scLSD-assigned rank of the gene based on the fate log fold-change. Transcription factors involved in axial mesoderm differentiation based on a previous report^53^ are shown in cyan. Top five genes whose in silico perturbation resulted in largest notochord fate decrease are highlighted with black border. Bottom: Same as above, but with x-axis showing the differential expression analysis gene rank and y-axis showing the log fold change in expression between terminally differentiated prechordal plate and notochord cells (negative values indicate higher expression in notochord cells). Same genes as above are highlighted. **(D)** Gene Set Enrichment Analysis (GSEA)^60^ of genome-wide perturbation results. Top five enriched terms based on GO Biological Processes^62^ and Reactome^63^ are shown. The x-axis represents gene rank (based on scLSD notochord fate logFC), and the y-axis shows the pathway Running Enrichment Score (RES). The leading-edge gene sets for the top enriched pathways are shown on the right. **(E)** UMAP (top) and differentiation state (bottom) representations of the mouse cortex development dataset, annotated by (left) inferred flow field from scLSD and (right) predicted terminal fates. **(F)** Similar to **C**, but with y-axis showing logFC of inhibitory fate after in silico gene perturbation. Negative values indicate a shift toward excitatory lineage commitment after gene knockout, whereas positive values indicate a shift toward inhibitory lineage. **(G)** Gene set overlap analysis of MGI (Mouse Genome Informatics)^71^ phenotypes associated with scLSD top genes from the mouse cortex in silico perturbations. Phenotypes with FDR<0.01 are shown (Fisher’s exact test; p-values shown in the barplot). **(H)** Expression trends of *Lhx2* (top) and *Ybx1* (bottom), stratified by scLSD-predicted cell fate. The shaded area represents the standard error of the mean. **(I)** Comparison of scLSD-predicted fate-abundance log fold-changes with experimental cell-type abundance log fold-changes for notochord at 24 hours post-fertilization (top; 9 genes) and melanophore at 48 hours post-fertilization (bottom; 10 genes). Each point represents a matched gene perturbation. Solid lines show linear fits with 95% confidence bands; dashed lines show the identity relation. Pearson correlations are 0.94 and 0.91, with 95% confidence intervals computed using Fisher’s transformation of [0.75, 0.99] and [0.66, 0.98], respectively.

Encouraged by the model’s ability to predict the effect of *noto* perturbations, we performed a systematic *in silico* knockout screen in early blastomere cells to identify genes with the strongest influence on fate allocation (the screen included all transcription factors). The upper panel of **Fig. 4C** shows the log fold-change in notochord fate proportions after each gene knockout; most genes had near-zero effect on fate allocation, whereas a small subset induced a large reduction in the proportion of notochord-committed cells. We note that the top four genes with the largest effect size, namely *twist2, ta* (also known as *tbxta), noto*, and *admp* all have established roles in notochord development^57-59^, underlining the ability of scLSD to identify master regulators of cell fate. Systematic comparison to a set of experimentally validated transcription factors known to be essential for notochord development^53^ revealed three additional developmental regulators, *foxa, foxa2*, and *cdx4*, among the top eight influential genes identified by scLSD. Thus, 7 out of 8 top scLSD genes have literature or experimental support. In contrast, sorting the genes by differential expression between terminal notochord and prechordal plate cells did not clearly separate these cell fate regulators from other genes, with only one of the top 30 genes with the largest log fold-change in expression overlapping notochord regulators (*cdx4*, **Fig. 4C** bottom). GSEA^60,61^ analysis of gene ontology (GO)^62^ terms and Reactome^63^ pathways, based on the ranked list of genes ordered by their influence on notochord fate, further supported the strong enrichment of developmental genes among scLSD top hits. Specifically, the top five significantly enriched pathways and GO terms included gastrulation, central nervous system development, generation of neurons, neuron differentiation, and negative regulation of cell differentiation (**Fig 4D**).

We next applied this perturbation framework to the mouse cortical development dataset^48^. At baseline, scLSD identified three terminal attractor states corresponding to two inhibitory neuronal subtypes (IN-MGE, inhibitory neurons derived from the medial ganglionic eminence, and IN-non-MGE, inhibitory neurons with non-MGE origins) and one excitatory subtype (DLPN, deep-layer projection neurons) (**Fig. 4E**). Gene knockout simulations in apical progenitor cells (spanning all transcription factors) again showed that only a small subset of genes exerted strong effects on fate allocation, mirroring the zebrafish results (**Fig. 4F**). Among the top-ranked regulators, inhibition of *Ybx1, Sox2*, and *Pax6* was predicted to promote excitatory neuron differentiation, while inhibition of *Nfib, Sox11*, and *Nfia* promoted inhibitory neuron differentiation; all six transcription factors are well-established regulators of central nervous system and cortical development^64-70^, supporting the biological plausibility of scLSD’s inferred gene-fate associations. Gene set overlap analysis using the predicted top 10 excitatory and top 10 inhibitory lineage regulators suggested that perturbation of these genes in mice has been linked to phenotypes linked to brain development (based on Mouse Genome Informatics data^71^), including abnormal cortical plate morphology and abnormal cerebral cortex morphology (Fisher’s exact test, FDR < 0.01, **Fig. 4G)**, demonstrating that this small set of influential TFs collectively impacts a broad spectrum of brain development and neuronal differentiation.

Next, we note the agreement between the temporal expression patterns of these genes and scLSD’s *in silico* perturbation predictions (examples are shown in **Fig. 4H** and **Supplementary Fig. 13**). Importantly, by predicting the (unperturbed) fate of each cell, our framework enables examination of these gene expression patterns based on the fate to which the cells are committed, revealing patterns that are not observable based on analysis of terminally differentiated cells alone. For example, *Lhx2* is expressed at similar levels in all terminal cells across the three major lineages; however, intermediate cells show divergent expression of *Lhx2* depending on their scLSD-inferred cell fate commitment (**Fig. 4H**). Specifically, *Lhx2* remains highly expressed in intermediate cells differentiating into excitatory neurons and only declines in late pseudotimes; in contrast, it declines early in inhibitory neuron lineages. *Ybx1* is another such example, which is expressed abundantly in both progenitor and terminally differentiated cells, and shows temporal down-regulation only in intermediate cells committed to excitatory neuron fate. These temporally resolved transcriptional patterns show that scLSD can capture the coordinated timing of the activity of lineage-specific regulators during differentiation.

Finally, we quantitatively evaluated scLSD’s in silico perturbation predictions against a large-scale experimental zebrafish perturbation atlas^72^ comprising 1,812 individually resolved embryos across 19 developmental time points and 23 genetic perturbations, totaling 3.2 million cells. We first compared scLSD-predicted changes in notochord abundance (same model described above) with experimentally observed cell-type abundance changes at 24 hours post-fertilization (hpf) for perturbations shared between the two analyses. Predicted and experimental log fold-changes showed strong agreement (Pearson r=0.94, 95% CI: 0.75–0.99; **Fig. 4I**, top), including concordant effects for key notochord regulators *tbxta* and *noto*. Of note, scLSD predicted a larger reduction in notochord fate after *tbxta* perturbation compared to *noto*, concordant with experimental observations. To assess whether this agreement generalized beyond the developmental system used for our initial perturbation analyses, we trained a separate scLSD model of zebrafish pigmentation development (using unperturbed cells from ref^72^) and predicted perturbation-induced changes in melanophore abundance at 48 hpf. These predictions again closely matched the experimental measurements (Pearson r=0.91, 95% CI: 0.66–0.98; **Fig. 4I**, bottom), with *foxd3* and *tfap2a* among the perturbations showing the strongest (and opposite) effects in both experimental and scLSD-derived estimates.

### scLSD’s developmental entropy is associated with plasticity during differentiation and cancer progression

scLSD jointly infers the Waddington potential, which defines the direction and stability of differentiation, and the developmental entropy, which quantifies the diversity of differentiation states accessible from a given cell state. To explore the biological interpretation of the developmental entropy, we first examined its evolution along differentiation trajectories in three representative models spanning linear to complex multilineage landscapes with multiple local minima: mouse erythroid gastrulation^43^, endocrinogenesis (pancreas)^30^, and hematopoiesis (bone marrow)^11^. Across all three systems, entropy followed a consistent pattern (**Fig. 5A**): it was elevated in early progenitors and progressively decreased toward terminally differentiated states. This monotonic decline, which is also quantitatively reflected in the negative correlation between entropy and pseudotime in all three systems (**Fig. 5B, Supplementary Fig. 14**), mirrors the expected loss of developmental plasticity, wherein progenitor cells possess broad lineage potential, while mature cells exhibit reduced flexibility and stable fate commitment. Thus, scLSD-inferred entropy may reflect developmental plasticity.

**Figure 5.**
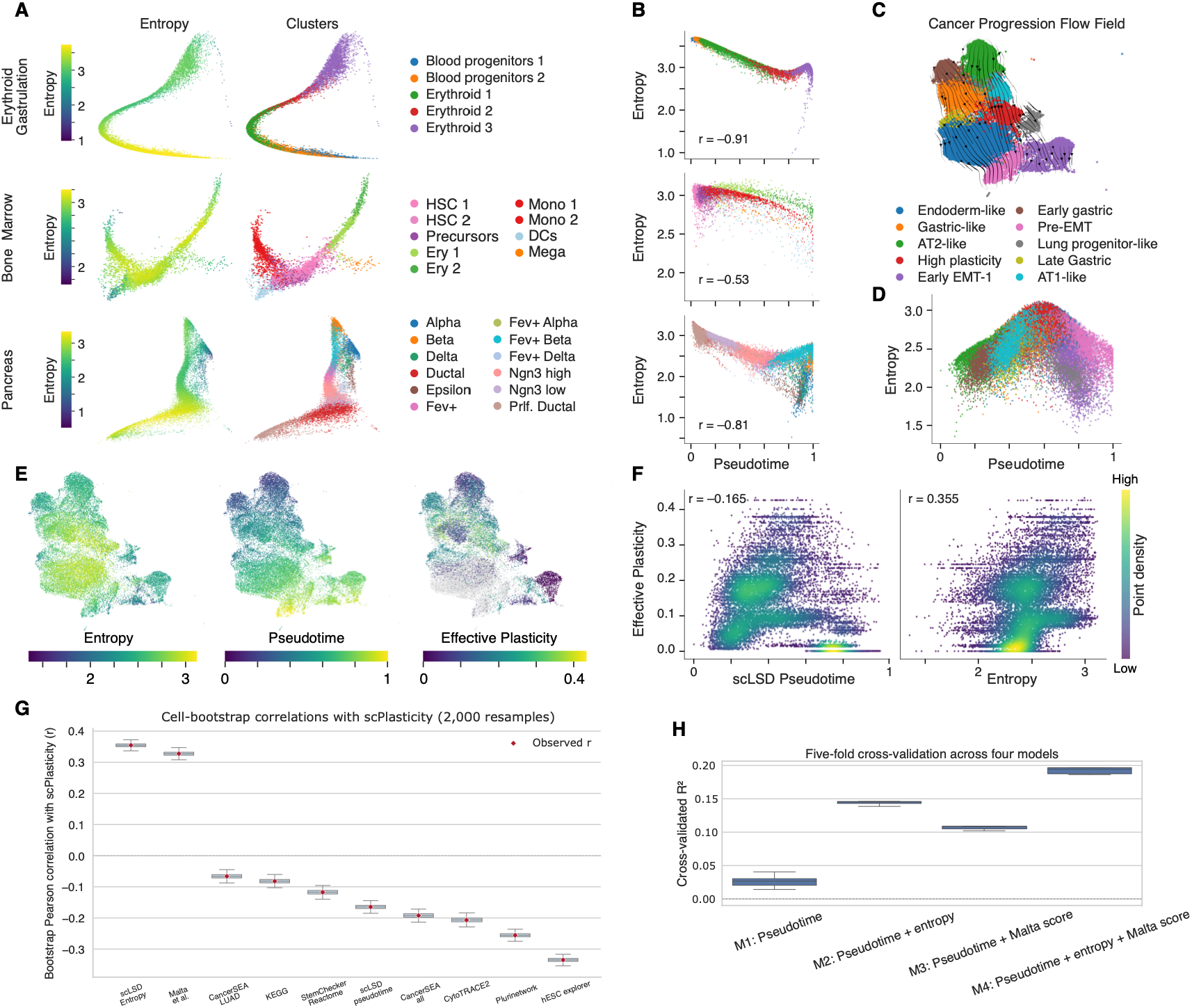
Entropy and local plasticity association across normal development and cancer progression. **(A)** Two-dimensional differentiation representations inferred by scLSD for developmental datasets. The first column shows the corresponding developmental entropy values for individual cells, while the second column visualizes the corresponding annotated cell-type clusters. **(B)** Scatterplots showing the relationship between developmental entropy and scLSD-inferred pseudotime in each dataset. **(C)** Flow field projections of scLSD-inferred velocity vectors overlaid on the UMAP embedding of single-cell transcriptomic data from a genetically engineered mouse model of lung adenocarcinoma. **(D)** Scatterplot showing entropy vs scLSD-inferred pseudotime for cells from the lung adenocarcinoma dataset. **(E)** UMAP visualization of single-cell transcriptomes from the lung adenocarcinoma dataset. First and second columns: scLSD-inferred developmental entropy and pseudotime, respectively. Third column: Effective plasticity values from Yang et al.^73^. **(F)** Scatter plots comparing effective plasticity values from Yang et al^73^. (y-axis) with (left) scLSD-inferred pseudotime and (right) scLSD entropy. **(G)** Bootstrap distributions of Pearson correlations between lineage tracing-based effective plasticity (scPlasticity) and scLSD entropy, scLSD pseudotime, and the indicated stemness-associated scores. Distributions use 2,000 resamples of cells with replacement; red diamonds indicate correlations in the original data. Methods are ordered by observed correlation. scLSD entropy and the Malta score have positive correlations, while the remaining measures have negative correlations. **(H)** Five-fold cross-validated R^2^ for linear models of effective plasticity using pseudotime alone (M1), pseudotime and entropy (M2), pseudotime and the Malta score (M3), or all three predictors (M4). The model containing all three predictors has the highest R^2^ across the displayed folds.

To further investigate this possibility, we examined plasticity in the context of cancer progression in a genetically engineered mouse model of lung adenocarcinoma^73^. The developmental flow field inferred by scLSD aligned closely with experimental observations reported in this model, including a trajectory in which alveolar type 2-like cells transition through a transient plastic intermediate state before stabilizing into metastatic phenotypes (Early EMT-1) (**Fig. 5C**). In this biological system, in contrast to normal developmental trajectories, the entropy revealed a distinctive non-monotonic trend as a function of pseudotime: entropy increased along the differentiation trajectory, reaching its highest values in the “endoderm-like” and “high-plasticity” clusters before declining toward the early EMT-1 attractor state (**Fig. 5D**). This pattern is indeed consistent with “effective plasticity” measurements in the original publication, which were derived by lineage tracing independent of gene expression (**Fig. 5E**): scLSD entropy measurements showed a Pearson correlation of 0.355 with lineage tracing-based plasticity estimates. scLSD pseudotime, a surrogate for Waddington potential, showed a comparably weaker correlation with plasticity (Pearson correlation of –0.165, **Fig 5F** and **Supplementary Fig. 14C**,**15**).

Finally, we compared scLSD developmental entropy with established stemness signatures^74-80^ using lineage tracing-based effective plasticity as an orthogonal reference. Among the evaluated methods, only the stemness score of Malta et al^80^ showed a positive correlation with effective plasticity, whereas the remaining signatures were negatively correlated (**Fig. 5G, Supplementary Fig. 16**). scLSD entropy showed the strongest positive association. We therefore compared the information captured by the Malta et al^*80*^ score and scLSD entropy using multivariate models of effective plasticity. In five-fold cross-validation, adding entropy to pseudotime increased the median (R^2^) from approximately 0.03 to 0.14, compared with approximately 0.11 when the Malta score was added instead (**Fig. 5H**). A model containing pseudotime, entropy, and the Malta score achieved the highest predictive performance (*R*^2^ ∽ 0.19) indicating that scLSD entropy captures plasticity-related information that is both stronger than, and complementary to, that contained in established stemness signatures.

## Discussion

In this work, we propose an interpretation of cell differentiation based on our central hypothesis that cell states evolve within a dynamical system^81^ governed by a gradient flow and a local uncertainty term. The gradient flow naturally links to the Waddington epigenetic landscape, enabling a quantitative formulation of the Waddington potential. We further conjecture a two-dimensional representation of differentiating cells, the differentiation state, which captures sufficient information to position cells along developmental trajectories. This representation allows us to define developmental entropy, interpreted throughout this study as a quantity associated with plasticity.

We implemented this framework in scLSD, which takes time-ordered single-cell gene-expression data as input and reconstructs the underlying dynamics by inferring the cell state, differentiation state, and Waddington potential. Applying scLSD to well-characterized biological processes such as hematopoiesis, we found that the inferred dynamics align with known differentiation trajectories and converge to biologically meaningful attractor states, indicating stable flow fields. To assess predictive power, we further examined scLSD’s ability to generalize to unseen cell types across multiple datasets. During inference, scLSD recovered expected differentiation trajectories and produced cell-fate predictions consistent with baseline models or ground truth in previously unseen cell populations. It should be noted that CellRank and related graph-based methods do not provide a directly matched inductive baseline for cells excluded during fitting, so we use their full-dataset performance only as contextual reference for the held-out experiments.

We next examined the gene-level structure of the scLSD-inferred Waddington potential. Although the Waddington potential describes the global dynamics of the system, it remains a mathematical abstraction unless it can be interrogated at the level of individual genes. To this end, we performed in silico perturbation experiments in three developmental systems, zebrafish axial mesoderm, zebrafish pigmentation and mouse cortex development. scLSD identified genes whose perturbation substantially altered the inferred dynamics, suggesting their roles in fate decisions. These predictions were supported by the literature, with many highlighted genes linked to relevant pathways and phenotypes in these systems. We further compared scLSD-predicted log fold-changes in fate abundance with experimentally observed log fold-changes in cell-type abundance for perturbations in the axial mesoderm and pigmentation systems, and found strong agreement between predicted and measured effects.

Finally, we evaluated developmental entropy as a quantitative surrogate for cellular plasticity. Across multiple developmental systems, entropy decreased along pseudotime, consistent with the expected loss of plasticity during differentiation. Application of scLSD to a cancer progression model, however, revealed a different pattern: scLSD inference on the lung cancer progression dataset showed a non-monotonic pattern, whereby entropy rose toward a highly plastic intermediate state and then declined as cells stabilized into metastatic states, consistent with orthogonal plasticity measurements previously reported in this system^73^. This association was further supported by comparison with established stemness signatures: scLSD entropy showed the strongest positive association with measured effective plasticity, whereas most stemness scores were negatively correlated. These results suggest that developmental entropy captures a component of plasticity that is not readily reflected by conventional stemness measures. More broadly, the transient increase in plasticity observed during tumor progression is consistent with non-genetic state transitions that can promote epithelial–mesenchymal transition, metastasis, therapy resistance, and immune evasion^82,83^.

We highlight several implicit features of our framework that collectively position it as a systematic approach for studying differentiation dynamics. Beyond reconstructing latent dynamical processes consistent with observed temporal structure, the model exhibits predictive capacity, including generalization to unseen cell types and in silico gene perturbations. This generalizability reflects agreement between the inferred dynamical structure and the underlying biological process of differentiation, enabling extrapolation beyond the specific cell populations observed during training. Importantly, the representation learning task for the cell state is shaped not only by the expression profile of cells but the temporal structure of the developmental system. The temporal structure can help distinguish features whose current values are informative of subsequent state evolution from features that are merely correlated with the same developmental state. This does not establish causal identifiability, but it provides additional information relevant to directional dependencies that is absent from a purely static representation. Furthermore, the notion of cell state in our framework is intrinsically modality-agnostic: the notion of cell state is flexible and can be defined in terms of any biological representation appropriate to the context. This generality enables natural extensions of the approach to additional data modalities beyond RNA sequencing. Together, these implicit features suggest possible extensions of our framework which can be used to study problems such as inferring time dependent gene regulatory networks or the role of alternative splicing in cell fate decisions or cellular plasticity^84-91^.

A particularly intriguing extension would be to incorporate spatial information from spatial transcriptomic assays^92-95^. scLSD treats the dynamics of each cell as a function of its inferred cell state and therefore does not explicitly model microenvironmental influences or interactions with neighboring cell populations. Spatial measurements provide a direct way to enrich this representation by augmenting the state of a focal cell with features describing its local niche, such as the identities, states, or signaling activities of nearby cells. This would allow the inferred dynamics to depend jointly on the intrinsic state of the cell and on the evolving state of its microenvironment, effectively extending scLSD from single-cell autonomous dynamics to a coupled dynamical system of cells and niches. In settings where the microenvironment is approximately homogeneous or slowly varying, or where its effects are already sufficiently encoded in the transcriptional state of the focal cell, such a formulation would reduce to the current scLSD framework. Biologically, incorporating spatial context could enable direct investigation of signaling-dependent changes in developmental trajectories, the role of cell-cell interactions in fate decisions, and context-specific state transitions in disease, for example how stromal or immune niches reshape tumor-cell dynamics.

Some recent approaches^96-98^ have pursued more explicit geometric reconstruction of Waddington landscapes, using reaction-coordinate theory, Morse-Smale topology, or signal-parametrized potentials to identify attractors, saddles, unstable manifolds, and bifurcations. These formulations provide greater resolution of landscape topology, but generally rely on stronger assumptions about the dimensionality, decision structure, relevant molecular features, or external signals of the system being studied. scLSD takes a complementary approach: it jointly learns a flexible neural potential and latent cell-state representation from high-dimensional transcriptomic data without prespecifying a particular bifurcation topology or reaction coordinate. This sacrifices explicit reconstruction of saddles and bifurcation structure in favor of applicability across diverse developmental systems and predictive tasks such as fate assignment and perturbation analysis. A possible future direction would be to bridge these two regimes by augmenting scLSD with more explicit geometric or topological constraints on the learned landscape while retaining its end-to-end representation learning and applicability to high-dimensional, less-characterized systems. Such an extension could improve resolution of critical landscape features without requiring the strong a priori structural assumptions used in more specialized reconstruction frameworks. For instance, identifying saddle points and barrier heights could enable transition rates to be modeled through Kramers-type rate theory^99^ rather than inferred only from local potential differences, providing a more mechanistic description of fate transitions. Such extensions could improve the resolution and physical interpretation of the inferred Waddington landscape without requiring the strong a priori structural assumptions used in more specialized reconstruction frameworks.

A current limitation of scLSD is its dependence on a prior temporal ordering. Some notion of temporal information is necessary for learning dynamics from snapshot data, but this information need not take the form of a fixed pseudotime supplied before training. In the present implementation, the prior ordering constrains the trajectories encountered during optimization and therefore influences the inferred dynamics, as demonstrated by our pseudotime perturbation analyses. A natural extension would be to weaken this dependence by jointly refining temporal ordering and the dynamical model. For datasets with coarse experimental time points, for example, an initial ordering could be inferred subject to these temporal constraints and subsequently updated iteratively using the dynamics learned by scLSD, analogous to an expectation-maximization procedure in which temporal assignment and dynamical inference are alternated until convergence. Such a formulation would retain the temporal information required for dynamical modeling while reducing reliance on an externally specified pseudotime.

A separate limitation of scLSD concerns cyclic dynamics in gene-expression space. Even under the idealized assumption of perfect reconstruction, such that the decoder defines an injective mapping between the latent state and the expression manifold, gradient structure in the latent space need not imply a curl-free vector field in ambient gene-expression coordinates. More generally, if the latent developmental manifold is endowed with a non-Euclidean Riemannian metric, Riemannian gradient flow dynamics can give rise, after pushforward through the decoder, to expression-space dynamics with a nonzero Euclidean curl component^100^. This geometric generalization, however, still does not permit genuine periodic orbits or limit cycles, because the Waddington potential remains a Lyapunov function and decreases monotonically along nonstationary trajectories. Capturing truly cyclic processes such as cell-cycle oscillations would therefore require an additional dynamical component, for example a fast phase or rotational variable coupled to the slower landscape-driven differentiation dynamics. Such a multiscale formulation could preserve the long-term gradient structure of differentiation while explicitly modeling shorter-timescale recurrent behavior.

## Methods

### The scLSD framework

scLSD models time-ordered single-cell gene expression through a hierarchical probabilistic representation. Let *G* denote the number of genes, *D* the cell-state dimension and *d*_*B*_ = 2 the differentiation-state dimension, with 2 < *D* ≪ *G*. Observed raw counts are denoted by 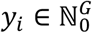 and their log-normalized counterparts by *x*_*i*_ ∈ ℝ^*G*^. A synthetic trajectory is *X* = (*x*_0_, *x*_1_,…, *x*_*T*−1_), paired with raw counts *Y*. We use *k, t* for discrete trajectory step and continuous ODE time respectively. *Z, B, L* denote random variables assigned to cell states, differentiation states and library sizes. *z, b, l* denote their realizations respectively.

We then define an encoder, implemented as a neural network,

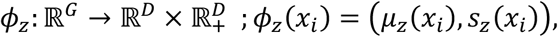

and the distribution of the latent cell states can be written as

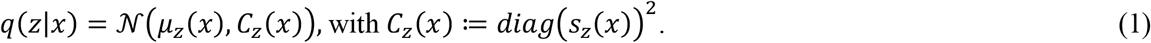

A conceptual stochastic model of cell state evolution is the Itô SDE

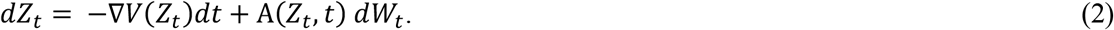

*V*: ℝ^*D*^ *→* ℝ_*+*_ represents the Waddington potential landscape, *W*_*t*_ is a *D* -dimensional Wiener process, A: ℝ^*D*^ × ℝ_*+*_ *→* ℝ^*D*×*D*^ is a diagonal noise-amplitude matrix, with infinitesimal noise covariance being AA^T^*dt*.

For integrable solutions of the SDE, the mean satisfies 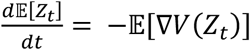. Replacing the expectation of the gradient by its value at the mean gives a first-order drift approximation^101^. The cell state representation described in scLSD specifies the mean by the first-order drift approximation but obtains the covariance from the expression encoder instead of solving the covariance moment closure equations:

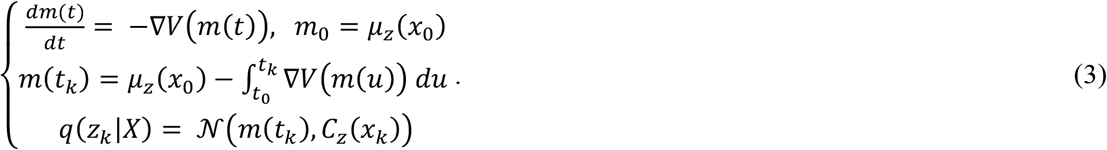

scLSD parameterizes the Waddington landscape *V* as a neural network and defines the latent dynamics through its negative gradient, 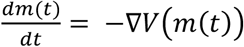, yielding a neural ODE that can be trained efficiently by gradient-based optimization.

As discussed earlier, we also obtain two-dimensional differentiation state representations using the cell state embeddings. Let 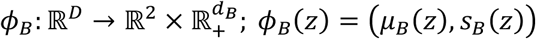 be the encoder function for the differentiation state representations, *B*_*i*_s, where

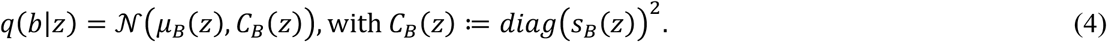

Under the Gaussian variational posterior defined in Eq. (4) the developmental entropy assigned to a particular cell state is

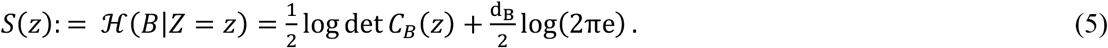

The hierarchical encoding tasks within scLSD are complemented by a corresponding decoding framework that reconstructs the gene expression profiles from the latent representations. To initiate the hierarchical decoding process, we first establish a prior distribution for the differentiation states. Let 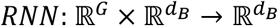 be a recurrent neural network, such that *h*_*t*_ := *RNN*(*x*_*t*_, *h*_*t*−1_) . The differentiation state representations are then assumed to follow a prior distribution *p*(*b*_*t*_|*x*_0_,…, *x*_*t*_) = *N*(*h*_*t*_, *I*), where *h*_*t*_ captures the temporal dependencies in the differentiation states through the RNN hidden states and *I* denotes the *d*_*B*_ × *d*_*B*_ identity matrix.

The decoding process proceeds through a probabilistic mapping that transforms the differentiation state representations back to the latent cell state space,

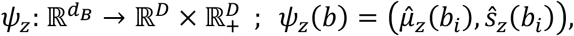

such that the reconstructed cell state representation is given by

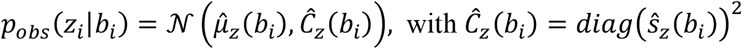

The final decoding step reconstructs the gene expression profiles from the latent cell states. Rather than decoding *z*_*i*_ to log-normalized expression *x*_*i*_, we choose to reconstruct the original raw count data to preserve the discrete nature and distributional properties of single-cell RNA sequencing data. We model the likelihood using a zero-inflated negative binomial (ZINB) distribution,

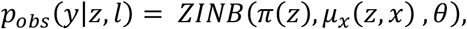

where *π* (*z*), *μ*_*x*_(*z, x*), *θ* are zero-inflation logits, mean and dispersion parameters, respectively. The zero-inflation and the mean satisfy the following relationships:

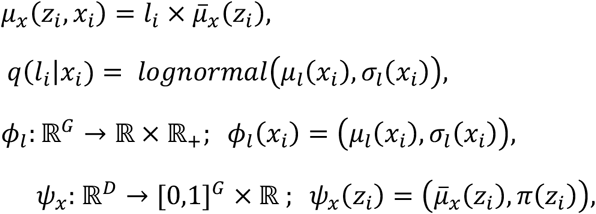

where *l*_*i*_ is the library size parameter, *ϕ*_*l*_ is the library size encoder and *Ψ*_*x*_ is the gene expression decoder function. It should be noted that the library size parameter also requires a prior given by

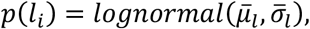

where 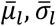 correspond to the mean and the standard deviation of the observed library sizes of the dataset.

### scLSD’s objective function

The scLSD framework is trained by maximizing the evidence lower bound (ELBO) associated with the hierarchical variational scheme described above. This variational formulation treats the cell states and differentiation states as latent variables, while the observed raw count data serves as the evidence. The hierarchical structure of our variational scheme, spanning the high-dimensional gene expression profiles, the intermediate cell states, and the low-dimensional differentiation states, necessitates careful consideration of the conditional dependencies and approximate posterior distributions at each level of the hierarchy.

The ELBO can be decomposed into two complementary terms that capture different aspects of the hierarchical reconstruction. The first term,

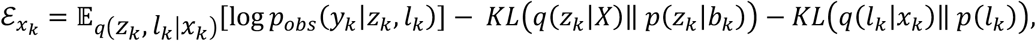

quantifies the reconstruction fidelity of raw counts from the latent cell states. This term ensures that the learned latent representations *z*_*t*_ contain sufficient information to accurately reconstruct the observed gene expression counts *y*_*k*_ through the ZINB likelihood model. The second term,

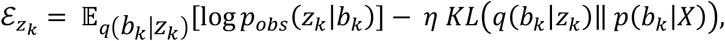

addresses the reconstruction of cell state representations from differentiation state representations. This term balances the ability to reconstruct the cell states from the differentiation states against the regularization imposed by the prior distributions. In these equations, *KL*(. ‖ . ) is the Kullback–Leibler (KL) divergence, *η* represents the KL annealing hyperparameter (detailed in supplementary notes), and the distributional notations follow the definitions established previously. Hence, the total ELBO for time series *X* can be written as:

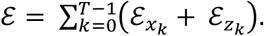

Optimization of this variational scheme requires maximizing the ELBO, which is equivalent to minimizing the negative likelihood loss function

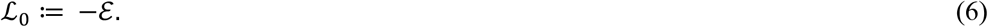

Furthermore, we introduce an optimal transport regularization term that constrains the learned dynamics to follow physically plausible paths:

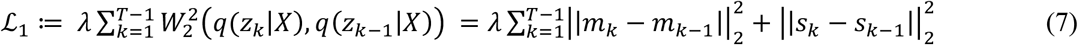

where *W*_2_ is the 2-Wasserstein distance, *λ* is a regularization coefficient, and,.,_2_ denotes the Euclidean distance.

### Dynamical constraints and thermodynamic interpretation of developmental entropy

Throughout this work, we define the developmental entropy Eq. (5). This quantity is thermodynamically motivated, but its relationship to thermodynamic entropy is not immediate and requires clarification. Mechanistically, *S*(*z*) is the conditional differential entropy of the variational posterior *q*(*b*|*z*) and is therefore shaped by the reconstruction–regularization trade-off associated with compressing the cell state into the differentiation-state representation. Importantly, however, this compression takes place within a latent space that is subject to stronger structural constraints than those of a conventional variational autoencoder.

In an unconstrained VAE, the latent representation is generally non-identifiable under a diffeomorphism of the latent coordinates. In scLSD, the cell-state representation is additionally constrained by the dynamical law 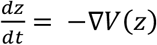. Consequently, a latent-space diffeomorphism is admissible only if the pushforward of the learned vector field remains representable as a gradient flow. The family of diffeomorphisms preserving an scLSD solution is therefore a strict subset of the diffeomorphisms available to the corresponding unconstrained variational model. Thus, the reconstruction–regularization trade-off determining *q*(*b*|*z*) operates within a dynamically restricted family of latent geometries, in which both the representation and the gradient-flow structure must remain mutually compatible.

#### Theorem 1

Let Ω < ℝ^*D*^be open and connected and assume the VAE model class is closed under smooth invertible reparameterizations of the latent variable. Define the ELBO by

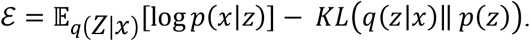

Then the following holds:

1. Let 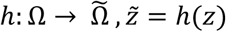. Transform the encoder and prior by the pushforward 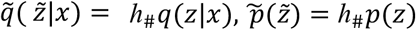 and define the transformed decoder 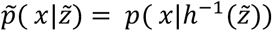. Then 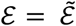
2. Suppose that the latent state satisfies 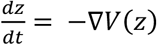 for some *V* ∈ *C*^2^(Ω)and let denote the Euclidean metric duality between vector fields and 1-forms. Then the diffeomorphism *h* gives an equivalent representation within the same class of Euclidean gradient-flow models if and only if *β*_*h*_:= (*h*_∗_(−A*V*))^*b*^ is exact on *h*(Ω).
3. Let dim Ω ≥ 2, A*V* ≢ 0, then Euclidean gradient structure is not preserved by arbitrary smooth invertible reparameterizations.

See **Supplementary notes** for the proof.

Although the scLSD latent coordinates are not uniquely identifiable, their residual degeneracy is restricted to transformations that preserve the gradient-flow structure. This equivalence class contains at least Euclidean isometries. We expect this coordinate-level non-identifiability to have limited effect on biologically meaningful outputs such as cell-fate predictions, ordering by Waddington potential, or developmental flow projected onto a fixed external representation. Because optimization may introduce variability beyond exact members of this equivalence class, we assessed this practical identifiability empirically by retraining scLSD across random seeds and comparing these downstream quantities (**Supplementary Fig. 17**).

This restricted non-identifiability does not render differential entropy invariant to reparameterization, but it limits the latent coordinate transformations under which equivalent scLSD solutions can be constructed. Accordingly, developmental entropy should be interpreted primarily through its relative variation across cells within a learned dynamical representation, rather than as an absolute coordinate-independent entropy scale.

A more explicit connection to thermodynamics follows by interpreting the cell state *z* as a microscopic state and the differentiation state *B* as a developmental macrostate. In equilibrium statistical mechanics, the entropy associated with a macrostate characterizes the heterogeneity of microscopic configurations compatible with that macrostate and can be expressed as *H*(*Z*|*B*). This quantity is distinct from *H*(*B*|*Z*): the former describes microscopic uncertainty conditional on a developmental macrostate, whereas the latter describes uncertainty in developmental state conditional on a particular cell state. Nevertheless, the two quantities are related through the joint distribution of *Z* and *B*.

To make this relationship explicit, consider a local decomposition of the cell state, *z* = (*ζ, ξ*) ∈ ℝ^*D*^, where 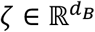 contains the developmental modes of the cell state represented by the differentiation state, while 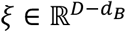 contains transverse cell-state variations. These transverse modes may capture biological or technical variation within the cell state but, by construction, do not directly determine position along the differentiation axis.

Under a local-equilibrium approximation of the Waddington potential, and assuming that the learned differentiation-state distribution reproduces the distribution of the developmental modes, the conditional thermodynamic entropy of the cell-state ensemble can be decomposed as *H*(*Z*|*B*) = *S*_⊥_ *+ H*(*B*|*Z*), where *S*_⊥_ represents the entropy associated with fluctuations along the transverse modes *ξ*. For a locally quadratic Waddington potential around *z*^∗^ this contribution is determined by the curvature of the potential along the transverse directions.

#### Theorem 2

Let cell state be decomposed as *Z* = (*Z, Z*_⊥_)with realization *z* = (*ζ, ξ*) ∈ ℝ^*D*^ where 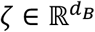 contains the developmental modes and *ξ* contains the remaining transverse modes. Assume that, in a neighborhood of a nondegenerate local minimum *z*^∗^ = (*ζ*^∗^, *ξ*^∗^), the Waddington potential is locally quadratic,

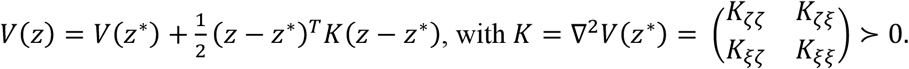

Assume further that the local equilibrium density is Boltzmann distributed *p*_*Z*_(*z*) ∝ *e*^−*βV*(*z*)^, where *β* is the inverse temperature, and that at the variational optimum the marginal distribution of the differentiation state matches that of the developmental modes, *p*_*B*_ = *p*_‖_ . Then 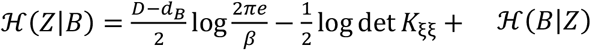, where 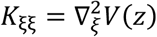 is the curvature of the Waddington potential along the transverse modes.

See **Supplementary notes** for the proof.

The entropy decomposition provides a thermodynamic interpretation of developmental entropy without identifying *H*(*B*|*Z*) with the full microscopic entropy of the cell state. Rather, developmental entropy captures the component associated with uncertainty in developmental macrostate after separating the contribution of transverse microscopic modes.

### Relation of scLSD dynamics to SDE formulation

A full stochastic implementation of Eq. (2) would require propagating not only the mean cell state but also its uncertainty. Under a Gaussian moment approximation cell state is normally distributed as in Eq. (1). However, the stochastic treatment requires coupled evolution equations for *m*_*t*_ and *C*_*t*_. The current scLSD implementation does not perform this full Gaussian moment closure: it propagates only the first-moment dynamics with Eq. (3), while the covariance is inferred locally from the variational encoder rather than dynamically propagated.

This distinction also separates scLSD from a purely deterministic ODE formulation. A deterministic latent ODE represents each state only by its mean trajectory *m*_*t*_and evaluates a nonlinear objective *l* directly at that point, *J*_ODE_ = *P*(*m*_*t*_) . In contrast, scLSD evaluates the variational objective over a Gaussian neighborhood of the propagated mean, so that *J*_scLSD_ = E[*P*(Z_*t*_)]. For a smooth loss, a second-order Taylorexpansion around *m*_*t*_ gives 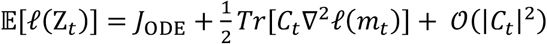.

The same second-order correction arises under a Gaussian moment approximation to Eq. (2), with the distinction of covariance evolving as 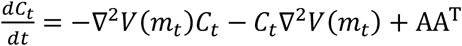. The essential difference is therefore not the form of this correction, but the origin of *C*_*t*_. In the SDE approximation, covariance is dynamically coupled to the curvature of Waddington landscape and the diffusion term, whereas in scLSD it is inferred locally by the variational encoder and is not constrained to satisfy this covariance evolution equation.

### Optimization

Based on the framework established in the previous sections, the complete loss function is defined as the combination of the variational objective function of Eq. (6) and the optimal transport regularization term of Eq. (7):

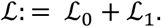

The optimization task is then defined as

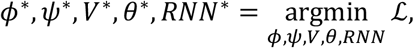

where *ϕ* = (*ϕ*_*z*_, *ϕ*_*B*_) and *Ψ* = *Ψ*_*z*_, *Ψ*_*x*_) are the encoder and the decoder functions for the cell states and the differentiation states, *V* is the neural network which estimates the Waddington potential, *θ* encompasses the overdispersion parameters of the ZINB distribution and *RNN* is the recurrent neural network used in the prior of the differentiation state representation.

To implement this optimization framework we employed Pyro^102^, a probabilistic programming language built on Python and PyTorch^103^, which provides native support for variational inference and stochastic optimization. The loss function was estimated using Pyro’s Stochastic Variational Inference (SVI) algorithm^40^, coupled with the Adam optimizer^104^ enhanced with warm restarts to improve convergence stability and escape local minima. The SVI framework enables efficient gradient estimation through automatic differentiation, while handling the stochastic nature of the variational approximations inherent in our hierarchical model.

### Time series generation from static single cell snapshots

A fundamental challenge in modeling single-cell dynamics as a dynamical system is the absence of high-resolution single-cell time series data. To address this limitation, we synthetically generate time series trajectories from static single-cell snapshots by leveraging pseudotime information. Consider single-cell log-normalized gene expression profiles of *N* cells represented as 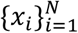, where *x*_*i*_ ∈ ℝ^*G*^ denotes the log-normalized expression vector of highly variable genes for cell i. Each cell i is associated with a pseudotime measure *t*_*i*_ ∈ [0,1], where 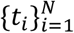 represents the inferred temporal ordering of cells along a developmental trajectory. Given a pre-computed k-nearest neighbor (KNN) adjacency matrix *w* ∈ ℝ^*N*×*N*^, we define a pseudotime-aware transition matrix as:

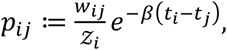

where *Z*_*i*_ is the normalization coefficient which ensures 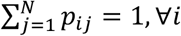, and *β* represents the inverse temperature parameter for the Boltzmann distribution.

The transition matrix *p*_*ij*_ incorporates two complementary terms that jointly govern the stochastic dynamics of synthetic trajectory generation. The KNN connectivity term *w*_*ij*_ enforces spatial localization of random walks by restricting transitions to occur only between nearest neighbors in the gene expression space, thereby preserving the local manifold structure of the data. This connectivity constraint ensures that synthetic trajectories remain biologically plausible by preventing unrealistic jumps between distant cellular states.

The pseudotime-dependent exponential term 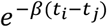 introduces directional bias toward states with higher pseudotime values. When *t*_*j*_ > *t*_*i*_, this term amplifies the transition probability, favoring forward progression along the developmental trajectory. Conversely, when *t*_*j*_ < *t*_*i*_, the term suppresses backward transitions, creating an implicit temporal direction that guides synthetic trajectories toward more mature cellular states. The inverse temperature parameter *β* controls the degree of stochasticity in these pseudotime-driven transitions: higher *β* values create more deterministic, directed trajectories that strictly follow pseudotime gradients, while lower β values allow for greater stochastic exploration around the pseudotime-defined path. This parameterization enables fine-tuning of the balance between biological realism (through KNN connectivity) and temporal directionality (through pseudotime bias) in the generated synthetic time series.

Using the pseudotime-aware transition matrix *p* ∈ ℝ^*N*×*N*^, we generate synthetic time series data through initializing random walks. For a random walk *s*, we first randomly select an initial cell state *s*_1_ ∈ {1, 2,…, *N*} from the dataset. The subsequent states in the trajectory are determined by iteratively sampling from the transition probabilities: 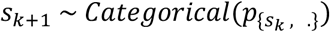, where *p*_{*i*, .}_ represents the *i*-th row of the transition matrix corresponding to the transition probabilities from state i to all other states. Then the gene expression profile of the walk is given by 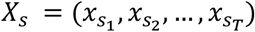, where *T* is the fixed hyperparameter corresponding to walk length.

This process is repeated for a predetermined walk length *T* and across M independent walks, yielding a synthetic time series dataset of shape (*M, T, G*), where *M* represents the number of generated trajectories, *T* denotes the temporal length of each trajectory, and *G* corresponds to the number of highly variable genes.

To generate more biologically accurate time series data when cluster phylogeny information is available, we implement a complementary approach that leverages phylogenetic relationships to constrain cellular transitions. This phylogeny-informed method addresses two critical limitations of pseudotime-based transitions: backward and parallel transitions. Backward transitions occur when cells transition from descendant clusters to their ancestral states, violating the natural developmental hierarchy. While pseudotime incorporation should theoretically limit such transitions, inaccuracies in pseudotime inference methods and non-uniform pseudotime distributions can still permit biologically implausible backward movements along the developmental trajectory. The more critical challenge involves parallel transitions, where cells transition between clusters located on separate branches of the phylogenetic tree. For instance, in endocrinogenesis datasets, cell types such as beta and alpha cells exhibit high transcriptional similarity, leading to spurious connectivity that permits transitions between these parallel lineages.

Given the cluster labels for the dataset 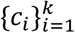, and a predefined descendent set for each cluster *D*(*c*_*i*_) = {*all cells in c*_*i*_ *or* descendent clusters of *c*_*i*_}, we define a design matrix *A* ∈ ℝ^*N*×*N*^ as 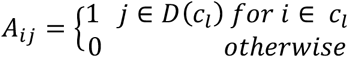.

Then, one can refine the transition matrix as

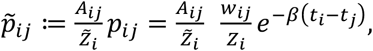

where 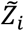 ensures 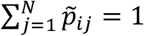. Therefore, the resulting transition matrix 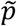 effectively prohibits both backwards and parallel transitions, thereby generating synthetic time series that strictly adhere to the known phylogenetic constraints, while maintaining the pseudotime-driven directionality.

### Prior pseudotime initialization

scLSD requires an initial temporal ordering to construct the transition matrix used for generating training trajectories. When a suitable ordering was available from the original analysis, we used it directly: bone marrow^11^ pseudotime was obtained from Palantir^11^, and lung adenocarcinoma^73^ pseudotime from PhyloVelo^105^. For zebrafish axial mesoderm^49^, we used a CellRank^44^ transition matrix constructed from CytoTRACE^106^-derived developmental potential. For the remaining datasets, prior pseudotime was inferred from known developmental structure using the scLSD preprocessing procedure. Briefly, biologically established origin clusters and, when available, cluster-level lineage relationships were supplied and cells were ordered within this structure using their transcriptomic neighborhood relationships to obtain a continuous prior pseudotime scaled to [0,1]. This prior ordering was subsequently used to construct the pseudotime-biased transition matrix.

### Posterior cell-to-cell transition matrix calculation

After training the scLSD model, we computed a posterior cell-to-cell transition matrix that captures the learned cellular differentiation dynamics. The transition probability from cell i to cell j was calculated as a Boltzmann distribution over potential changes:

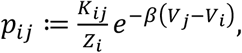

where *p*_*ij*_ is the transition probability of cell i to cell j, *Z*_*i*_ is the normalization coefficient and *V*_*i*_ is the Waddington potential of cell i, respectively. In these equations, *K* is the KNN graph adjacency matrix, *β* is the Boltzmann distribution inverse temperature parameter. For all datasets, we set *β* = 1.

### Velocity projection

To visualize low-dimensional velocity streamlines, we employed the CellRank method, which involves projecting the cell-to-cell transition matrix onto a latent embedding. Consider a cell i with neighborhood *N*(*i*), and let *p* denote the posterior transition matrix. The projected velocity *v*_*i*_ of cell i on a latent embedding *u* is defined as:

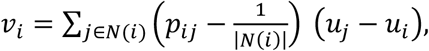

where |*N*(*i*)| represents the cardinality of the neighborhood *N*(*i*) and *v*_*i*_ is the velocity vector of cell i, sharing the same dimensionality as *u*_*i*_. In the special case where *u*_*i*_ corresponds to a two-dimensional latent embedding, such as UMAP or t-SNE, the velocity vector field can be directly visualized on this embedding.

### CBDir Metric

To quantitatively assess the accuracy of the scLSD-predicted transition matrix, we employed the CBDir metric. As previously described, the transition matrix enables the calculation of a velocity vector field on the data manifold. Given a known phylogeny or lineage tree of cell clusters, CBDir estimates the degree to which the predicted velocities align with the expected developmental progression. Specifically, for each cell located at the interface between two adjacent clusters in the phylogeny, CBDir computes the cosine similarity between the predicted velocity vector and the direction vector pointing from the source cluster to the target cluster. The overall CBDir score is then obtained by averaging these similarities across all relevant cross-boundary cells. A higher CBDir value indicates that the predicted transitions are consistent with the expected biological lineage, providing a quantitative measure of the directionality correctness of the inferred cell state transitions.

The CBDir metric is defined as:

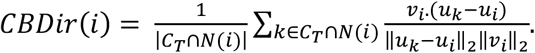

In the equation above, *v*_*i*_ denotes the predicted velocity vector for cell i, while *u*_*i*_ and *u*_*k*_ represent the positions of cell i and cell k in the latent embedding space, respectively. The set *N*(*i*) consists of the neighbors of cell i, *C*_*T*_ denotes the set of cells belonging to the target cluster adjacent to cell i’s source cluster, and |*C*_*T*_ ∩ *N*(*i*)| is the cardinality of the set *C*_*T*_ ∩ *N*(*i*).

To calculate the CBDir metric for both RNA velocity methods and pseudotime, we utilized CellRank’s velocity and pseudotime kernels, respectively, to construct the transition matrix. Subsequently, we employed CellRank’s projection function, TmatProjection, to compute the velocity projection for each cell. Using these projected velocities, we calculated the CBDir score for each cell according to the formula described above. The overall CBDir value reported in Figure 2E corresponds to the mean CBDir score averaged across all cells. Details regarding the cluster phylogeny used for each dataset can be found in the supplementary notes. Since CBDir reflects both the inferred velocity field and the geometry of the embedding space, its value can vary across representations. Accordingly, in addition to the standard UMAP or tSNE embeddings, we report CBDir scores on PCA, latent cell state, and differentiation state embeddings (Supplementary Fig. 4).

For the OT model comparison (OT-CFM^32^, PRESCIENT^33^ and scNODE^45^) inferred local velocities were converted into cell-to-cell transition probabilities using their positive cosine alignment with neighboring-cell displacements on a common connectivity graph. Specifically, let *z*_*i*_ and *v*_*i*_ denote the cell-state representation and inferred velocity of cell *i* respectively. For each *j* ∈ *N*(*i*) we compute 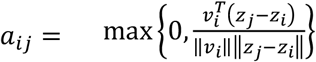, and normalize these scores to obtain 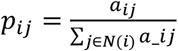. When no neighboring displacement had positive alignment with *v*_*i*_, transition probability was distributed uniformly across the neighbors. The resulting transition matrices were projected onto the same UMAP embedding and evaluated using CBDir, with scores averaged equally across annotated lineage edges.

### Benchmarking against continuous optimal transport models

We compared scLSD with OT-CFM, PRESCIENT and scNODE on human germline differentiation^47^, human neurogenesis^46^ and pancreas development^30^. The comparison models were trained on all cells and observed time groups using seed 42 and method-specific optimization settings, with checkpoints selected by training loss. OT-CFM and PRESCIENT used ten principal components, whereas scNODE and scLSD were evaluated in ten-dimensional encoder-mean cell states. The comparison models used ordinal sampling-time labels; scLSD used its dataset-specific pseudotime prior, with lineage constraints for human germline and pancreas. These experiments evaluated fitted dynamics on the observed populations. All benchmark trajectories were propagated deterministically with Dopri5^107^, using absolute and relative tolerances of 10^-5^. PRESCIENT was evaluated using its drift without diffusion noise. scLSD trajectories were propagated up to *t* = 5; the comparison models were integrated from each cell’s sampling stage to the final stage, giving final-stage cells zero propagation duration.

We reconstructed trajectories from all cells with prior pseudotime below 0.1, 0.3 or 0.5, evaluating each threshold separately. Stability was quantified for each trajectory as the ratio of mean velocity norm over its final 10% to that over its initial 10%. We reported the median and interquartile range across trajectories and the unweighted mean of dataset medians; the main stability comparison used the 0.5 threshold.

To assess reconstruction of developmental progression, propagated points were annotated with the pseudotime of their nearest observed reference cell. We compared the propagated and observed pseudotime distributions using one-dimensional Wasserstein distance and squared maximum mean discrepancy (MMD^2^) with a radial basis function kernel. Temporal ordering was assessed by the median within-trajectory

Spearman correlation between integration time and projected pseudotime. State-distribution agreement was evaluated using MMD^2^ after scaling each model’s coordinates by its observed-reference mean and standard deviation (**Supplementary Fig. 6**).

### Propagating cells in cell state space and predicting cell fates

After training, the scLSD model enables the propagation of cells in the cell state space. Consider a cell i with a log-normalized gene expression profile *x*_*i*_. Its corresponding cell state is computed as *z*_0,*i*_ = *ϕ*_*z*_(*x*_*i*_) via the cell state encoder. This latent representation then serves as the initial condition for a neural ODE that captures the cell’s temporal dynamics,

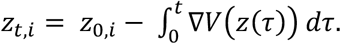

Although this integral is analytically intractable, it can be efficiently approximated using numerical solvers. Throughout our work, we used Torchdiffeq’s dopri8^38,107^method to numerically solve the neural ODE. As *t* increases, *z*_*t,i*_ converges to a steady state *z*_∞,*i*_, corresponding to a local minimum of the potential function *V*(*z*) and representing the predicted cell fate for the initial state *z*_0,*i*_ . However, *z*_∞,*i*_ may not precisely coincide with a point on the observed data manifold. To assign a cell fate cluster, we project *z*_∞,*i*_ onto the nearest observed cell state *z*_*j*_ on the manifold, defined as:

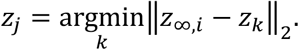

The cell j is subsequently treated as the effective fate of cell i for the purposes of cell fate assignment.

To quantify uncertainty in fate commitment, we sampled initial cell states from *q*(*z*|*x*), propagated each sample through the learned flow field and assigned its endpoint to the nearest observed cell state. The fraction of samples assigned to each cluster defined cell-specific fate probability. Endpoint assignment included all observed clusters, allowing trajectories to remain associated with intermediate states (**Supplementary Fig. 10**).

### Lineage-aware fate prediction accuracy

We defined an allowed set of terminal fates for each annotated source population using the dataset-specific lineage hierarchy. For each cell, fate accuracy was the summed probability assigned to these allowed fates, equivalent to one minus the probability assigned to incompatible fates. scLSD probabilities were estimated from ten sampled initial states per cell; RNA velocity and pseudotime fate probabilities were obtained using CellRank. We reported the mean and median across eligible cells, excluding populations without a defined lineage assignment. The cross-method fate comparison included bone marrow, erythroid and dentate gyrus and used Cell2Fate^14^, UniTVelo^15^, VeloVI^16^, both scVelo^4^ models and prior pseudotime as comparators. Overall scores were unweighted means of dataset-level scores, with unavailable method–dataset results retained as missing.

For OT models (OT-CFM^32^, PRESCIENT^33^ and scNODE^45^) fate predictions were obtained by propagating each eligible cell and assigning the endpoint a cluster through inverse-distance-weighted voting among its 25 nearest reference cells in the model’s state space. All annotated clusters were eligible for assignment. A prediction was correct when its assigned cluster belonged to the source population’s allowed terminal-fate set. Accuracy was calculated within each source lineage, including terminal source populations, and macro-averaged across lineages (**Supplementary Fig. 7**).

### kNN baseline for bone marrow fate assignment

To assess whether fate assignments could be recovered from local transcriptomic similarity alone, we constructed a 15-nearest-neighbor baseline in PCA space. Terminal reference cells were defined as Mono, DCs, Ery_2, and Mega, with Mono_1 and Mono_2 merged into a single Mono fate. For each remaining cell, the fractions of its 15 nearest terminal reference cells belonging to each fate defined a four-dimensional fate-probability vector, with the highest fraction determining the hard fate assignment. We compared this vector with the fate assigned by the full-data scLSD model using cell-wise Pearson correlation and summarized the correlations within each source cluster.

### In silico gene perturbation

As discussed before, after training, each cell’s state can be propagated over time by numerically integrating the learned cell state dynamics. To model genetic perturbations, we modified the original gene expression profiles, for example by setting a gene to zero for knockouts and encoded these altered profiles as initial conditions in the cell state space. To distinguish between transient and sustained gene perturbations, we further implemented an iterative procedure that repeatedly enforces the perturbation during trajectory propagation, thereby maintaining the altered gene expression throughout the simulation.

Before describing the experimental setup, we introduce our notation. Let *x*_*i*_ denote the unperturbed and 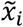 the perturbed, log-normalized gene expression profile of cell i. The corresponding cell states in the latent space are given by 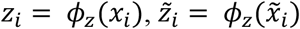, where *ϕ*_*z*_ denotes the encoder. To map any point *z* ∈ ℝ^*D*^ in the cell state space back onto the data manifold, we define the projection operator 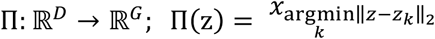. This operator projects an arbitrary latent state *z* onto its nearest neighbor among observed data points, retrieving the corresponding gene expression profile. We denote the propagation of a cell state *z*_0_ in cell state space under the dynamics *f*(*z*) for time *t* as

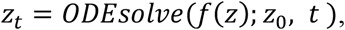

which represents the numerical solution to the initial value problem

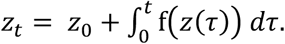

Hence, the iterative scheme of gene perturbation setup will be as follows.

For each cell *i* and a chosen step size *ε*:

1. **Initialization**: Encode the perturbed gene expression profile to obtain the initial cell state:

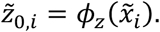
2. **Propagation**: Propagate the cell state for a small time interval *ε* using the learned dynamics:

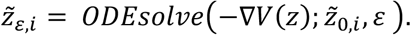
3. **Projection**: Map the propagated cell state back to the data manifold:

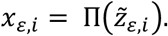
4. **Re-application of Perturbation**: Apply the perturbation to obtain the modified expression profile:

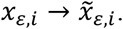
5. **Iteration**:

Set 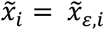 and repeat steps 1–4 until convergence.

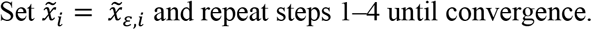

### Ablation of model components and identifiability analysis

We evaluated potential non-negativity, Wasserstein regularization and the recurrent differentiation-state prior through separate ablations, retaining the remaining dataset-specific settings. For bone marrow, dentate gyrus, erythroid and pancreas, we removed the nonnegative output constraint from the potential network and assessed trajectory convergence using the latent velocity norm and signed changes between adjacent propagated states. We varied the constant Wasserstein coefficient *λ* over 0, 10^−4^, 10^−3^, 10^−2^, 0.1, 0.5, 0.9 and 1. Encoder-ODE consistency was quantified by the area under the curve of variance-normalized squared discrepancies between propagated states and states obtained after projection to expression space and re-encoding. These comparisons used three seeds per condition and were summarized by the mean and standard error across successful runs.

To assess the recurrent prior, we replaced its learned mean with a fixed zero mean and compared differentiation-state CBDir and the signed Pearson correlation between entropy and the original pseudotime. The expanded comparison used 15 seeds per condition for bone marrow, dentate gyrus and erythroid. One-sided Mann–Whitney U tests^108^ assessed higher CBDir and more negative entropy–pseudotime correlations with the recurrent prior; reported P values were nominal (**Supplementary Fig. 2**).

For identifiability analysis we trained ten independent models for each of the bone marrow and erythroid datasets, keeping the dataset-specific configuration fixed and varying the random seed. For all 45 pairs of models, we compared cell-specific fate probability vectors and velocities projected onto a fixed external embedding using cosine similarity, and compared Waddington potentials using Spearman correlation. Fate probabilities were estimated from 40 posterior samples per cell. These quantities assess reproducibility of downstream predictions without requiring alignment of latent coordinates (**Supplementary Fig. 17**).

### Sensitivity to differentiation-state dimensionality

To evaluate sensitivity to the dimensionality of the differentiation state, we trained scLSD with differentiation-state dimensions of two, three and four on bone marrow, erythroid, pancreas and lung adenocarcinoma data, keeping the remaining dataset-specific settings and random seed fixed. We aligned cells across configurations and calculated Pearson correlations between their developmental entropy values and between entropy and posterior pseudotime from each configuration. In lung adenocarcinoma, we additionally compared entropy and posterior pseudotime with lineage tracing-based effective plasticity. Dimensionality refers to the model configuration; entropy and pseudotime remain scalar quantities (**Supplementary Fig. 14**).

### Prior pseudotime sensitivity analysis

We retrained scLSD on bone marrow, erythroid, pancreas and dentate gyrus after perturbing the initial pseudotime, using three seeds per condition and no phylogeny constraints. Pseudotime was scaled to [0,1]. Gaussian perturbations added zero-mean noise, followed by clipping and rescaling; rank perturbations displaced cell ranks uniformly within ±*k* and mapped the resulting ordering to [0,1]. Noise standard deviation and relative rank displacement *k*/*N* were varied over 0, 0.05, 0.1, 0.2, 0.3, 0.4 and 0.5. We measured Pearson correlation between posterior and original pseudotime and defined the correction ratio γ as this correlation divided by the correlation between perturbed and original pseudotime. Projected velocities were compared with the corresponding same-seed, unperturbed model using cell-wise cosine similarity, summarized by their mean and median (**Supplementary Fig. 8**).

### Data Preprocessing

Prior to applying the scLSD framework, all single-cell RNA sequencing datasets were preprocessed following standard practice in single-cell analysis to ensure comparability across datasets. We employed the Scanpy^109^ toolkit for all preprocessing steps.

Preprocessing included total-count normalization to a target sum of 10^4^ per cell using *scanpy*.*pp*.*normalize_total*, followed by log-transformation with *scanpy*.*pp*.*log1p*. Principal components were computed from the log-normalized expression profiles using *scanpy*.*pp*.*pca* with default settings (50 principal components). A k-nearest-neighbor (kNN) graph was then constructed using *scanpy*.*pp*.*neighbors* with the default number of neighbors (k = 15).

In addition, the mean and standard deviation of the empirical library size distribution were estimated from each dataset and used to parameterize the prior of the scLSD library size encoder.

For all analyses, we selected the 2,000–5,000 most highly variable genes (HVGs), retaining only genes with at least 20 detected counts. The exact number of HVGs was adjusted based on dataset size: for smaller datasets (bone marrow^11^ and dentate gyrus^42^), we used 2,000 HVGs, whereas for larger datasets we used 5,000 HVGs. The MERFISH human neurogenesis dataset^46^ was treated separately because it profiled a targeted panel of only 300 genes; accordingly, the available measured genes were used rather than applying the 2,000–5,000 HVG selection procedure.

### Axial Mesoderm Development Differential Expression Analysis

To identify transcriptomic differences between Notochord and Prechordal Plate populations, we performed differential expression analysis on the filtered single-cell RNA-seq data. To address the sparsity and zero-inflation inherent in the dataset, we employed the ZINB-WaVE framework^110^ to estimate observational weights (*K* = 2, *ϵ* = 1000), effectively downweighting technical zeros during downstream modeling. Differential expression was subsequently assessed using edgeR^111^, where a generalized linear model (GLM) was fitted to the weighted counts, and significance was determined via a likelihood ratio test (LRT) comparing the Notochord lineage against the Prechordal Plate lineage.

### Pathway Enrichment Analysis of Axial Mesoderm Development Perturbations

To assess pathway-level effects of large-scale gene perturbations in axial mesoderm development^49^, we analyzed the number of cells adopting a Notochord fate following perturbation of each gene. Perturbed Notochord counts were modeled using a Negative Binomial distribution, with Gamma and Beta priors placed on the dispersion and probability parameters, respectively. Model parameters were inferred via maximum a posteriori estimation using stochastic variational inference in Pyro.

Gene-wise significance scores were computed as tail probabilities under the fitted Negative Binomial model and transformed to − log *p*-values, which were used to rank genes by perturbation effect size. Ranked gene lists were then subjected to preranked gene set enrichment analysis using gseapy^112^ against Gene Ontology Biological Process^62^ and Reactome pathway^63^ databases (zebrafish annotations), with 1,000 permutations. For the top enriched pathways, running enrichment scores were extracted to characterize the cumulative contribution of individual genes across the ranking.

### Phenotype Enrichment Analysis of Mouse Cortex Gene Perturbations

For the mouse cortex development dataset^48^, we quantified the effect of each gene perturbation by counting the number of cells terminating in an inhibitory neuronal fate and computing the *log*_2_ fold change (*log*_2_*FC*) relative to an unperturbed baseline. Genes were ranked by inhibitory fate *log*_2_*FC* and the top 10 genes with positive and top 10 with negative *log*_2_*FC* were selected to form a perturbation gene set.

Phenotype enrichment analysis was performed using the Enrichr^113^ implementation in gseapy^112^, testing the selected gene set against a background universe comprising all genes present in the training dataset. Enrichment was evaluated using the MGI Mammalian Phenotype Level 4 (2024) library^71^, with significance assessed via Fisher’s exact test. Enrichr reports nominal p-values and FDR-adjusted p-values for each phenotype term, along with the contributing genes. The most significantly enriched phenotypes were retained for downstream interpretation and visualization.

### Zebrafish perturbation atlas analysis

Experimental cell-type abundance changes were calculated from the zebrafish perturbation atlas^72^ using embryos as biological replicates. Notochord and melanophore cells were identified from the broad cell-type annotations. To account for differences in cell recovery, each embryo’s cell-type count was divided by its total recovered cell count relative to the geometric mean across all embryos and rounded. Perturbed embryos were compared with stage-matched injected controls or designated mutant controls. Comparisons required at least one embryo per arm and at least 20 observed cells of the target type across the control arm. Embryos containing no cells of the target type were retained, with no minimum target-cell count imposed on the perturbed arm.

Abundance log_2_ fold-changes were calculated as 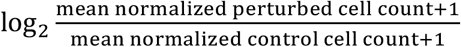. Complete losses, defined by zero observed target cells across the perturbed arm, were represented by a fixed log_2_ fold-change of –10.

We compared these experimental effects with scLSD-predicted changes in fate abundance for notochord at 24 hours post-fertilization and melanophore at 48 hours post-fertilization. Gene identifiers were harmonized, including the ta–tbxta alias, and comparisons were restricted to genes shared between the corresponding analyses; retained experimental log_2_ fold-changes mapping to the same gene were averaged. Compound (combinatorial) perturbations were excluded. Agreement was quantified by Pearson correlation across genes without filtering by experimental significance. The reported 95% confidence intervals used Fisher’s transformation^114^, followed by back-transformation to the correlation scale. Bootstrap distributions were calculated separately using 10,000 resamples of paired gene observations.

### Estimation of stemness-associated scores

Stemness-associated scores were estimated in the lung adenocarcinoma dataset^73^ using CytoTRACE2^74^ and complementary expression signatures. The mRNAsi score was calculated as the Pearson correlation between log-normalized single-cell expression and published stemness-model weights, followed by min–max normalization as described by Malta *et al*^*80*^. We additionally evaluated positively associated CancerSEA^75^ gene sets from lung adenocarcinoma^76^ and pan-cancer analyses, and the KEGG^115^, PluriNetWork^116^, hSCExplorer^117^ and Reactome^118^ signatures from StemChecker^77^. Gene-set activities were calculated using the univariate linear model method in decoupleR^78^. Human gene symbols were mapped to mouse orthologs using babelgene^79^.

## Data availability

The data sets analyzed in this paper are publicly available and published. The raw datasets of dentate gyrus neurogenesis^42^, erythroid gastrulation^43^ and hematopoiesis^11^ are available in scVelo package using *scvelo*.*datasets*.*dentategyrus, scvelo*.*datasets*.*gastrulation_erythroid, scvelo*.*datasets*.*bonemarrow*, respectively. The raw datasets of pancreas development^30^ and mouse cortex development^48^ are available in Gene Expression Omnibus repository under accession numbers GSE275562 and GSE249416 respectively. The raw dataset of lung adenocarcinoma progression^73^ is available on Zenodo (DOI 10.5281/zenodo.5847461). Human gonadal dataset^47^ is available in ArrayExpress with accession number E-MTAB-10551. Zebrafish perturbation atlas dataset^72^ is available in Gene Expression Omnibus under accession number GSE202639. The preprocessed human neurogenesis MERFISH dataset^46^ is available in Dryad (DOI 10.5061/dryad.2280gb612). All the preprocessed data used in this study are available via Zenodo (DOI 10.5281/zenodo.18331586).

## Code availability

The scLSD framework is packaged as sclsd and is available via pip. The source code is openly accessible at https://github.com/csglab/sclsd.

All notebooks and scripts required to reproduce the analyses and figures presented in this study are available at https://github.com/csglab/sclsd-manuscript.

## Author Contributions

AP, SH, HSN, and AE conceived the project. HSN and AE supervised the project and obtained funding. AP conducted model design, implementation, training and validation. SH ran baseline model comparisons. AP performed computational preprocessing of all single-cell datasets. AMN performed the downstream differential expression analysis. A.M. conducted single-cell analyses of stemness-based signatures. AP, AMN and HSN designed the data visualizations and figures. AS refactored the code. AP and AS packaged the code. AP, HSN, and AE wrote the first draft of the manuscript. All authors revised and approved the final manuscript.

## Acknowledgements

This work was supported by a grant from the Canadian Institutes of Health Research (CIHR) [PJT-173317] to HSN, by a grant jointly funded by Natural Sciences and Engineering Research Council of Canada (NSERC) [ALLRP 586837-23] and FRQNT [2024-NOVA-344140] to AE and HSN, and by a grant from NSERC [RGPIN-2019-04460] to AE. This work was also supported by resource allocations from Digital Research Alliance of Canada (alliancecan.ca) and Calcul Québec (www.calculquebec.ca) to HSN and AE. HSN holds a CIHR Canada Research Chair.

