## Supplementary Figures for "A Latent Space Thermodynamic Model of Cell Differentiation"

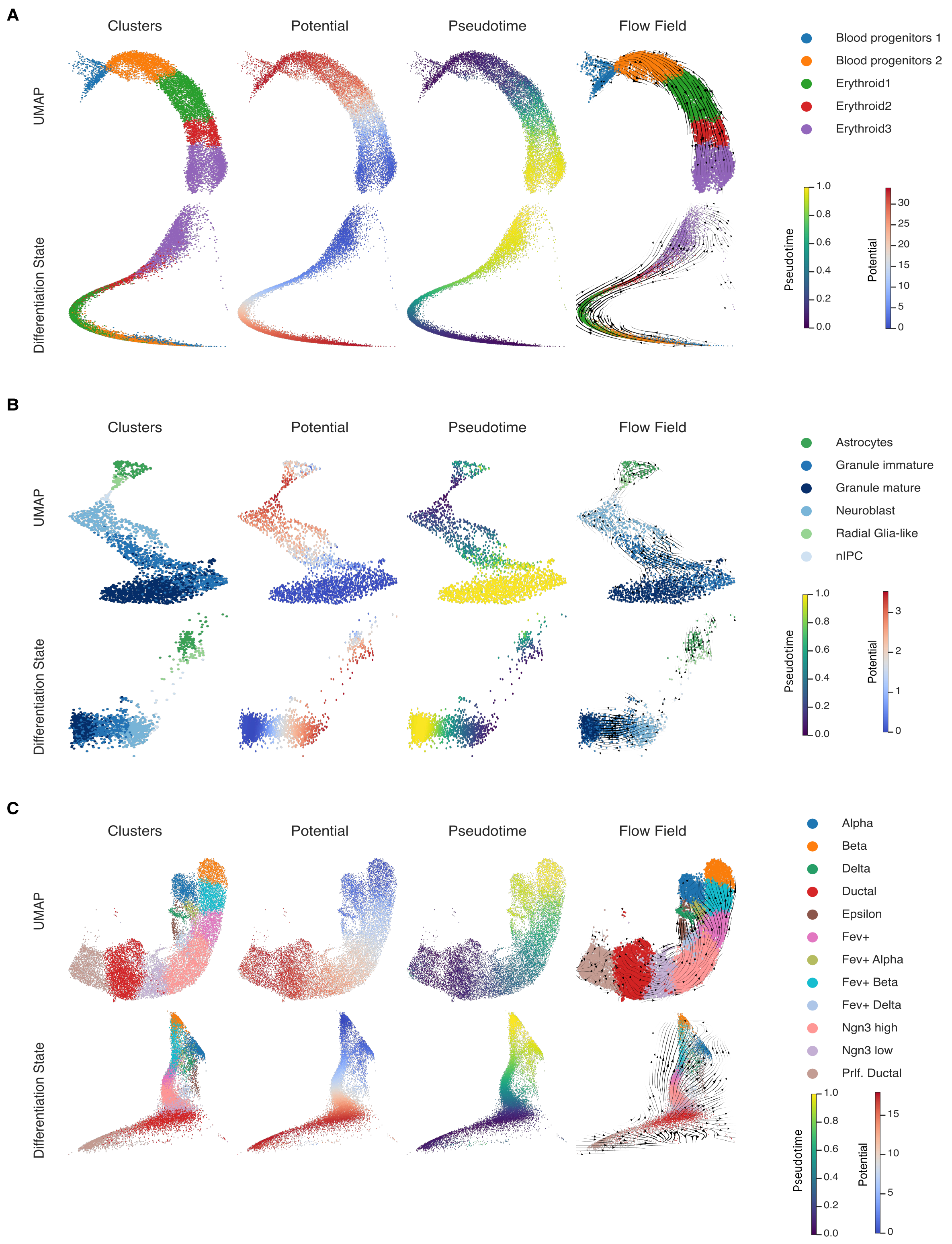


**Supplementary Figure 1. scLSD-inferred differentiation dynamics across multiple biological systems.**

For each dataset, the original UMAP embedding (top row) and the corresponding scLSD-inferred differentiation state (bottom row) are shown. Columns indicate cell-type clusters, inferred Waddington potential, pseudotime, and flow field, respectively. Colors in the cluster and flow-field plots indicate annotated cell types.

**(A)** Mouse erythroid gastrulation, showing blood progenitor populations and erythroid lineages.

**(B)** Dentate gyrus development, showing neuronal intermediate progenitor cells (nIPC), neuroblasts, granule neurons, radial glia-like cells, and astrocytes.

**(C)** Pancreatic endocrinogenesis, showing ductal cells, endocrine progenitors, and alpha, beta, delta, and epsilon lineages.

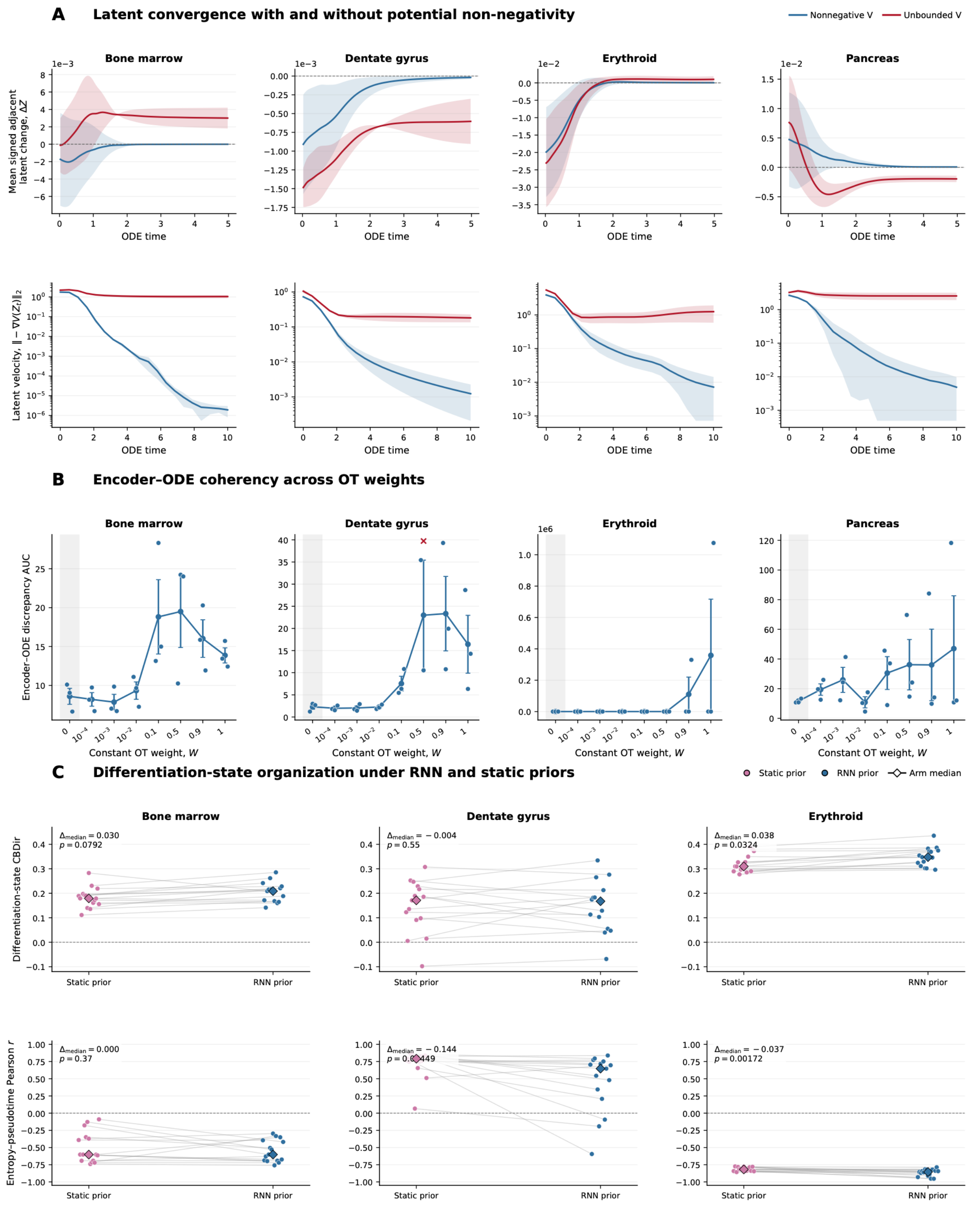


**Supplementary Figure 2. Ablations of potential non-negativity, optimal transport regularization, and the RNN prior.**

**(A)** Temporal evolution of the mean signed change between adjacent latent states (top) and latent velocity norm, $\|-\nabla V(z)\|_2$ (bottom), for models with nonnegative or unconstrained potential outputs. Columns show bone marrow, dentate gyrus, erythroid, and pancreas. Lines and shading indicate mean ± standard error of the mean (SEM) across three training seeds. Late-time velocity norms are lower in the nonnegative-potential models in all four datasets.

**(B)** Area under the variance-normalized encoder–ODE discrepancy curve across constant optimal transport (OT) weights, W; W = 0 removes the OT penalty. Discrepancy compares propagated latent states with their projected and re-encoded counterparts. Small points show individual seeds; connected circles and error bars show mean ± SEM. The lowest mean discrepancies occur at nonzero weights between $10^{-4}$ and $10^{-2}$, depending on the dataset.

**(C)** Differentiation-state cross-boundary direction correctness (CBDir; top) and Pearson correlation between entropy and the original input pseudotime (bottom) for RNN and static-zero priors. Points represent 15 training seeds per condition, gray lines connect matching seeds, and diamonds indicate medians. Annotations report the RNN-minus-static median difference and nominal one-sided Mann–Whitney U-test P values, with alternatives RNN > static for CBDir and RNN < static for entropy–pseudotime correlation.

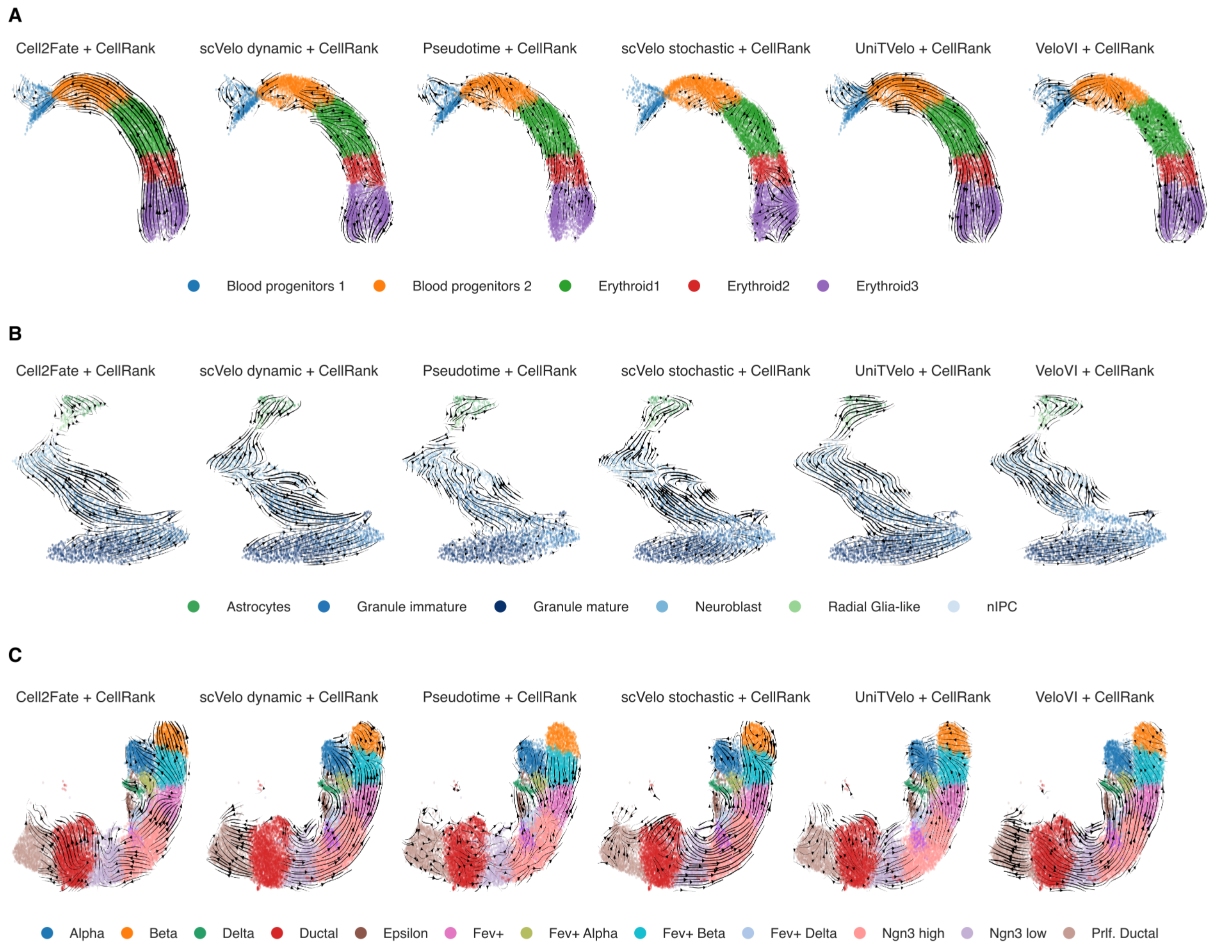


**Supplementary Figure 3. CellRank-inferred flow fields across upstream velocity and pseudotime methods.**

Cell-to-cell transition flow fields inferred by CellRank across three benchmark datasets. Columns correspond to CellRank initialized with Cell2Fate, scVelo dynamical, pseudotime, scVelo stochastic, UniTVelo, and VeloVI, respectively. Cells are colored by annotated cell types.

- **(A)** Mouse erythroid gastrulation.
- **(B)** Dentate gyrus development.
- **(C)** Pancreatic endocrinogenesis.

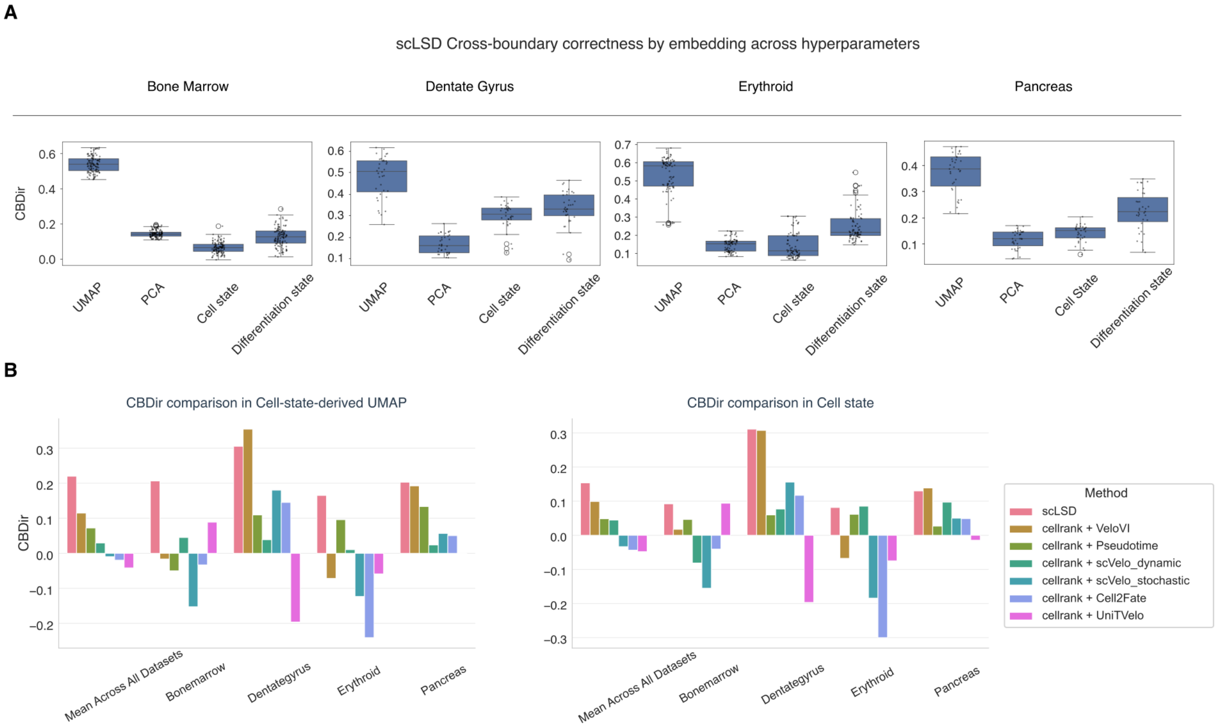


**Supplementary Figure 4. CBDir across model configurations and shared cell-state representations.**

**(A)** Distributions of scLSD CBDir scores across hyperparameter configurations in bone marrow, dentate gyrus, erythroid, and pancreas. Scores are evaluated in the original low-dimensional embedding, 50-dimensional PCA, 10-dimensional cell state, and two-dimensional differentiation state.

**(B)** CBDir for scLSD and CellRank initialized with the indicated RNA velocity or pseudotime methods, evaluated in a UMAP derived from the scLSD cell state (left) and directly in the scLSD cell state (right). Within each dataset, methods are evaluated using the same cell-state representation. Bars show dataset-specific scores and their unweighted mean across the four datasets. scLSD has the highest mean CBDir in both representations.

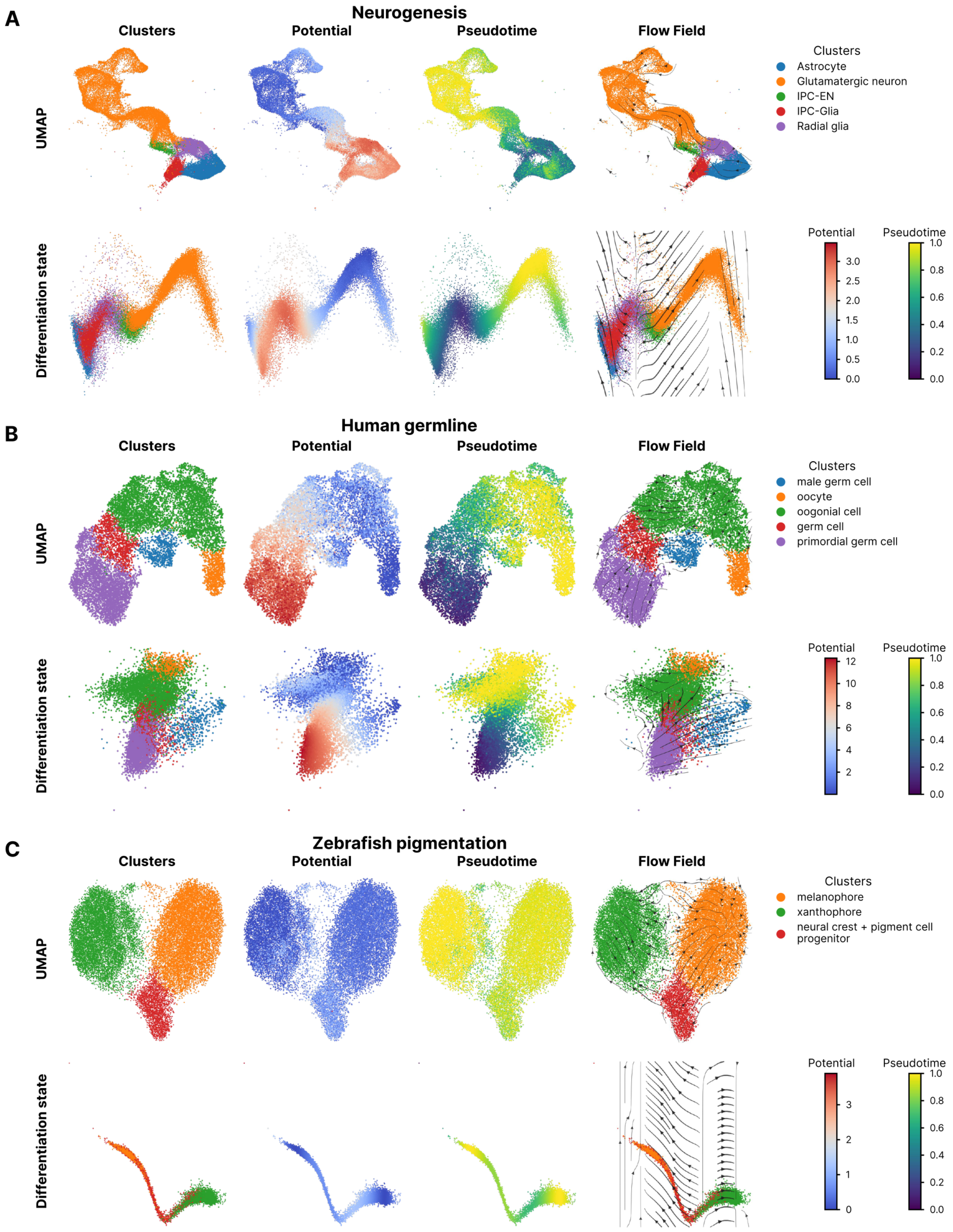


**Supplementary Figure 5. scLSD-inferred landscapes and flow fields in neurogenesis, human germline development, and zebrafish pigmentation.**

**(A-C)** Neurogenesis (A), human germline development (B), and zebrafish pigmentation (C). For each dataset, the upper row shows the UMAP embedding and the lower row shows the scLSD-inferred differentiation state. Columns display annotated clusters, potential, pseudotime, and flow field, respectively. Cluster colors are retained in the flow-field plots; continuous color scales indicate potential and pseudotime. The datasets include neuronal and glial populations in A, male and female germline populations in B, and neural crest/pigment progenitors, melanophores, and xanthophores in C.

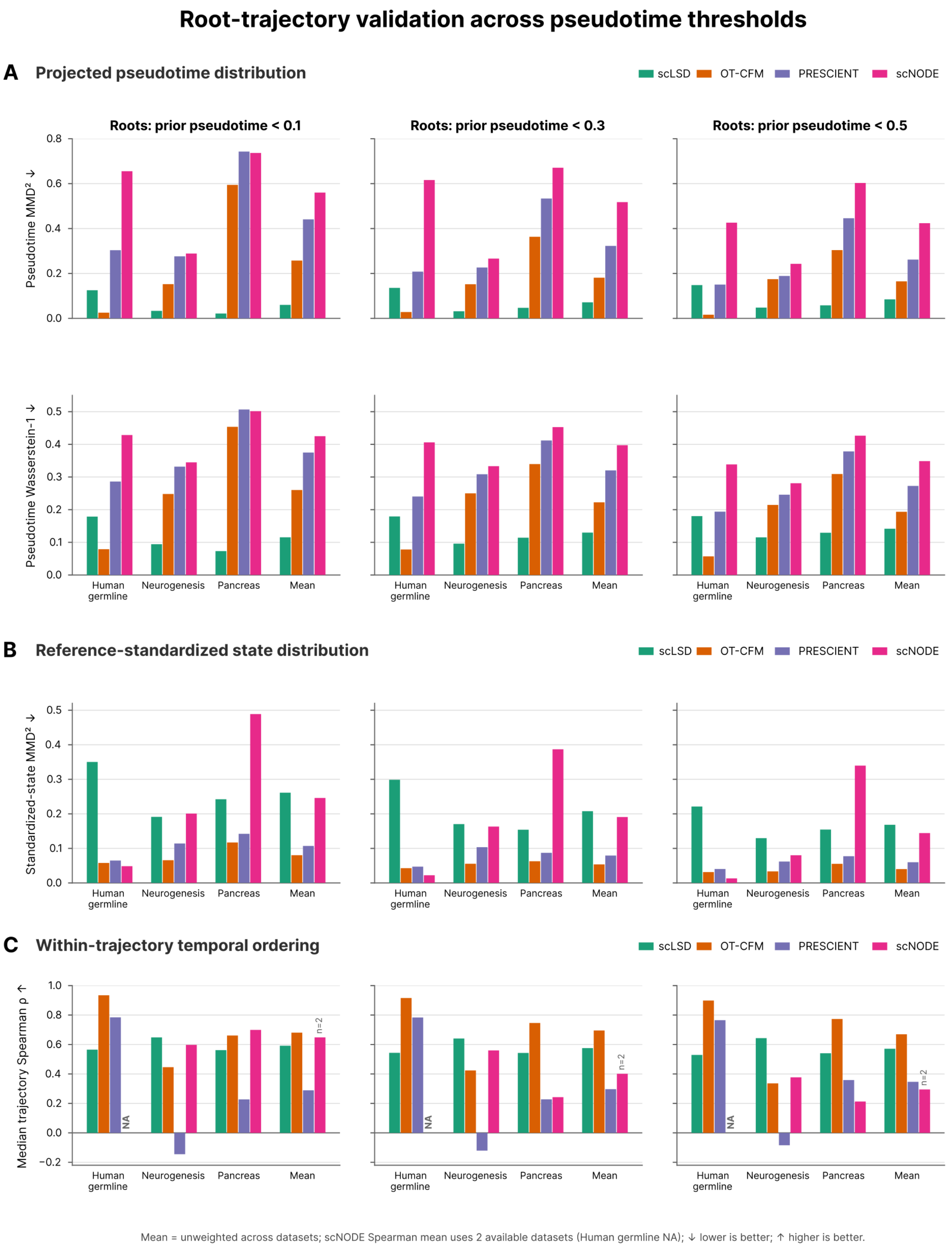


**Supplementary Figure 6. Root-trajectory distributions and temporal ordering across pseudotime thresholds.**

Comparison of scLSD, OT-CFM, PRESCIENT, and scNODE in human germline, neurogenesis, and pancreas. Columns use root cells with original input pseudotime below 0.1, 0.3, or 0.5, respectively. The final bar group in each subplot shows the unweighted mean of available dataset scores.

- **(A)** Squared maximum mean discrepancy ($MMD^2$; top) and Wasserstein-1 distance (bottom) between the pseudotime distribution of propagated trajectory points and that of observed cells. Trajectory-point pseudotime is assigned from the nearest observed cell. Lower values indicate smaller distributional differences.
- **(B)** $MMD^2$ between propagated and observed cell-state distributions after standardization using the observed-cell mean and standard deviation in each method's state space.
- **(C)** Median within-trajectory Spearman correlation between model integration time and projected pseudotime. NA denotes an unavailable value; the scNODE mean in C uses neurogenesis and pancreas only (n = 2), excluding the unavailable human germline value.

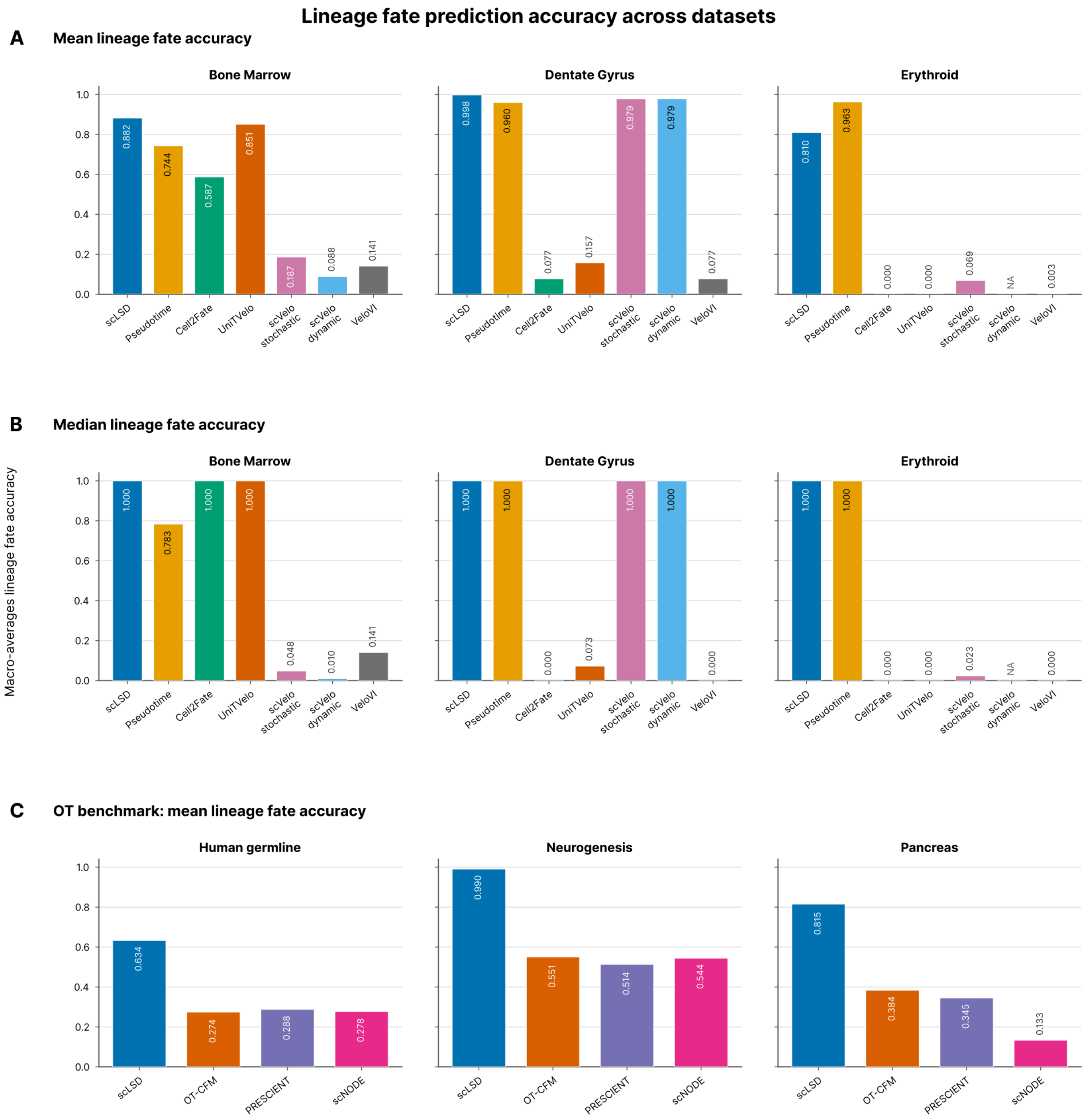


**Supplementary Figure 7. Lineage fate prediction accuracy across datasets and methods.**

- **(A)** Mean probability assigned to lineage-compatible fates across eligible cells in bone marrow, dentate gyrus, and erythroid. scLSD is compared with CellRank initialized using pseudotime, Cell2Fate, UniTVelo, scVelo stochastic, scVelo dynamical, or VeloVI.
- **(B)** Median lineage-compatible fate probability across cells for the same methods and datasets.
- **(C)** Endpoint fate classification accuracy for scLSD, OT-CFM, PRESCIENT, and scNODE in human germline, neurogenesis, and pancreas, macro-averaged across source lineages. A prediction is counted as correct when its assigned endpoint cluster belongs to the source

cell's allowed fate set. scLSD has the highest displayed accuracy in each of these three datasets. Numbers indicate bar values; NA denotes an unavailable result.

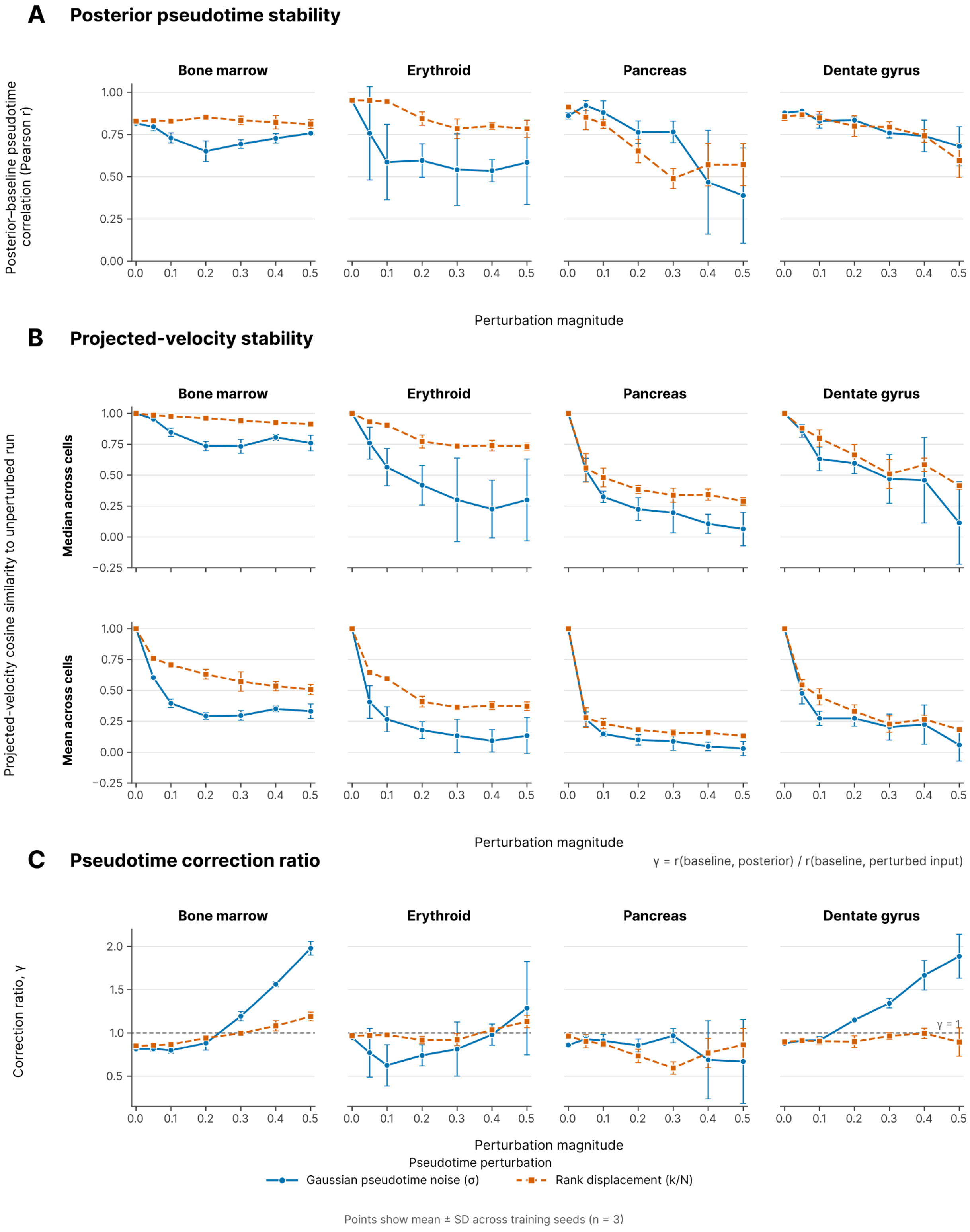


**Supplementary Figure 8. Sensitivity of scLSD outputs to perturbations of input pseudotime.**

Gaussian pseudotime noise (standard deviation σ) and rank displacement (relative magnitude $\frac{k}{N}$) are evaluated in bone marrow, erythroid, pancreas, and dentate gyrus. Points and error bars show mean ± standard deviation across three training seeds.

**(A)** Pearson correlation between scLSD posterior pseudotime after training on perturbed inputs and the original, unperturbed input pseudotime.

**(B)** Cell-wise cosine similarity between projected velocities from perturbed and unperturbed models with the same training seed. The upper and lower rows summarize the median and mean across cells, respectively, before averaging across seeds. Similarities generally decrease with increasing perturbation magnitude.

**(C)** Pseudotime correction ratio, $\gamma = \frac{\mathrm{r(baseline,posterior)}}{\mathrm{r(baseline,perturbed\ input)}}$, where baseline denotes the original input pseudotime and r is Pearson correlation. The dashed line marks $\gamma = 1$. At larger Gaussian noise levels, $\gamma$ exceeds one in bone marrow and dentate gyrus, whereas it remains below one in pancreas.

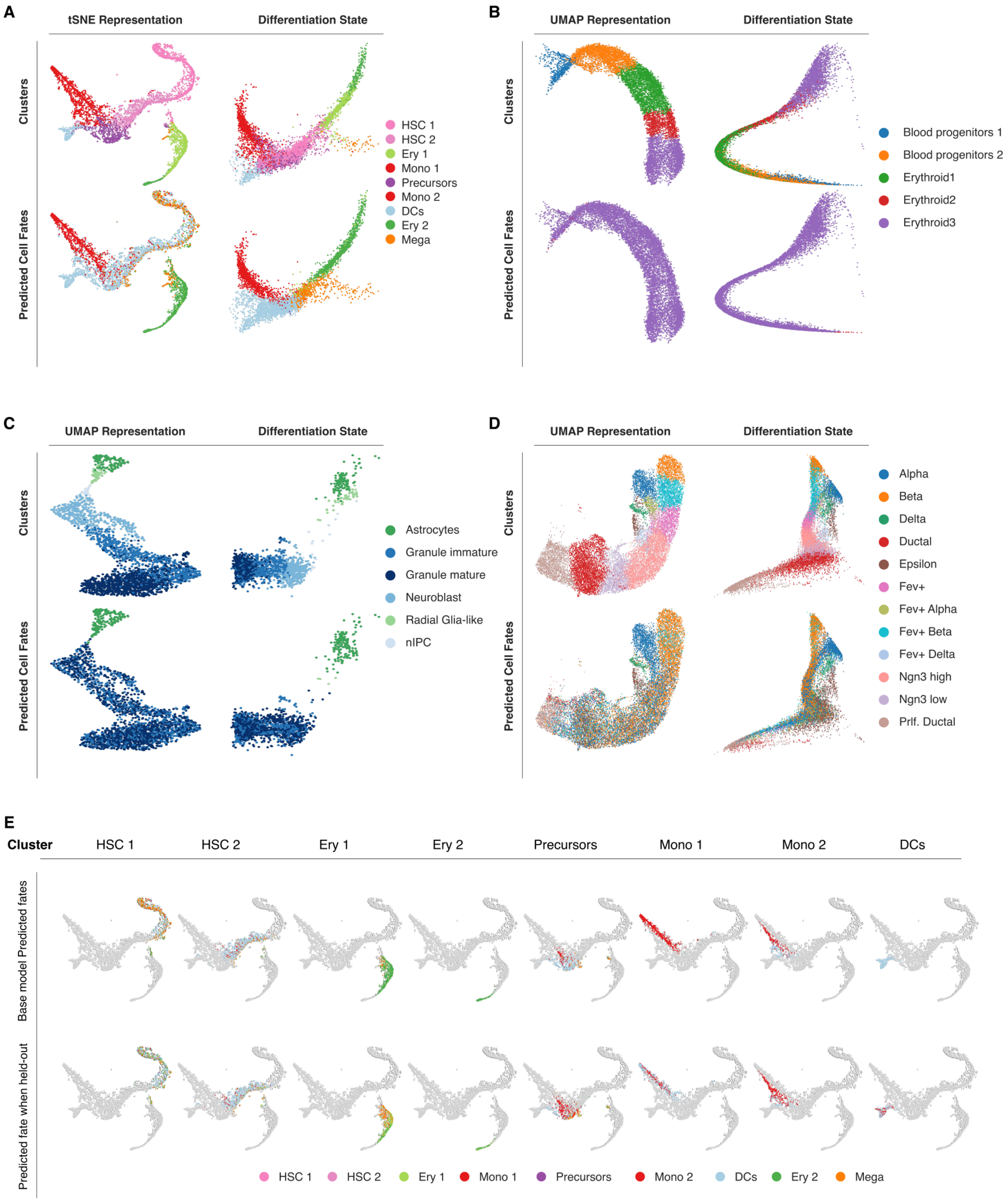


**Supplementary Figure 9. Cell-fate assignments across datasets and held-out bone marrow populations.**

**(A-D)** Annotated cell-type clusters and corresponding scLSD-predicted cell fates in bone marrow hematopoiesis (A), erythroid gastrulation (B), dentate gyrus (C), and pancreatic endocrinogenesis (D). For each dataset, the first column shows the original low-dimensional embedding (t-SNE or UMAP), while the second shows the scLSD-inferred differentiation state. Rows show annotated clusters and predicted fates, respectively.

**(E)** Leave-out-cluster evaluation in bone marrow. Each column shows fate assignments for the indicated population when included in training (base model; top) or excluded from training (held-out model; bottom). Cells in the evaluated population are colored by predicted fate; other cells are shown in gray.

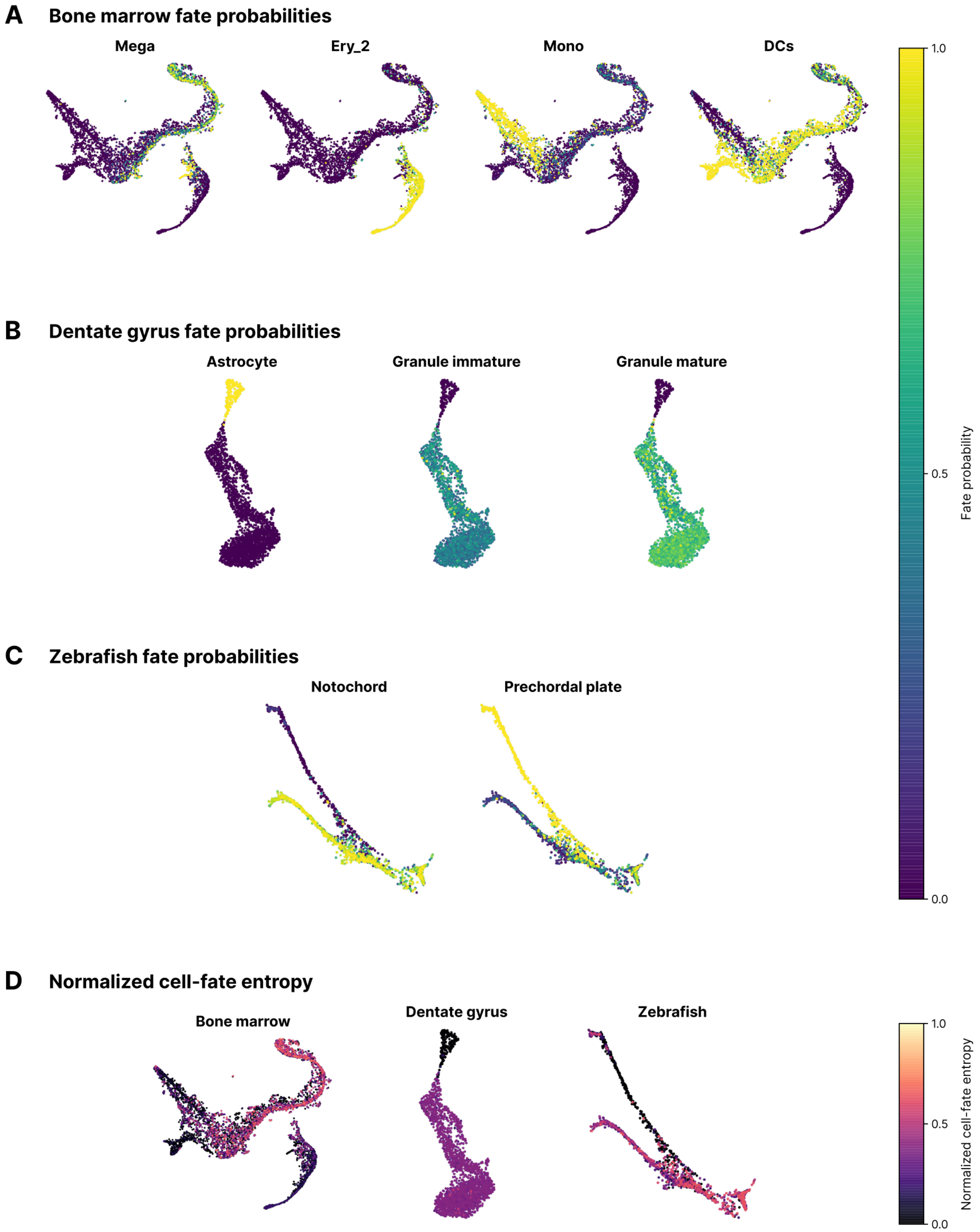


**Supplementary Figure 10. Posterior-sampled cell-fate probabilities and fate entropy.**

Fate probabilities are estimated by sampling latent initial states from each cell's encoder posterior, propagating them through the learned dynamics, and assigning endpoint cluster labels.

- **(A)** Bone marrow t-SNE embedding colored by the probabilities of Mega, Ery_2, Mono (Mono_1), and DC fates.
- **(B)** Dentate gyrus UMAP colored by astrocyte, immature granule, and mature granule fate probabilities.
- **(C)** Zebrafish axial mesoderm force-directed embedding colored by notochord and prechordal plate fate probabilities.
- (D) Normalized cell-fate entropy in the three datasets, calculated as the Shannon entropy of the full endpoint-label probability distribution divided by the logarithm of the number of possible labels.

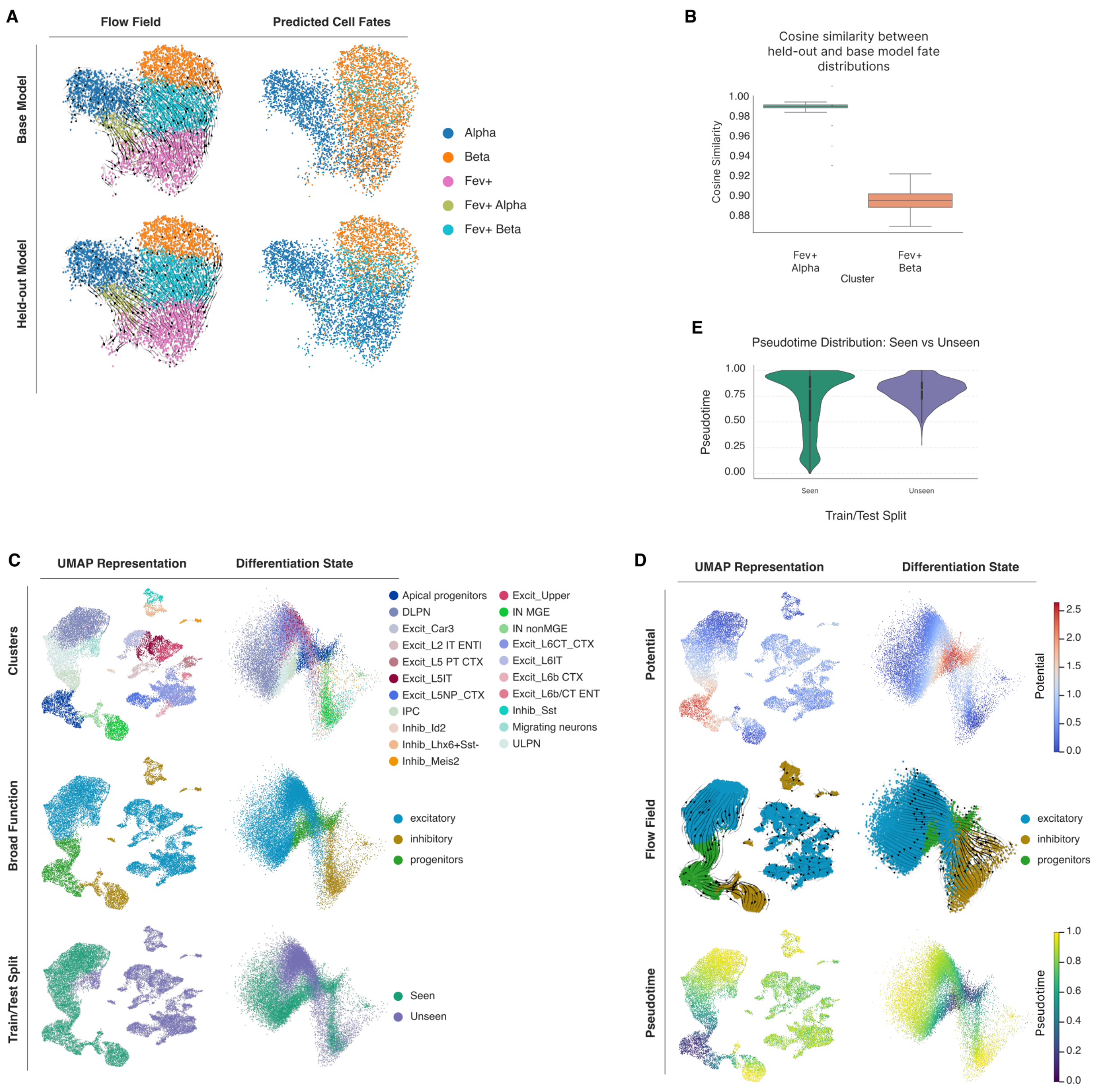


**Supplementary Figure 11. Predicted dynamics and fates for held-out pancreatic and cortical cell populations.**

**(A)** Flow field and predicted cell fates for the base model (top) and a held-out model (bottom) in pancreatic endocrinogenesis. Fev$^+$ Alpha and Fev$^+$ Beta populations, descendants of the Fev$^+$ progenitor cluster, are jointly excluded during held-out-model training.

**(B)** Cosine similarity between fate distributions predicted by the base and held-out models for Fev$^+$ Alpha and Fev$^+$ Beta, with values centered near 0.99 and 0.90, respectively.

**(C)** UMAP and scLSD-inferred differentiation state of the mouse cortex dataset. Rows show annotated fine-grained cell types, broad functional classes (progenitors, excitatory neurons, and inhibitory neurons), and the train/test split. The training dataset contains progenitors and neuronal populations; the held-out dataset contains mature neuronal subtypes.

**(D)** scLSD-inferred potential, flow field, and pseudotime in the mouse cortex dataset, shown on UMAP (left) and differentiation state (right).

**(E)** Violin plots of pseudotime for seen and unseen cortical cells. Unseen cells are concentrated at later pseudotime values.

A

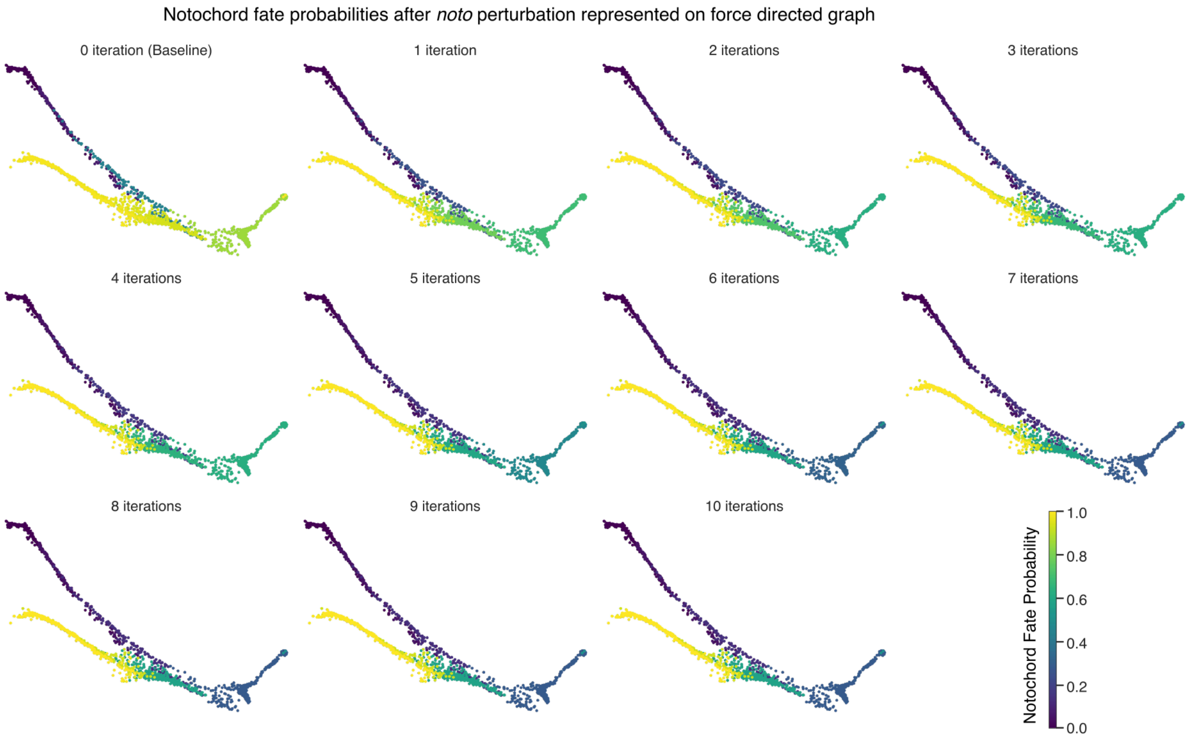
Notochord fate probabilities after *noto* perturbation represented on force directed graph
0 iteration (Baseline)
1 iteration
2 iterations
3 iterations
4 iterations
5 iterations
6 iterations
7 iterations
8 iterations
9 iterations
10 iterations
Notochord Fate Probability
1.0
0.8
0.6
0.4
0.2
0.0

B

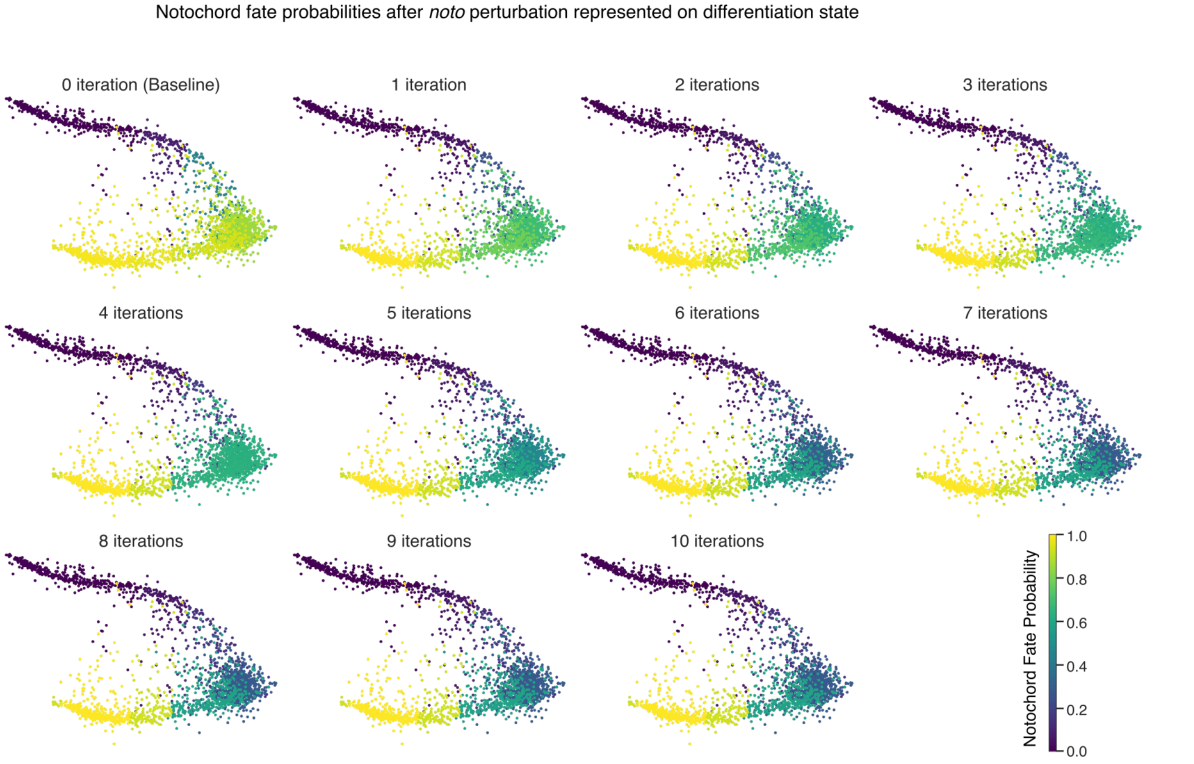
Notochord fate probabilities after *noto* perturbation represented on differentiation state
0 iteration (Baseline)
1 iteration
2 iterations
3 iterations
4 iterations
5 iterations
6 iterations
7 iterations
8 iterations
9 iterations
10 iterations
Notochord Fate Probability
1.0
0.8
0.6
0.4
0.2
0.0

**Supplementary Figure 12. Notochord fate probabilities following iterative in silico noto perturbation.**

Notochord fate probabilities following repeated in silico knockout of *noto*. Perturbations are applied by setting *noto* expression to zero at successive steps of trajectory propagation. Subplots show the unperturbed baseline (0 iterations) and increasing numbers of perturbation iterations, up to 10.

**(A)** Fate probabilities on the force-directed representation of zebrafish axial mesoderm development.

**(B)** The same probabilities on the scLSD-inferred differentiation state. Notochord fate probabilities decrease in part of the cell population as the number of perturbation iterations increases.

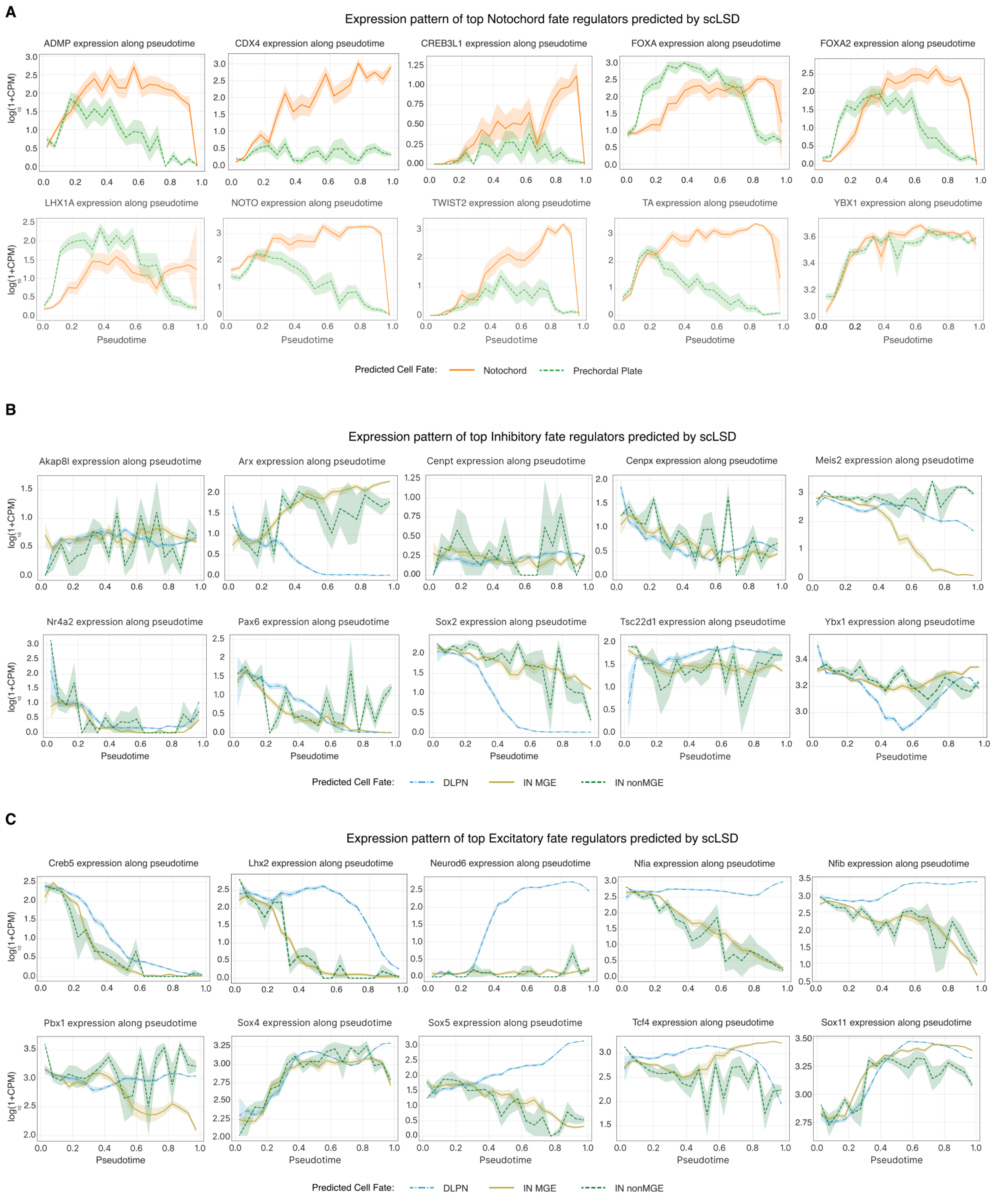


**Supplementary Figure 13. Expression patterns of genes ranked by in silico lineage perturbations.**

Expression patterns of the highest-ranked genes from the scLSD in silico perturbation analysis, stratified by predicted cell fate and plotted along scLSD pseudotime. Expression is shown as $\log_{10}(1 + \mathrm{CPM})$, where CPM denotes counts per million.

- **(A)** Ten top-ranked genes associated with notochord fate in zebrafish axial mesoderm, comparing notochord and prechordal plate trajectories.
- **(B)** Ten top-ranked genes associated with inhibitory neuronal fate in mouse cortex, shown across DLPN (excitatory), IN-MGE, and IN-non-MGE (inhibitory) fate groups.
- **(C)** Ten top-ranked genes associated with excitatory neuronal fate in the same dataset and fate groups.

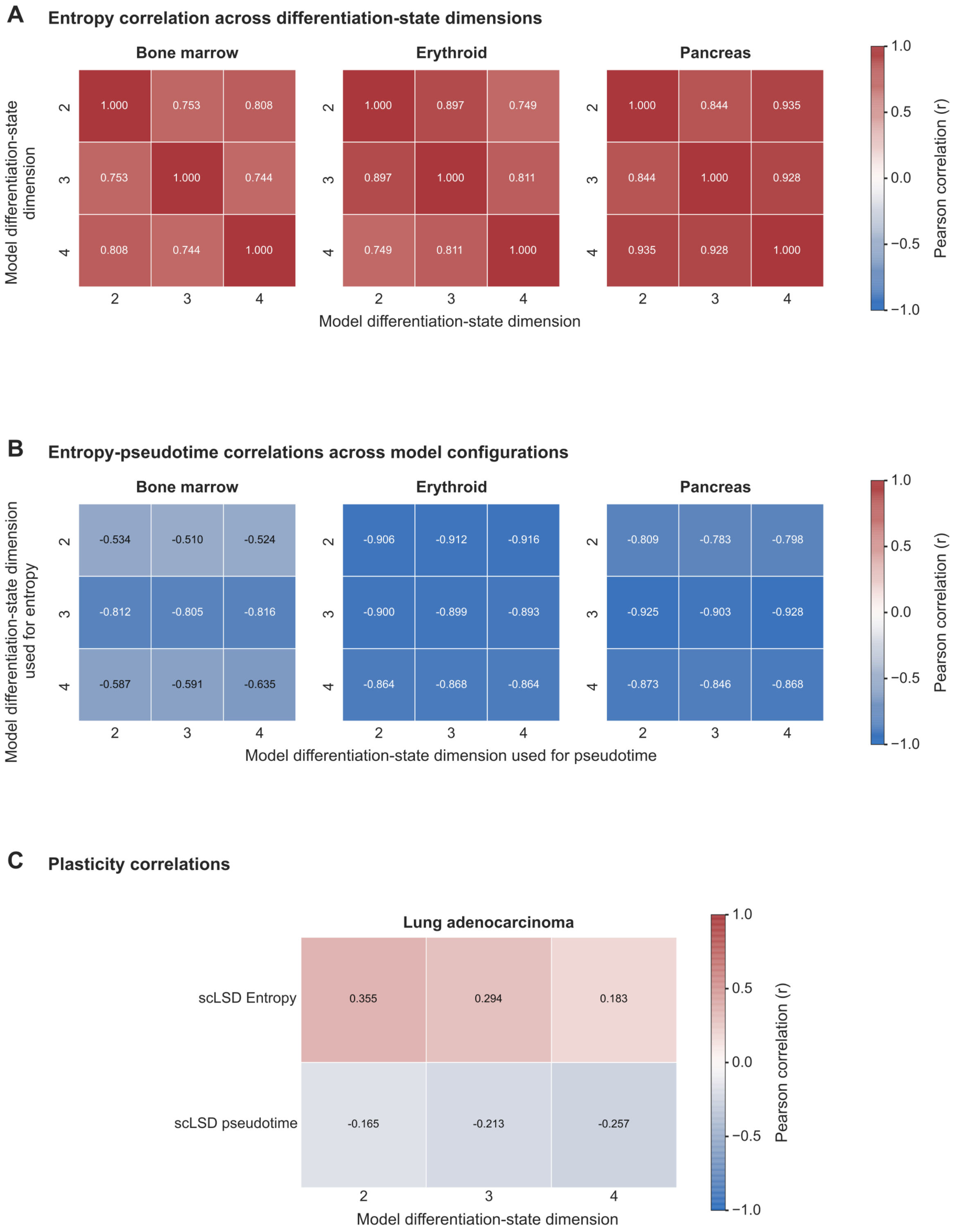


**Supplementary Figure 14. Entropy and pseudotime associations across differentiation-state dimensions.**

Models with differentiation-state dimensions of 2, 3, and 4 are compared in bone marrow, erythroid, pancreas (**A-B**), and lung adenocarcinoma (**C**). Heatmap colors and annotations indicate Pearson correlations.

- **(A)** Pairwise correlations of cell-level entropy values between model configurations.
- **(B)** Correlations between entropy and pseudotime across configurations. Rows identify the model dimension used to obtain entropy, and columns identify the model dimension used to obtain pseudotime; both outputs are scalar cell-level quantities.
- **(C)** Correlations of entropy and scLSD pseudotime with lineage-tracing-derived effective plasticity in lung adenocarcinoma. Entropy correlations are positive and pseudotime correlations are negative at each displayed dimension. The entropy correlation decreases from 0.355 at dimension 2 to 0.183 at dimension 4.

A

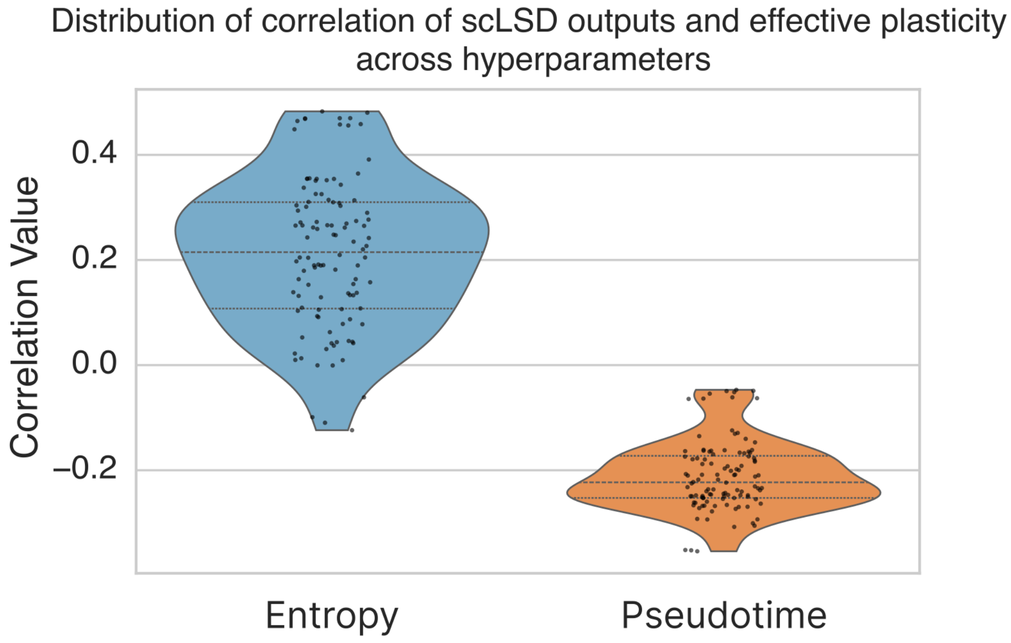


**Supplementary Figure 15. Entropy–plasticity and pseudotime–plasticity correlations across model configurations.**

Distributions of correlations between scLSD-inferred entropy or pseudotime and effective plasticity across hyperparameter configurations in lung adenocarcinoma. Points represent individual model configurations; violins show the distributions, with internal lines marking the median and quartiles. Entropy correlations are predominantly positive, whereas pseudotime correlations are negative across the displayed configurations.

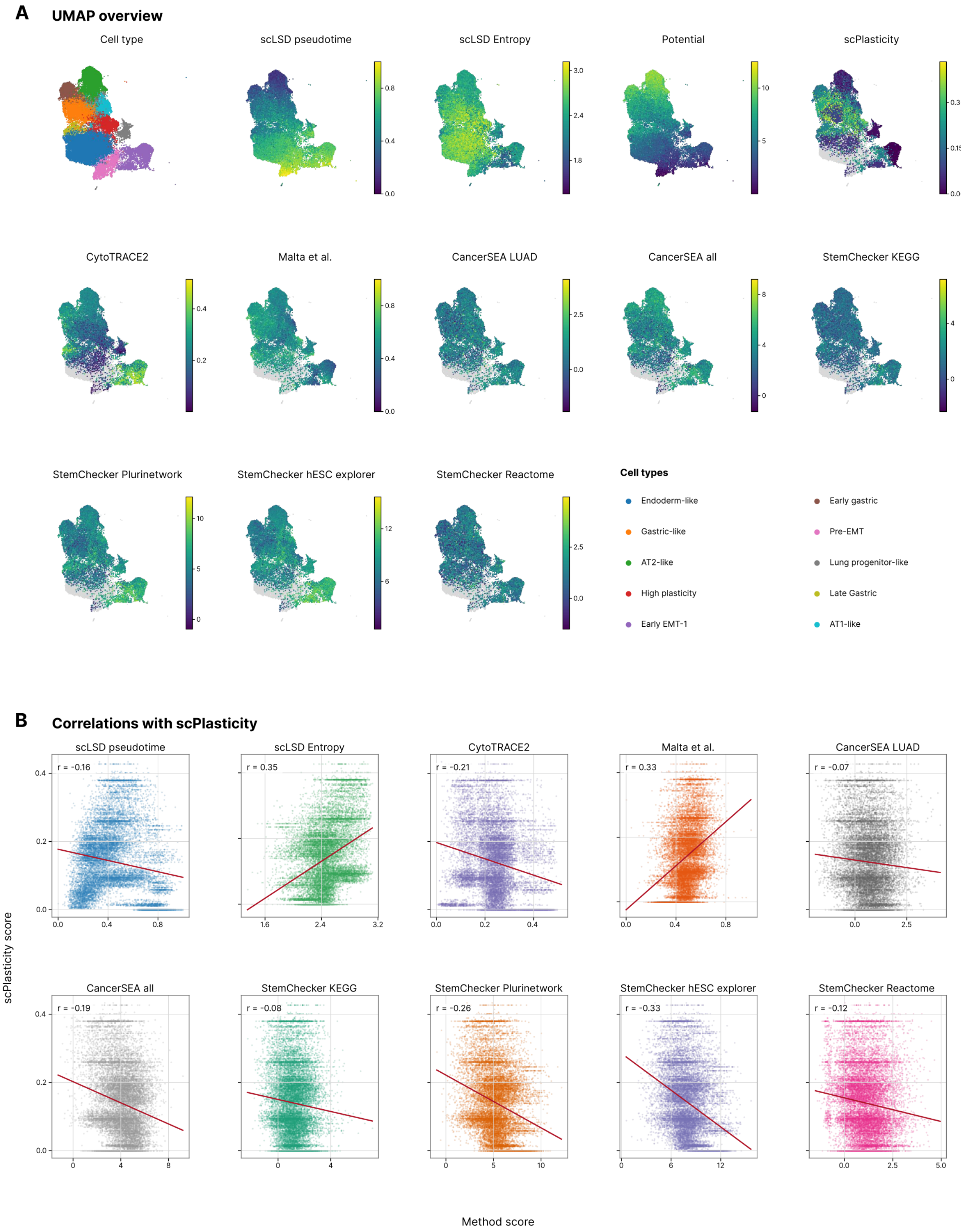


**Supplementary Figure 16. scLSD outputs and stemness-associated scores in lung adenocarcinoma.**

**(A)** UMAP embeddings colored by cell type, scLSD pseudotime, developmental entropy, potential, lineage-tracing-derived effective plasticity (scPlasticity), and the indicated stemness-associated scores. Scores include CytoTRACE2, Malta et al., CancerSEA lung adenocarcinoma (LUAD) and pan-cancer signatures, and StemChecker gene sets. Each continuous measure uses its own color scale.

(B) Scatterplots comparing each indicated score or scLSD output with scPlasticity. Each point represents a cell; red lines show linear fits, and annotations give Pearson correlations. Entropy and the Malta score have positive correlations (r = 0.35 and 0.33, respectively), while the remaining displayed measures have negative correlations.

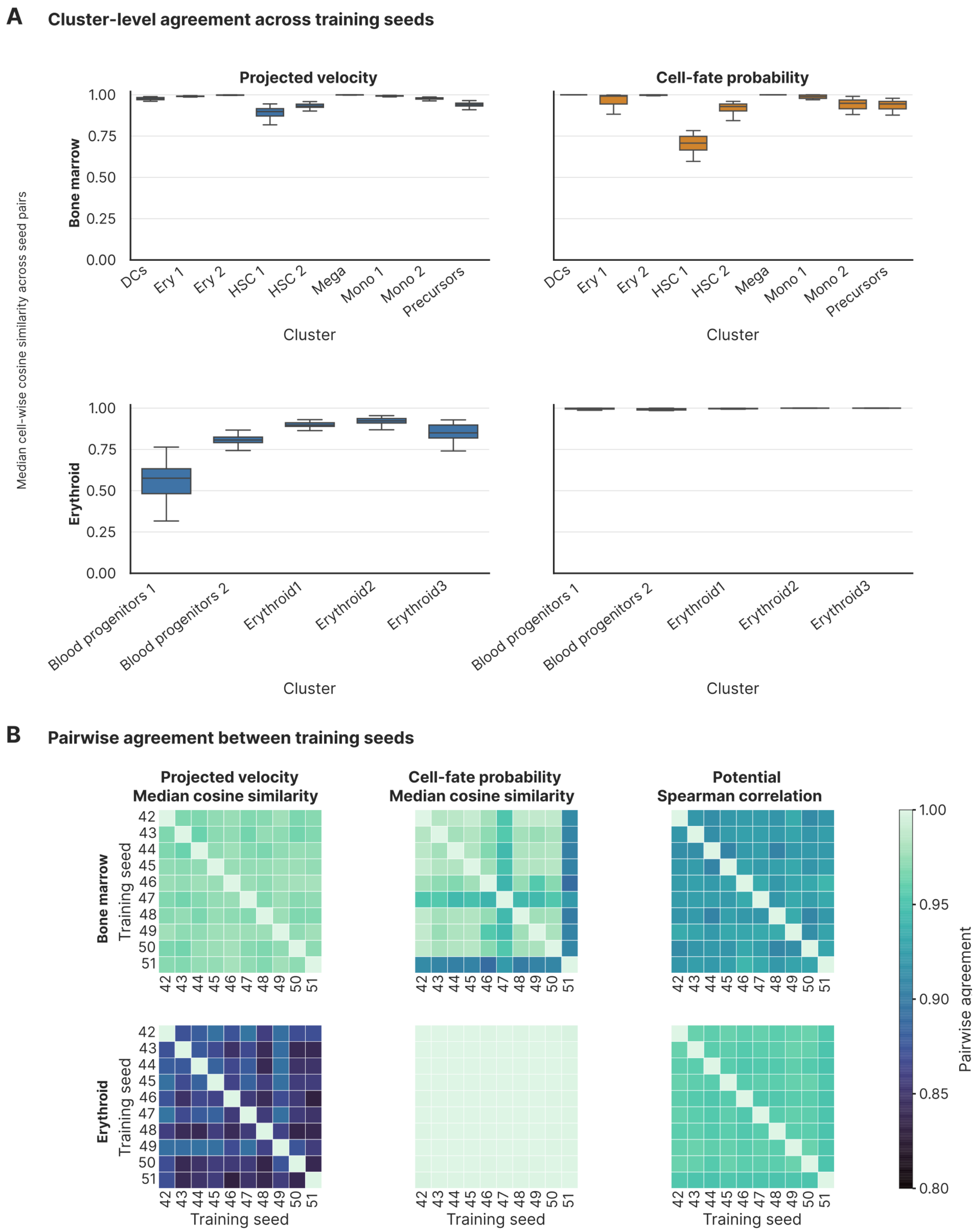


**Supplementary Figure 17. Agreement of projected velocities, fate probabilities, and potentials across training seeds.**

Ten models per dataset are trained with fixed hyperparameters and random seeds 42–51. Bone marrow and erythroid results are shown in the upper and lower rows of each panel, respectively.

- **(A)** Cluster-level agreement of projected velocity vectors (left) and cell-fate probability vectors (right). Each box summarizes 45 seed-pair values; each value is the median cell-wise cosine similarity within the indicated cluster for one seed pair. Velocities are projected onto the shared t-SNE embedding for bone marrow and UMAP for erythroid. Projected-velocity similarity is lower in blood progenitors 1 than in the erythroid clusters, whereas fate-probability similarities are close to one throughout the erythroid dataset.
- **(B)** Seed-by-seed matrices of median cell-wise cosine similarity for projected velocity and fate probabilities, and Spearman correlation of cell-level potential values. Each matrix entry compares the indicated pair of training seeds across cells; colors use a shared agreement scale from 0.8 to 1.0.
