## Supplementary Notes for "A Latent Space Thermodynamic Model of Cell Differentiation"

### Thermodynamics-inspired motivation and variational uncertainty

Stochastic thermodynamics provides a theoretical framework for extending the laws of thermodynamics to small, fluctuating systems[1]. Mathematically, the natural description of such noisy dynamics is through stochastic differential equations (SDEs). When a system is coupled to a heat bath, the forces acting on it decompose into deterministic components (e.g., conservative or active driving forces) and random components generated by thermal fluctuations. Similarly, in developmental biology, cells traverse Waddington's epigenetic landscape; in this picture, the gradient of the landscape generates the deterministic drift term that drives differentiation.

Formally, let the cell state be represented as a vector $z_t \in \mathbb{R}^D$, and the Waddington potential, the term we use to refer to quantitatively measure the "height" of each point in Waddington landscape, to be defined as function $V: \mathbb{R}^D \rightarrow \mathbb{R}_+$. Hence, we write down the equation of cell state evolution as the following SDE

$$dz_t = -\nabla V(z)\, dt + \Sigma(z,t)\, dW_t, \tag{1}$$

where $\Sigma(z,t)$ is a $D \times D$ matrix that denotes the noise amplitude and $-\nabla V(z)$ is the deterministic drift pulling cells toward local minima of $V(z)$. Thermodynamically, the noise amplitude can be seen as a local "temperature": where $\Sigma$ is large, fluctuations are stronger, and cells can more easily escape shallow wells or differentiation barriers.

the probability distribution associated with the stochastic process described in equation (1), $p(z,t)$, is then derived from the associated Fokker-Planck equation

$$\frac{\partial}{\partial t} p(z,t) = \nabla.\left(p(z,t)\, \nabla V(z)\right) + \sum_{i,j=1}^{D} \frac{\partial^2}{\partial z_i \partial z_j}\left(p(z,t)\, \Gamma_{ij}(z,t)\right) \tag{2}$$

where $\Gamma = \frac{1}{2}\Sigma(z,t)\,\Sigma(z,t)^T$. Then the entropy associated with cell state evolution is defined as the Shannon entropy, $S = -\int dz\, p(z,t) \log p(z,t)$.

Since no closed-form solutions for equation (1) exists, Gaussian moment closure[2] can be used to approximate the solutions of equation (2) as

$$p(z,t) = N(\mu_z, \sigma_z), \tag{3}$$

where first order closure implies

$$\frac{d\mu_z}{dt} = -\nabla V(z) \tag{4}$$

$$\frac{d\sigma_z}{dt} = -\nabla^2 V(\mu_z)\sigma_z - \sigma_z \nabla^2 V(\mu_z) + \Sigma(\mu_z,t)\, \Sigma(\mu_z,t)^T \tag{5}$$

In practice, neither the Waddington potential $V(z)$ nor the noise amplitude term $\Sigma(z,t)$ is known a priori. Instead, they must be inferred from high-dimensional single-cell data, where the observable cell states are noisy, sparse samples drawn from the underlying stochastic process. The Gaussian moment closure approximation in Eq. (3) motivates a parametric representation of both the drift and diffusion terms. To operationalize this idea, we parameterize the potential $V(z)$ with a neural network whose gradients define ordinary differential equation (4). Furthermore, we avoid solving equation (5) and directly parameterize $\sigma_z$ through a neural network as measure of local uncertainty.

### Theorems

*Theorem 1:* Let $\Omega \subset \mathbb{R}^D$ be open and connected and assume the VAE model class is closed under smooth invertible reparameterizations of the latent variable. Define the ELBO by

$\mathcal{E} = \mathbb{E}_{q(Z|x)}[\log p(x|z)] - KL\big(q(z|x) \| p(z)\big).$

Then the following holds:

*i)* Let $h: \Omega \to \tilde{\Omega}, \tilde{z} = h(z)$. Transform the encoder and prior by the pushforward $\tilde{q}(\tilde{z}|x) = h_{\#}q(z|x), \tilde{p}(\tilde{z}) = h_{\#}p(z)$ and define the transformed decoder $\tilde{p}(x|\tilde{z}) = p(x|h^{-1}(\tilde{z}))$.
Then $\mathcal{E} = \tilde{\mathcal{E}}$

*ii)* Suppose that the latent state satisfies $\frac{dz}{dt} = -\nabla V(z)$ for some $V \in C^2(\Omega)$ and let $\flat$ denote the Euclidean metric duality between vector fields and 1-forms. Then the diffeomorphism $h$ gives an equivalent representation within the same class of Euclidean gradient-flow models if and only if $\beta_h := \big(h_*(-\nabla V)\big)^\flat$ is exact on $h(\Omega)$.

*iii)* Let $\dim \Omega \geq 2$, $\nabla V \not\equiv 0$ and define $\mathcal{G}_V := \Big\{h \in \mathrm{Diff}(\Omega): \big(h_*(-\nabla V)\big)^\flat$ is exact on $h(\Omega)\Big\}$. Then $\mathcal{G}_V$ is a strict subset of $\mathrm{Diff}(\Omega)$.

*Proof:* By change of variables $\tilde{q}(\tilde{z}|x) = q(z|x)\,|\det Dh(z)|^{-1}$ and $\tilde{p}(\tilde{z}) = p(z)|\det Dh(z)|^{-1}$. Hence

$\frac{\tilde{q}(\tilde{z}|x)}{\tilde{p}(\tilde{z})} = \frac{q(z|x)}{p(z)}$. This means $KL\big(q(z|x) \| p(z)\big) = KL\big(\tilde{q}(\tilde{z}|x) \| \tilde{p}(\tilde{z})\big)$. Similarly, by definition of pushforward $\int f(\tilde{z}) d(h_{\#}q)(\tilde{z}) = \int f\big(h(z)\big) dq(z)$ for an arbitrary integrable function $f$. Letting $f(\tilde{z}) = \log \tilde{p}(x|\tilde{z})$ yields $\mathbb{E}_{q(Z|x)}[\log p(x|z)] = \mathbb{E}_{\tilde{q}(\tilde{z}|x)}[\log \tilde{p}(x|\tilde{z})]$ which proves the part (*i*) result.

For the second part let $F := -\nabla V$ denote the latent vector field, so that $\frac{dz}{dt} = F(z)$. Under diffeomorphism $h: \Omega \to \tilde{\Omega}$ the transformed trajectory $\tilde{z} = h(z)$ satisfies $\frac{d\tilde{z}}{dt} = Dh(z)F(z) = (h_*F)(\tilde{z})$. Therefore $h$ gives an equivalent representation within the class of Euclidean gradient flows if and only if there exist a potential $\tilde{V}: \tilde{\Omega} \to \mathbb{R}$ such that $h_* F = -\nabla \tilde{V}$. In other words, defining the 1-form $\beta_h := \left(h_*(-\nabla V)\right)^{\flat}$, the transformed dynamics are a Euclidean gradient flow if and only if $\beta_h = -d\tilde{V}$.

For part (*iii*) it suffices to exhibit diffeomorphism $h$ such that $\beta_h$is not exact. There are two distinct mechanisms. First $h$ may destroy the gradient structure locally, meaning $d\beta_h \neq 0$. For example let $V(x,y) = \frac{1}{2}(x^2 + y^2)$ and $h(x,y) = (x, y + \epsilon x^2)$. In transformed coordinated $(u,v)$,

$\beta_h = -u\, du - (v + \epsilon u^2) dv$ , $d\beta_h = -2\epsilon u\, du \wedge dv \neq 0$. Hence $h \notin \mathcal{G}_V$.

A second topological obstruction can occur when the manifold has non-trivial first de Rham cohomology group. For example, let $\Omega = \{(r,\theta): 1 < r < 2\}, V(r,\theta) = r$ and $h(r,\theta) = \left(r, \theta - \frac{c}{r}\right), c \neq 0$. Then

$\beta_h = -dr - c\, d\theta$. While $d\beta_h = 0$, one can see that $\oint \beta_h = -2\pi c$, and hence $\beta_h$ is not exact.

□

*Theorem 2:* Let cell state be decomposed as $Z = (Z_{\parallel}, Z_{\perp})$with realization $z = (\zeta,\ \xi) \in \mathbb{R}^D$ where $\zeta \in \mathbb{R}^{d_B}$ contains the developmental modes and $\xi$ contains the remaining of transverse modes. Assume that, in a neighborhood of a nondegenerate local minimum $z^* = (\zeta^*,\ \xi^*)$, the Waddington potential is locally quadratic,

$$V(z) = V(z^*) + \frac{1}{2}(z - z^*)^T K (z - z^*), \text{ with } K = \nabla^2 V(z^*) = \begin{pmatrix} K_{\zeta\zeta} & K_{\zeta\xi} \\ K_{\xi\zeta} & K_{\xi\xi} \end{pmatrix} \succ 0.$$

Assume further that the local equilibrium density is Boltzmann distributed $p_Z(z) \propto e^{-\beta V(z)}$, where $\beta$ is the inverse temperature, and that at the variational optimum the marginal distribution of the differentiation state matches that of the developmental modes, $p_B = p_{Z_{\parallel}}$. Then $\mathcal{H}(Z|B) = \frac{D - d_B}{2} \log \frac{2\pi e}{\beta} - \frac{1}{2} \log \det K_{\xi\xi} + \mathcal{H}(B|Z)$, where $K_{\xi\xi} = \nabla_{\xi}^2 V(z)$ is the curvature of the Waddington potential along the transverse modes.

*Proof:* under the local quadratic approximation,

$$p(\zeta,\xi) \propto \exp\left( \frac{-\beta}{2} \begin{pmatrix} \zeta - \zeta^* \\ \xi - \xi^* \end{pmatrix} \begin{pmatrix} K_{\zeta\zeta} & K_{\zeta\xi} \\ K_{\xi\zeta} & K_{\xi\xi} \end{pmatrix} \begin{pmatrix} \zeta - \zeta^* \\ \xi - \xi^* \end{pmatrix} \right).$$

Define the displacements from the local minimum as $\delta\zeta = (\zeta - \zeta^*), \delta\xi = (\xi - \xi^*)$. Using the block decomposition of $K$, the quadratic potential can be expanded as

$V(z) = V_* + \frac{1}{2}\delta\zeta^T K_{\zeta\zeta}\delta\zeta + \frac{1}{2}\delta\xi^T K_{\xi\xi}\delta\xi + \delta\zeta^T K_{\zeta\xi}\delta\xi$. Since $K \succ 0$, its principal block $K_{\xi\xi}$ is also positive definite and therefore invertible. Completing the square with respect to $\delta\xi$ gives

$$p(\zeta, \xi) \propto \exp\left(\frac{-\beta}{2}\left(\delta\xi_{\mathrm{eff}}^T K_{\xi\xi} \delta\xi_{\mathrm{eff}} + \delta\zeta^T K_{\mathrm{eff}} \delta\zeta\right)\right),$$

where $K_{\mathrm{eff}} = K_{\zeta\zeta} - K_{\zeta\xi} K_{\xi\xi}^{-1} K_{\xi\zeta}$ and $\delta\xi_{\mathrm{eff}} = \delta\xi + K_{\xi\xi}^{-1} K_{\xi\zeta} \delta\zeta$.

This yields $p(\xi|\zeta) = \mathcal{N}\left(\xi^* - K_{\xi\xi}^{-1} K_{\xi\zeta} \delta\zeta, \frac{1}{\beta} K_{\xi\xi}^{-1}\right)$. The cross-curvature $K_{\xi\zeta}$ therefore changes the conditional mean of the transverse modes but not their conditional covariance. Because $\xi$ has dimension $D - d_B$, the entropy of this Gaussian is $\mathcal{H}(Z_\perp|Z_\parallel) = \frac{D-d_B}{2} \log\left(\frac{2\pi e}{\beta}\right) - \frac{1}{2} \log\det K_{\xi\xi}$.

We next relate this transverse entropy to the conditional entropy of the full cell state. By the entropy chain rule, $\mathcal{H}(Z|B) = \mathcal{H}(Z) - \mathcal{H}(B) + \mathcal{H}(B|Z)$. Furthermore $\mathcal{H}(Z) = \mathcal{H}(Z_\parallel) + \mathcal{H}(Z_\perp|Z_\parallel)$.

By assumption $p_B = p_{Z_\parallel}$, hence $\mathcal{H}(B) = \mathcal{H}(Z_\parallel)$. Therefore, the developmental marginal entropies cancel, leaving $\mathcal{H}(Z|B) = \mathcal{H}(Z_\perp|Z_\parallel) + \mathcal{H}(B|Z)$. Substituting the expression for the transverse Gaussian entropy yields $\mathcal{H}(Z|B) = \frac{D-d_B}{2} \log\frac{2\pi e}{\beta} - \frac{1}{2} \log\det K_{\xi\xi} + \mathcal{H}(B|Z)$.

□

### Model Architecture

**Potential Network:** The Waddington potential $V(z)$ is parameterized by a fully connected MLP with $ln\ cosh$ activations. Since $ln(cosh(x)) \geq 0$ for all $x$, this architecture guarantees a non-negative potential without requiring additional constraints or penalties. The output layer produces a scalar potential value for each latent state. To stabilize training, we apply an $L_\infty$penalty on the potential output, controlling the maximal excursion of the landscape while retaining flexibility in its shape.

**Other Networks:** All remaining components of scLSD, including encoders and decoders, are parameterized by two-layer MLPs with Softplus activation functions by default. These choices ensure smoothness and positivity where required (e.g., for scale parameters). The scLSD framework, however, is fully modular and allows users to specify alternative network architectures as needed

**Training Procedure:** All components of the architecture—including encoder, decoder, state encoder, and potential network—are trained jointly using the Adam optimizer with a CosineAnnealingWarmRestarts scheduler. The default parameters are:

- **Learning rate:** $2 \times 10^{-3}$ (decayed to $10^{-5}$minimum)
- **Warm restart period:** $T_0 = 30$
- **Multiplier:** $T_{mult} = 1$
- **Gradient clipping:** norm capped at 10

#### Hyperparameter Selection and Ablation Analysis

In addition to architectural choices, scLSD depends on several key hyperparameters that control regularization strength, stochastic sampling, and training dynamics. These include the potential regularization coefficient, learning rate, the KL annealing coefficient, and parameters governing random-walk generation in latent space (number and length of walks). We performed a systematic ablation study using grid search across these hyperparameters to assess robustness and sensitivity.

The potential regularization coefficient controls the amplitude of the inferred Waddington landscape. We evaluated values in the range $\{10^{-4}, 10^{-3}, 10^{-2}\}$ balancing landscape smoothness with expressive flexibility.

For KL annealing, unlike conventional settings that use annealing factors smaller than one, we explored values between 1 and 3. This choice reflects the structure of our model, in which temporal priors—introduced via the RNN—already impose a time-dependent inductive bias on latent representations.

The learning rate was selected from $\{10^{-4}, 10^{-3}, 10^{-2}\}$ covering a range from conservative to aggressive updates, and was included in the hyperparameter ablation to assess training stability and convergence.

The number of random walks is primarily determined by dataset size. For smaller datasets such as dentate gyrus development and zebrafish axial mesoderm development, we used 1024–2048 walks. For larger datasets, including lung cancer progression and pancreas endocrinogenesis, we evaluated values between 4096 and 16384 walks.

The most sensitive hyperparameter in practice is the random-walk length, which must be chosen carefully. Walks that are too short may fail to encounter fully differentiated, high-pseudotime states, while overly long walks can spend most steps in terminal regions, reducing informative transitions. We therefore explored walk lengths ranging from 4 to 64 across datasets. In practice, most datasets performed optimally with walk lengths between 8 and 32, though the optimal value should be determined on a per-dataset basis.

To evaluate hyperparameter robustness, we used multiple complementary metrics. For the inferred Waddington potential and flow field, we applied the CBDir metric to assess the consistency of induced directional dynamics. To evaluate reproducibility of large-scale gene perturbation results, we perturbed only the top 50 scLSD-ranked genes across different models, as exhaustive perturbation is computationally prohibitive. Because the magnitude of fate log-fold change varies across models, we focused on sign consistency: genes with negative notochord fate logFC in one model should exhibit the same directional effect in others. We quantified this using bootstrap resampling (n=1000) to estimate the empirical probability of sign conservation across models.

For the lung cancer progression dataset, we assessed reproducibility by computing the Pearson correlation between entropy, pseudotime, and effective plasticity across models.

Component ablations separately removed the nonnegative potential output constraint, varied the constant Wasserstein coefficient, or replaced the recurrent differentiation-state prior mean with zero, retaining the remaining dataset-specific settings. The Wasserstein coefficient was evaluated at 0, $10^{-4}$, $10^{-3}$, $10^{-2}$, 0.1, 0.5, 0.9 and 1. Encoder–ODE consistency was summarized by the area under the curve of variance-normalized squared discrepancies between propagated states and states reconstructed by projection and re-encoding. These comparisons used three seeds per condition. The recurrent-prior comparison used 15 seeds per condition on bone marrow, dentate gyrus and erythroid. One-sided Mann–Whitney tests evaluated the prespecified directional comparisons; their reported P values were nominal.

Separately, ten models were trained with fixed configurations and different random seeds for bone marrow and erythroid. Each of the 45 seed pairs was compared using cell-wise cosine similarity of fate-probability vectors and externally projected velocities, and Spearman correlation of potentials. Fate probabilities used 40 posterior initial-state samples per cell. Seed-pair matrices summarize agreement across cells, whereas cluster-level boxplots summarize each cell's median agreement across seed pairs. These comparisons assess reproducibility of downstream quantities without aligning latent coordinates. Seed pairs share fitted models and should not be treated as 45 independent replicates. Velocity agreement across seeds is distinct from the variance of CBDir across seeds.

#### Posterior fate uncertainty

Cell-level fate probabilities were estimated by sampling initial states from $q(z|x)$, propagating each sample through the deterministic flow and assigning its endpoint to the nearest observed reference state. Endpoint assignment included all annotated clusters, so intermediate outcomes were retained. Discrete fate entropy was calculated as $-\Sigma_c\, p_c \log p_c$ over the full cluster probability vector.

For the principal bone marrow, dentate gyrus and zebrafish axial-mesoderm analysis, the documented protocol uses 50 initial-state samples per cell and 20 model-time units. The lineage-probability benchmark and cross-seed analysis use 10 and 40 samples, respectively; these analysis-specific sampling settings should not be conflated. Probabilistic endpoints characterize posterior initial-condition uncertainty and are not calibrated probabilities of diffusion-driven fate switching.

#### Implementation and Training of RNA Velocity Baselines

Every workflow starts by harmonizing scRNA-seq inputs using scVelo-style filtering (20–30 shared counts, 2–5k HVGs) and 30 principal components with 15–30 neighbors, often recomputing UMAP or borrowing an existing embedding to keep comparisons fair, after which each velocity engine applies a model-specific but intentionally lightweight training schedule: Cell2Fate typically runs 10 epochs with 64–128 cell batches, leverages a data-driven n_modules, and anchors the timescale with a Tmax_prior centered at 500 before exporting posterior velocities (sometimes sweeping num_samples and batch_size to stress-test memory), DeepVelo trims to ~10 epochs regardless of CPU or GPU target while keeping a 0.001 learning rate (or the original trainer defaults for 100-epoch legacy runs and single-epoch debugging) and enforces dense velocity tensors prior to graphing, UniTVelo fixes its configuration to unified-time mode with 5k genes and GPU ID 0 so every dataset shares the same $R^2$-adjusted fit recipe, scVelo baselines toggle between stochastic quick passes and dynamical ODE fitting using the exact same neighbor graph to provide apples-to-apples embeddings, and VeloVI either trains 50–100 epochs directly on spliced/unspliced layers or 500 epochs on the Ms/Mu moments (sampling 25 trajectories, rescaling the velocity norm, and persisting both latent time and velocity fields) before rendering standard stream plots across Bone Marrow, Pancreas, Dentate Gyrus, and Mouse Erythroid datasets.

#### Benchmark metric definitions and weighting

For lineage-aware probabilistic accuracy, each source population was assigned a predefined set of allowed endpoint labels. A cell's score was the total predicted probability assigned to that set; probability assigned to other labels, including nonterminal endpoints outside the set, remained in the denominator. The RNA-velocity/pseudotime comparison summarized these scores across eligible cells. The continuous-model comparison instead scored hard endpoint assignments within source lineages and then averaged lineage accuracies equally. The two summaries therefore use different within-dataset weighting and prediction conventions. Cross-dataset means gave each available dataset equal weight.

For the continuous-model comparison, CBDir was averaged within each directed lineage edge and then equally across edges. Trajectory stability was the ratio of mean velocity norm over the final 10% of a trajectory to the mean over its initial 10%, summarized by the median across trajectories. This is a comparison of trajectory segments, not a comparison between cells selected from early and late pseudotime. Initial-cell pseudotime thresholds of 0.1, 0.3 and 0.5 were evaluated separately. Undefined measurements were retained as missing rather than assigned zero.

Distributional agreement was evaluated using one-dimensional Wasserstein distance on reference-assigned pseudotime and an unbiased squared MMD estimator with an RBF kernel. In the inspected evaluation code, the RBF coefficient is the inverse of twice the median positive squared distance among up to 1,024 deterministically sampled reference points; a coefficient of one is used if no positive reference distance is available. An unbiased $MMD^2$ estimate may be slightly negative because of finite-sample estimation. State-distribution comparisons used each model's reference-derived coordinate standardization and therefore do not constitute evaluation in a single common learned geometry.

#### Lineage Graphs Used for CBDir Evaluation

To quantitatively assess the directional consistency of inferred velocity fields using the CBDir metric, we specified a lineage graph for each dataset based on established biological knowledge of differentiation hierarchies. These graphs define directed edges between annotated cell clusters, representing known parent–child relationships along developmental trajectories. The CBDir score is then computed by evaluating whether inferred velocities align with these predefined lineage directions.

The lineage graphs used for each dataset are as follows:

**Dentate Gyrus Development**

Directed edges capture the progression from neural progenitors to mature granule neurons and astrocytes:

nIPC → Neuroblast , Radial Glia-like, Neuroblast → Granule immature

Granule immature → Granule mature, Radial Glia-like → Astrocytes

**Erythroid Gastrulation**

Edges represent successive stages of erythroid maturation:

Blood progenitors 1 → Blood progenitors 2, Blood progenitors 2 → Erythroid1

Erythroid 1 → Erythroid 2, Erythroid 2 → Erythroid 3

**Bone Marrow Hematopoiesis**

The graph encodes branching from hematopoietic stem cells toward erythroid, myeloid, and dendritic lineages:

HSC 1 → HSC 2, Ery 1, Ery 1 → Ery 2, HSC 2 → Precursors

Precursors → DCs, Mono 1, Mono 2

**Pancreatic Endocrinogenesis**

Edges describe the transition from ductal progenitors through endocrine commitment to terminal hormone-producing cell types:

Prlf. Ductal → Ductal, Ductal → Ngn3 low, Ngn3 low → Ngn3 high

Ngn3 high → Fev+, Epsilon, Fev+ → Fev+ Alpha, Fev+ Beta

Fev+ Alpha → Alpha, Fev+ Beta → Beta, Fev+ Delta → Delta

These lineage definitions provide a biologically grounded reference for evaluating whether inferred differentiation dynamics are directionally consistent with known developmental programs.

#### Computational cost and scalability

We evaluated the computational cost of scLSD using end-to-end training benchmarks on the Bone Marrow and Cancer datasets. The benchmark scripts reproduced the model construction, prior-transition construction, random-walk generation, and training settings used in the corresponding manuscript notebooks. Plotting, downstream postprocessing, checkpoint loading or saving, and writing of model artifacts were excluded so that the measurements reflect the computational requirements of trajectory construction and model training.

Benchmarks were performed on a server equipped with an NVIDIA H100 80 GB HBM3 GPU and an Intel Xeon Platinum 8468 CPU, with 32 CPU threads available to each process. The software environment consisted of Python 3.10.12, PyTorch 2.4.1 with CUDA 12.1, and sclsd version 0.3.0. Hardware and software information was detected and recorded automatically at runtime to facilitate reproduction on other systems.

The Bone Marrow benchmark contained 5,292 cells and 2,000 genes. Training used 4,096 synthetic trajectories of length 10, a batch size of 256, and 200 epochs. End-to-end execution required 1,596.7 s (approximately 26.6 min), of which 1,596.1 s corresponded to model training. Peak CPU resident memory was 2.89 GiB, peak GPU process memory was 1.67 GiB, and the maximum PyTorch GPU allocation was 0.78 GiB. Mean GPU utilization during training was approximately 37%.

To assess performance on a substantially larger dataset, we benchmarked the lung Cancer dataset containing 33,179 cells and 1,999 genes. This configuration used 8,192 synthetic trajectories of length 8, a batch size of 256, and 100 training epochs. End-to-end execution required 698.3 s (approximately 11.6 min), with 696.9 s spent in model training. Peak CPU resident memory was 6.62 GiB, peak GPU process memory was 1.39 GiB, and peak PyTorch allocation was 0.54 GiB. Mean GPU utilization during training was approximately 34%. The shorter runtime of this dataset despite its larger cell population reflects differences in the number of training epochs, trajectory configuration, and model settings; these two runs should therefore not be interpreted as a direct empirical estimate of runtime scaling with cell number.

Construction of the prior-transition structure and generation of synthetic trajectories represented only a small fraction of the total runtime. For Bone Marrow, prior-transition construction required approximately 0.006 s and random-walk generation approximately 0.44 s. For the 33,179-cell Cancer dataset, these stages required approximately 0.03 s and 0.65 s, respectively. Random-walk generation used approximately 10 MiB of additional PyTorch allocation and was performed on CPU, leaving GPU memory available for model optimization.

The current trajectory-generation procedure operates on the nonzero edges of the local neighbor graph rather than requiring storage of a complete cell-by-cell transition structure. Consequently, the memory requirement for transition construction and sampling scales with the number of nonzero graph edges, $O(nnz)$. For a k-nearest-neighbor graph with fixed k, this is approximately linear in the number of cells, $O(kN)$. The 33,179-cell benchmark demonstrates that the complete scLSD workflow can therefore be trained with modest GPU memory requirements on datasets containing tens of thousands of cells.

These measurements represent individual end-to-end runs rather than timing distributions, and computational requirements will depend on the number of cells and genes, latent-state dimensions, number and length of synthetic trajectories, batch size, and number of training epochs. In the configurations examined here, model optimization dominated total runtime, whereas construction and sampling of synthetic developmental trajectories contributed negligibly to overall wall time.


1. Shiraishi, N. An introduction to stochastic thermodynamics. *Fundamental Theories of Physics. Springer, Singapore* (2023).
2. Kuehn, C. Moment closure—a brief review. *Control of self-organizing nonlinear systems*, 253-271 (2016).
